# Canalization of genome-wide transcriptional activity in *Arabidopsis thaliana* accessions by MET1-dependent CG methylation

**DOI:** 10.1101/2022.07.14.500095

**Authors:** Thanvi Srikant, Wei Yuan, Kenneth Wayne Berendzen, Adrián Contreras Garrido, Hajk-Georg Drost, Rebecca Schwab, Detlef Weigel

## Abstract

**BACKGROUND:** Eukaryotes employ epigenetic marks such as DNA methylation at cytosines both for gene regulation and genome defense. In *Arabidopsis thaliana*, a central role is played by methylation in the CG context, with profound effects on gene expression and transposable element (TE) silencing. Nevertheless, despite its conserved role, genome-wide CG methylation differs substantially between wild *A. thaliana* accessions.

**RESULTS:** We hypothesized that global reduction of CG methylation would reduce epigenomic, transcriptomic and phenotypic diversity in *A. thaliana* accessions. To test our hypothesis, we knocked out *MET1*, which is required for CG methylation, in 18 early-flowering *A. thaliana* accessions. Homozygous *met1* mutants in all accessions suffered from a range of common developmental defects such as dwarfism and delayed flowering, in addition to accession-specific abnormalities in rosette leaf architecture, silique morphology and fertility. Integrated analysis of genome-wide methylation, chromatin accessibility and transcriptomes confirmed that inactivation of *MET1* greatly reduces CG methylation and alters chromatin accessibility at thousands of loci. While the effects on TE activation were similarly drastic in all accessions, the quantitative effects on non-TE genes varied greatly. The expression profiles of accessions became considerably more divergent from each other after genome-wide removal of CG methylation, although the expression of genes with diverse expression profiles across wild-type accessions tended to become more similar in mutants.

**CONCLUSIONS:** Our systematic analysis of MET1 requirement for genome function in different *A. thaliana* accessions revealed a dual role for CG methylation: for many genes, CG methylation appears to canalize expression levels, with methylation masking regulatory divergence. However, for a smaller subset of genes, CG methylation increases expression diversity beyond genetically encoded differences.

## BACKGROUND

Eukaryotic gene expression can be fine-tuned by epigenetic changes such as modifications to DNA, histones, and changes in chromatin architecture. DNA methylation is established and maintained by a cohort of methyltransferases, including MET1/DNMT1 (DNA METHYLTRANSFERASE 1), which semi-conservatively copies methylation marks in the CG nucleotide context from the template to the daughter strand. In plants, MET1 is the principal enzyme for establishing cytosine methylation in replicating cells, especially during fertilization and embryogenesis [1, 2].

In *A. thaliana*, even partial inactivation of MET1 can profoundly alter the genome-wide distribution of cytosine methylation, often causing phenotypic abnormalities due to the emergence of epialleles affecting the activity of developmental genes [3, 4]. These effects are aggravated when *MET1* activity is reduced further, as in the EMS-induced *met1-*1 mutant and the T-DNA insertion mutant *met1-*3, in which genome-wide CG methylation is largely eliminated, particularly at pericentromeric heterochromatin [5, 6]. This in turn affects histone methylation [7–9], chromatin accessibility and long-range chromatin interactions [10], and also leads to ectopic methylation by *de novo* cytosine methylation pathways [11, 12].

The genomes of natural accessions of *A. thaliana* vary considerably, with an average of one single nucleotide polymorphism (SNP) every 200 base pairs of the genome in a given pairwise comparison of accessions from different parts of the geographical range [13]. Natural accessions also vary substantially in their methylome, transcriptome, and mobilome (transposable element, TE) landscapes [14–17]. Large-scale structural variation along with methylome variation at TEs is influenced by genetic variation at loci encoding components of the methylation machinery, suggesting that the methylation machinery is a target of selection during adaptation to the environment [15,16,18–21]. Substantial variation in methylation is also apparent in genic regions, functioning as a storehouse of epialleles, some of which can impact key developmental processes and fitness under new environments [22, 23]. Despite the documented variation in methylome patterns and the known connections between DNA methylation and gene expression, how much variation in DNA methylation contributes to adaptive variation in gene transcriptional activity remains a matter of intense debate [24–27].

Given the large variation in methylomes across *A. thaliana* accessions, we hypothesized that reduction of methylation would reduce differences in chromatin accessibility and gene expression between accessions. To study genetic-background-dependent responses to genome-wide CG hypomethylation, we generated *met1* loss-of-function mutants in 18 *A. thaliana* accessions. The number of differentially expressed genes varied greatly in accessions. While TE-related genes behaved very similarly across accessions, responding nearly uniformly with a substantial increase in expression, non-TE associated genes were much more variable, both in terms of the number of differentially expressed genes and the extent of expression change of individual genes. However, a small group of genes with divergent expression profiles across wildtypes became more similar in expression once MET1 was lost. We conclude that MET1-dependent DNA methylation has dual roles, reducing differences in the transcriptional activity of diverse genomes for most genes, but increasing transcriptional diversity of a minority of genes.

## RESULTS

### Generation of *met1* mutants

We used CRISPR-Cas9 mutagenesis [28] to target the *MET1* gene (*AT5G49160*) in 18 early-flowering accessions of *A. thaliana*, creating frameshift mutations in exon 7 **(Methods, Table S1)**. For each accession, transgene-free lines were genotyped at the *MET1* locus and propagated. In the next generation, we obtained homozygous *met1* mutants from 17 accessions, with two mutant lines of independent origin for 14 accessions and one mutant line for Cvi-0, Ler-1 and Col-0. For Bl-1, we did not recover a sufficient number of homozygous progeny for in-depth analysis. Because Bl-1 heterozygotes already had morphological defects, we included these in our analyses, along with heterozygous mutants in Col-0, Ler-1 and Bu-0. We also included second-generation *met1* homozygotes from Tsu-0 and Tscha-1 (descended from siblings of first-generation homozygotes) to glean first insights into progressive changes at later generations of homozygosity. For one Bs-1 line with biallelic mutations in *MET1,* we only analyzed second-generation progeny that was homozygous for one of the two alleles.

Previous work showed that epigenetic states across the genome can diverge in different lineages of *met1* mutants over several generations [5, 11]. To ensure that we could directly link chromatin state and gene expression, we performed paired BS-seq, ATAC-seq and RNA-seq on leaf tissue of the same plant rosettes, collected as three biological replicates for both wild-type and *met1* mutant lines **(Methods)**. All together, we obtained 73 BS-seq, 158 ATAC-seq and 158 RNA-seq libraries that passed quality control.

### MET1 can both buffer and increase transcriptomic variation across accessions

Since natural accessions of *A. thaliana* are known to express diverse transcriptomes in their wild-type state [16], we hypothesized that MET1-induced CG methylation may contribute to generating this diversity, and that the transcriptomes would become more similar to each other in *met1* mutants. We therefore first grouped all wildtypes (54 samples) and all *met1* mutants (104 samples) separately, and analyzed 19,473 genes with sufficient read counts in both groups. Contrary to our hypothesis, UMAP projections of read counts at these genes revealed greater differences between accessions in *met1* mutants compared to their respective wild-type samples (**Figure 1a**). This suggested an alternate hypothesis where MET1 functions in buffering transcriptomic diversity across accessions, with its absence therefore unmasking larger regulatory differences. Wild-type samples had 1,210 differentially expressed genes (DEGs, defined as genes with |log_2_(fold change)|≥1, FDR adjusted p-value≤0.01) between accessions, fewer than those identified between accessions in *met1* mutant samples (1,868 DEGs). There were only 406 DEGs that were shared by wildtypes and *met1* (**Figure 1d**), i.e., which were differentially expressed independently of *MET1* activity. These are likely to include genes that are associated with structural variation or major differences in cis-regulatory sequences. There were 804 DEGs that were unique to wild-type samples (**Figure 1b**), i.e., where differences in expression between accessions were greatly reduced upon loss of *MET1*. Finally, the largest group in this comparison comprised 1,462 DEGs that were unique to *met1* samples (**Figure 1c**), i.e., which were expressed at similar levels across wildtypes, but became differentially expressed in *met1* mutants. These inferences were also apparent from density distributions (**Figure S1a**) and heatmaps (**Figure 1b-d, Figure S1b**) of Pearson Correlation coefficients between accessions. Together, these observations indicate that there is evidence for both of our alternative hypotheses - MET1 reduces transcriptomic diversity across accessions for many genes, but for a smaller group of genes adds another layer of expression complexity to diversify transcriptomes.

**Figure 1.**
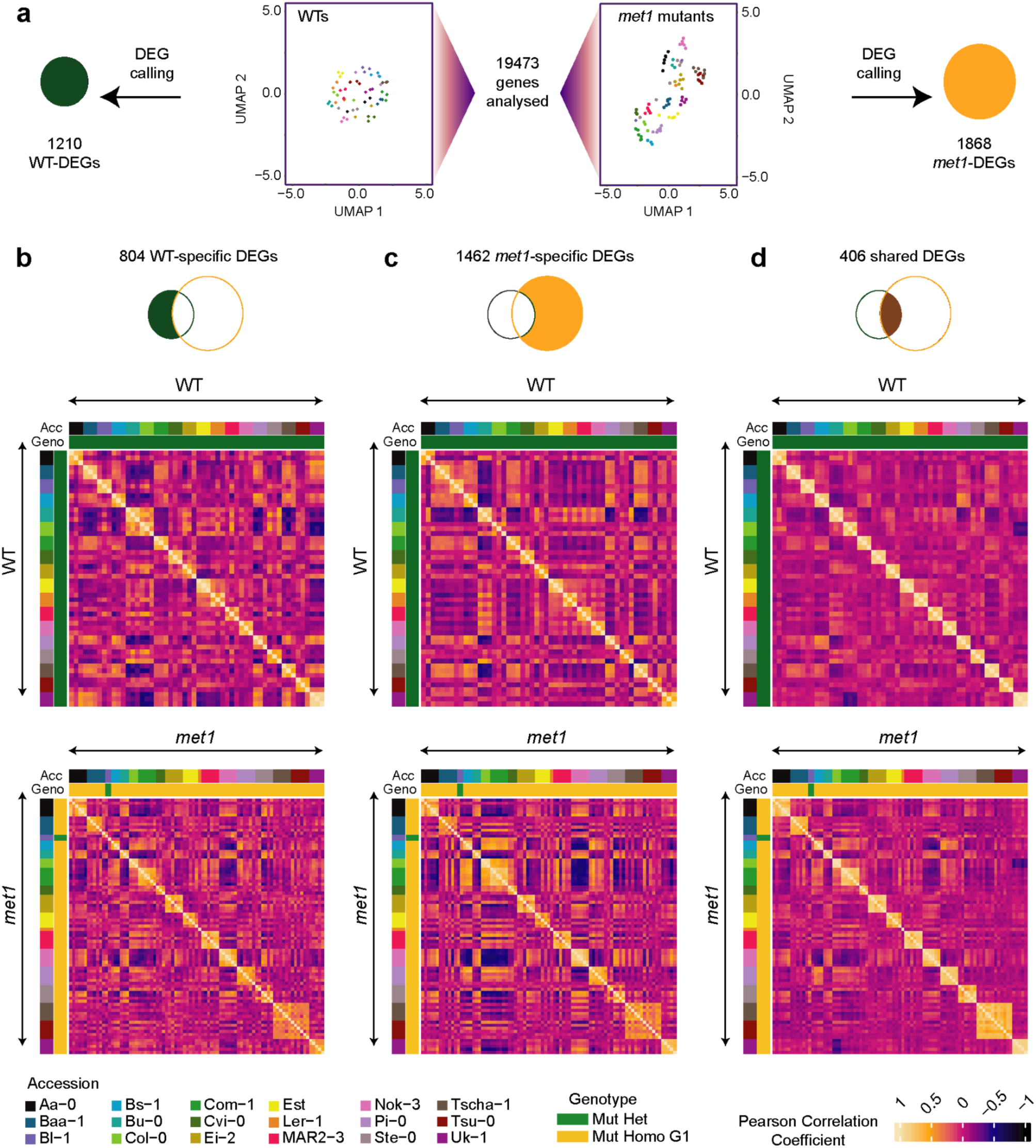
Transcriptomic variation among accessions in *met1* mutants and wildtypes. (a) UMAP projections of read counts in 19,473 genes for wildtypes (left) and *met1* mutants (right). These genes were further analyzed to identify DEGs. (b-d) Pearson correlation heatmaps of gene expression levels between accessions in *met1* mutants and wildtypes, for DEGs specific to wildtypes (b), DEGs specific to *met1* mutants (c) and DEGs shared between *met1* mutants and wildtypes (d).

### TE- and Non-TE genes are differentially expressed in *met1* mutants

A major role of *MET1* is the transcriptional silencing of TEs by establishing and maintaining CG methylation [29, 30]. We next asked whether there were any accession-specific differences in the requirement of *MET1* in regulating the expression of both TE genes (genes associated with TEs, see Methods) and other protein-coding genes. We first analyzed RNA-seq read counts of 21,657 genes in all samples (wildtypes and *met1* mutants) and generated a UMAP visualization. Two distinct clusters of samples could be observed, one encompassing wild-type and the other *met1* plants (**Figure 2a**). In agreement with our initial results, wild-type samples were less spread out than mutant samples, indicating greater gene expression heterogeneity in *met1* mutants than in wild-type plants. Comparison of first- and second-generation homozygous mutants in the Tscha-1 and Tsu-0 accessions did not indicate major changes upon propagation of mutant lines.

**Figure 2.**
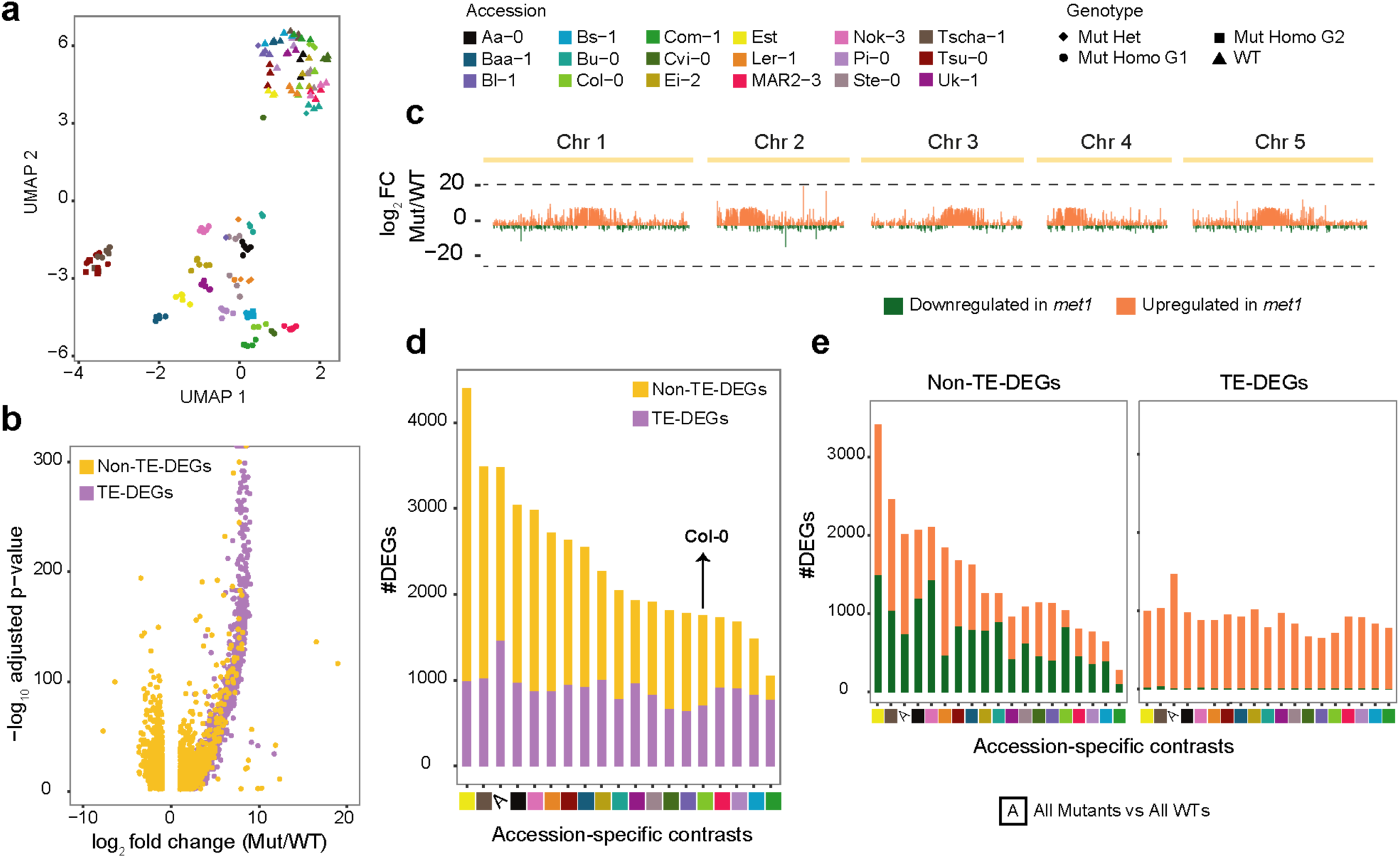
Accession-specific DEGs in *met1* mutants. **(a)** UMAP representation of transformed RNA-seq counts from 158 samples (104 hetero- or homozygous *met1* mutants and 54 wild-type plants) across 21,657 genes. Colors indicate accessions, and shapes indicate genotype (see top right). WT, wild type; Mut Het, heterozygous *met1* mutants; Mut Homo G1, first-generation homozygous *met1* mutants; Mut Homo G2, second-generation homozygous *met1* mutants. **(b)** Volcano plot of 3,479 DEGs identified in a contrast between all *met1* mutant samples and all wild-type samples. TE-associated DEGs (TE-DEGs) are colored purple, and Non-TE-DEGs yellow. **(c)** Chromosomal distribution of 3,479 DEGs from the all-*met1*-against-all-wild-type contrast, and their log_2_(fold change) in mutants relative to the corresponding wildtypes. Upregulated DEGs are colored orange and downregulated DEGs green. **(d)** DEGs in the 18 accession-specific contrasts, compared to the all-*met1*-against-all-wild-type contrast (denoted by ’A’, third column from the left). **(e)** Variation in numbers of upregulated and downregulated Non-TE-DEGs and TE-DEGs across different contrasts.

To obtain first insights into MET1 function across all accessions, we examined protein-coding genes whose expression levels changed significantly in a contrast of all *met1* mutants against all wild-type plants, thereby identifying 3,479 DEGs. Of these, 1,466 genes (42% of all DEGs) were associated with TEs, which we called TE-DEGs, and these corresponded to 87% of the 1,678 TE-associated genes with sufficient information in our dataset **(Methods)**. Almost all of them were upregulated in *met1* mutants compared to wild-type plants, and often greatly so (**Figure 2b**). This was consistent with previous findings from early- and late-generation homozygous *met1* mutants [31–35], including the observation that Class II DNA TEs of the En/Spm superfamily were most often among activated TE-DEGs [36, 37]. Consistent with TE density being highest near the centromeres [38], TE-DEGs were enriched in pericentromeric regions. Many of the TE-DEGs were strongly overexpressed in *met1* mutants, hundreds of times or more **(Figure 2b, 2c)**.

The picture was different for the remaining 2,013 DEGs that were not associated with TEs (Non-TE-DEGs), and which corresponded to 10% of the 19,979 Non-TE genes with sufficient information in our dataset. Although the majority was also upregulated in *met1* mutants, more than a third, 728, was downregulated (**Figure 2b**). Non-TE-DEGs were also more uniformly distributed along the chromosome arms, and overall expression changes for both up- and downregulated genes were much more moderate (**Figure 2c**). Gene Ontology (GO) enrichment of Non-TE-DEGs revealed several terms related to abiotic and biotic stresses and stimulus response **(Figure S2)**. We conclude that across all accessions, Non-TE-associated genes vary much more in their sensitivity to loss of MET1 than TE-associated genes.

### Mis-regulated genes vary among *met1* mutants of different accessions

A closer examination of DEGs called by contrasting all *met1* mutants against all wildtypes showed that DEGs were not uniformly induced or repressed across accessions (a random subset of such DEGs is shown in **Figure S3**). Hence we individually examined each of the 18 accessions for genes that were differentially expressed between *met1* mutants and the corresponding wild-type parents. We found that the number of DEGs in each accession varied substantially (**Figure 1d**), but that this was much more true for Non-TE-DEGs than TE-DEGs. In addition, while down-regulated TE-DEGs were rare in all accessions, the ratio of up- to down-regulated Non-TE-DEGs in *met1* mutants was much more variable **(Figure 1e, Figure S4).** The number of Non-TE-DEGs ranged from 278 (26% of all DEGs) in Com-1 to 3,409 (77% of all DEGs) in Est. Notably, even though we could only analyze heterozygous Bl-1 *met1* mutants, these had more Non-TE-DEGs than homozygous *met1* mutants in several other accessions, suggesting that even the removal of only one of the two functional *MET1* copies was sufficient to alter expression levels of a large number of genes, at least in this accession.

In total, there were 10,151 Non-TE-DEGs that were differentially expressed in at least one of the accessions or in the all-*met1*-against-all-wild-type contrast, which was approximately half of all Non-TE genes that we could assay. We used this set of Non-TE-DEGs to build a weighted gene co-expression network **(Methods)**. The 10,151 Non-TE-DEGs clustered into nine gene modules. Genes from module ’D’ were the most consistent in their expression levels across all accessions, being on average always up-regulated in *met1* mutants compared to the respective wild-type plants **(Figure S5)**. GO enrichment analyses revealed that many ’D module’ genes were associated with nucleic acid metabolic processes, DNA repair and transcription, although only weakly significantly so. This suggests that MET1 likely affects core metabolic machinery genes, which may further impact a different subset of downstream genes in each accession.

We next generated a frequency spectrum to examine the overlap of DEGs between accessions. When we focused on the two extreme accessions, Est and Com-1, a great majority of TE-DEGs were shared across most contrasts, while the picture for Non-TE-DEGs was very different. While almost 30% of the 3,409 Non-TE-DEGs in Est were not detected in any other accession, 98% of the 278 Com-1 Non-TE-DEGs were found in at least one other accession **(Figure 3a-b, Figure S6)**. The 983 unique Non-TE-DEGs in Est were enriched for functions with a common theme of RNA and DNA metabolism (**Figure 3c**), compatible with a scenario in which mis-regulation of one or a few specific master regulators had led to expression changes in numerous Non-TE-DEGs in Est.

**Figure 3.**
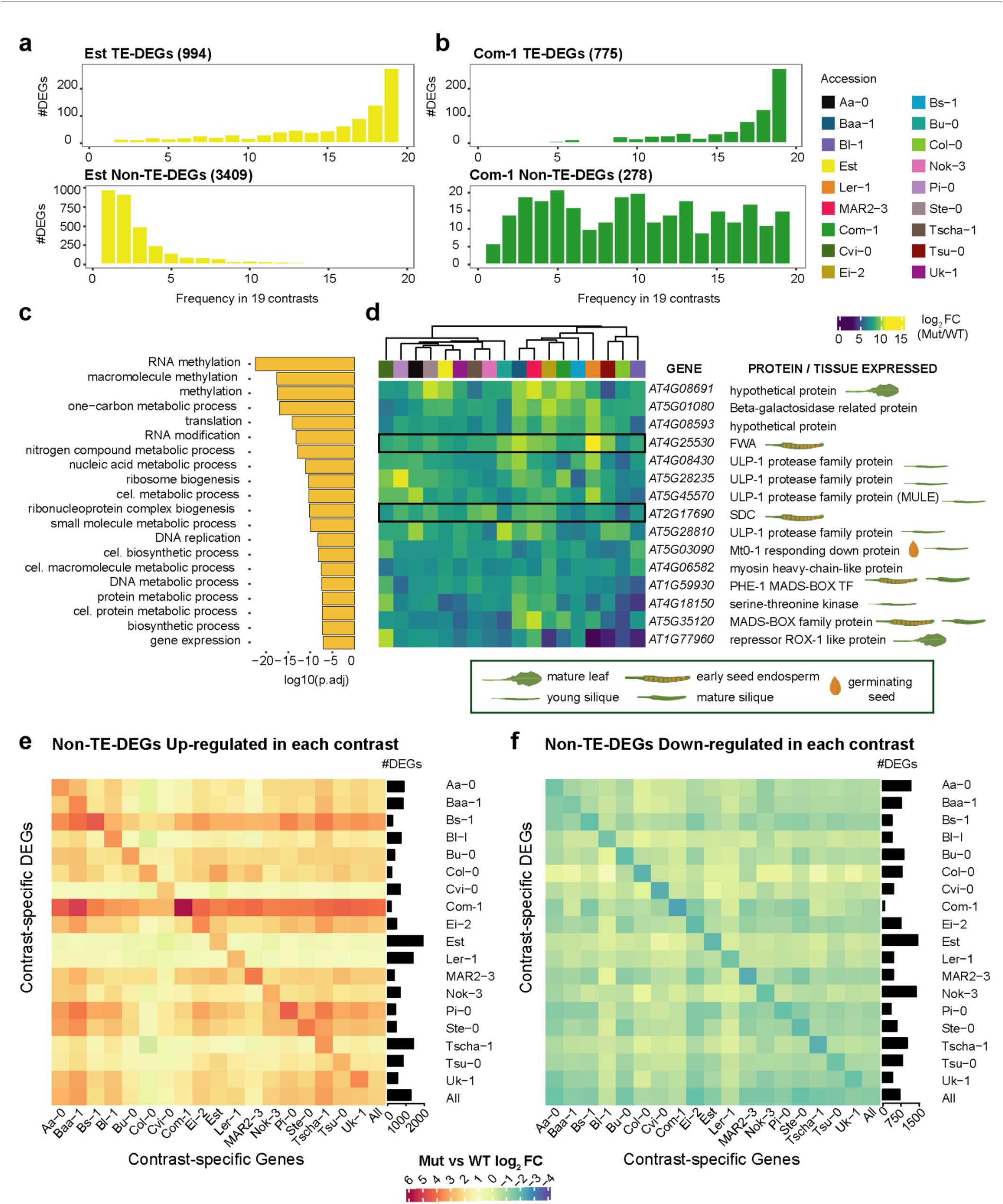
Qualitative and quantitative comparisons of accession-specific DEGs. **(a, b)** Frequency spectrum of Est and Com-1 TE-DEGs and Non-TE-DEGs across all other accessions**. (c)** The top 20 GO terms enriched for 983 DEGs unique to Est. **(d)** Heatmap of log_2_(fold change) of 15 universal Non-TE-DEGs across all accessions; *FWA* and *SDC* are highlighted with a black outline. TAIR10 gene names, encoded proteins and their preferential tissues of expression in wild type (where known) are shown. (e, f) Heatmaps of average log_2_(fold change) for all Non-TE-DEGs from one accession in each of the other accessions. Barplots on the right indicate the absolute frequency of Non-TE-DEGs in each accession.

When overlaying DEGs from the 18 accession-specific *met1*-against-wild-type comparisons and the all-*met1*-against-all-wild-type comparison, we found 291 universal DEGs, albeit with the extent of expression change in *met1* mutants differing considerably across accessions. Only 15 of the universal DEGs were Non-TE-DEGs (**Figure 3d**), and eight of these 15 genes are strongly expressed in siliques in Col-0 wild-type plants [39]. We had measured gene expression in rosette leaves, and many of these genes were expressed at low levels in our wild-type samples, but strongly upregulated in *met1* mutants. Included among the 15 genes were three maternally imprinted genes, *FWA, SDC* and *AT1G59930* [40, 41]. Ectopic *FWA* expression is known to delay flowering in both Col-0 and Ler-1 accessions [42, 43], while ectopic activation of *SDC* has been shown to lead to dwarfism and leaf curling in Col-0 [41, 44]. This is consistent with the whole-plant phenotypes described in detail below.

Finally, we assessed to what extent DEGs from one accession changed in the other accessions. Genes identified as up- or down-regulated Non-TE-DEGs after loss of MET1 in one accession changed on average almost always in the same direction in each of the other accessions, but did so less strongly (**Figure 3e**). TE-DEGs on the other hand, were similarly upregulated in all accessions, but down-regulated TE-DEGs were often variably expressed in other accessions (**Figure S7**). This finding is consistent with the idea that MET1 activity often homogenizes gene expression levels in different genetic backgrounds.

### *met1* mutants exhibit reduced DNA methylation and increased overall chromatin accessibility

To investigate how much of the variation in DEGs across accessions arose from differences in DNA methylation and chromatin architecture, we characterized the methylomes and chromatin accessibility in our collection of *met1* mutants and corresponding wild-type lines. As expected from the knockout of *MET1*, cytosine methylation in the CG sequence context was drastically and consistently reduced in homozygous *met1* mutants, dropping to a mean genome-wide level of 0.2%, representing a 98.5% decrease from mean wild-type levels **(Figure S8).** Cytosine methylation in non-CG contexts was moderately affected in first-generation *met1* mutants, being on average 6.8% higher in the CHG context and 21% lower in the CHH context. Second-generation *met1* mutants of Tsu-0 and Tscha-1 had greatly increased CHG methylation, both relative to first-generation *met1* mutants and wild-type parental lines **(Figure S8).** This increase in methylation during successive rounds of inbreeding has previously been described for *met1-*3 mutants in the Col-0 accession [11].

Because overall methylation patterns were greatly altered in *met1* mutants, we wanted to closely examine genomic regions with the most significant changes in methylation. We contrasted 73 methylomes including both *met1* mutants and the wild-type parents from all 18 accessions **(Methods)** to identify differentially methylated regions (DMRs) in the CG, CHG and CHH contexts. The 2,388 CG-DMRs overlapped with the vast majority of the 350 CHG-DMRs and 1,023 CHH-DMRs. Approximately half of all CG-DMRs overlapped with TE sequences, a quarter overlapped TE genes, while 42% overlapped Non-TE genes **(Figure S9, Table S2).**

While most CG-DMRs in *met1* mutants retained minimal or no CG methylation, the extent of reduction in methylation differed across accessions, consistent with accession-specific methylation patterns in the presence of MET1 [16] (**Figure 4a-b, Figure S10**). To assess the functional consequences of this variation, if any, we asked how loss of cytosine methylation impacted genome-wide chromatin architecture in each accession. To this end, we identified accessible chromatin regions (ACRs) by ATAC-seq in all *met1* mutants and the corresponding wildtypes. This analysis allowed us to define a union set of 34,993 ACRs **(Methods),** from which we retained all with an accessibility change in at least two accessions **(Methods)**. This resulted in 31,295 regions that we refer to as differential ACRs (dACRs) **(Figure S11)**. We visualized variation in accessibility levels at these dACRs using UMAP (**Methods**), which similarly to the RNA-seq data (**Figure 2a**) revealed two distinct clusters of wild-type plants and *met1* mutants (**Figure 4c**), with wild-type plants from different accessions being more similar to each other than *met1* mutants from different accessions. Approximately two-thirds of all dACRs positionally overlapped with Non-TE genes, 8% with TE genes and 30% with TE sequences **(Figure S11, Table S2)**.

**Figure 4.**
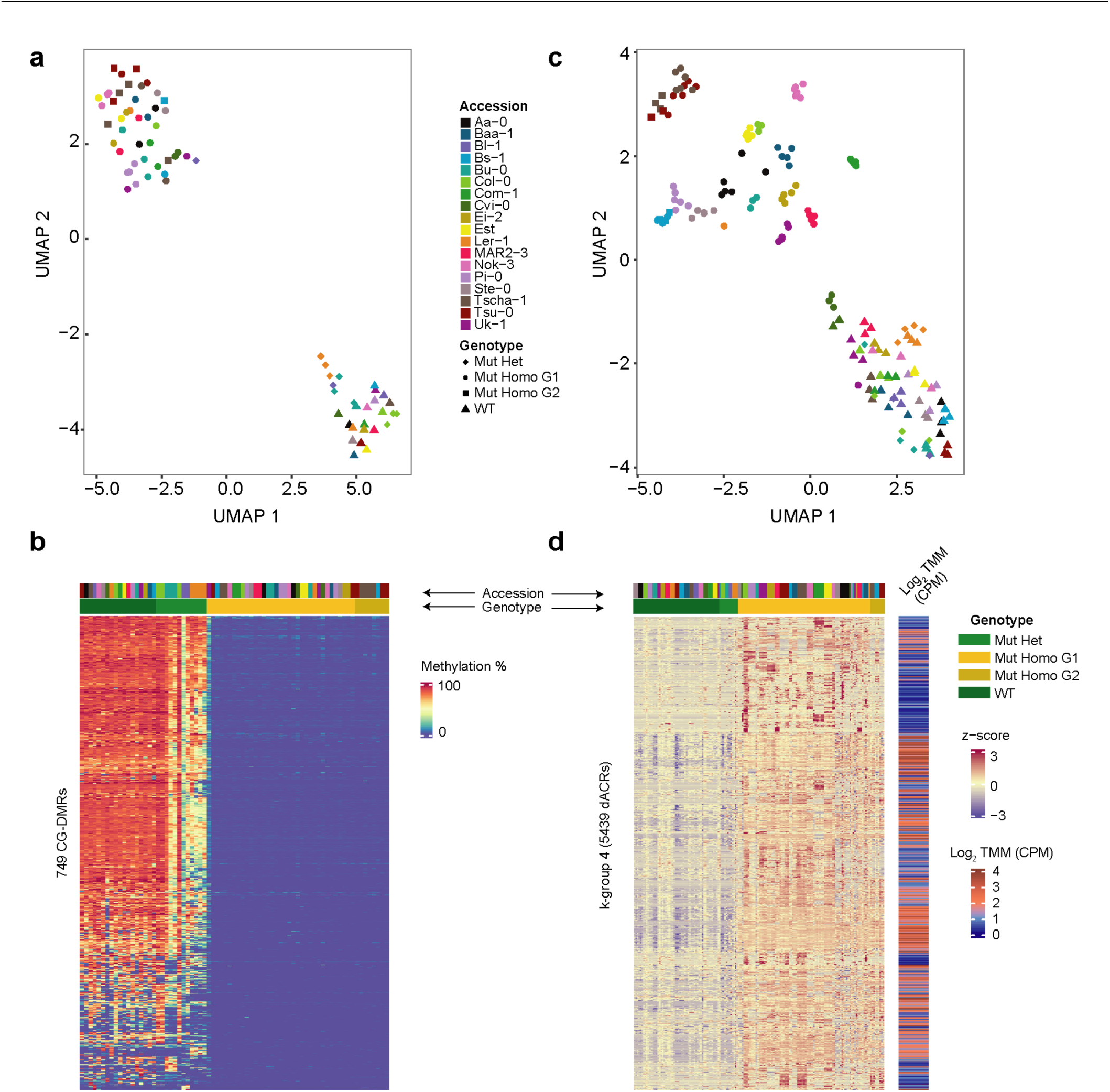
Reduced CG methylation and increased chromatin accessibility in *met1* mutants. **(a, b)** UMAP representation and heatmap of CG methylation levels in wild-type plants and *met1* mutants across 749 CG-DMRs (from a total of 2,388 CG-DMRs). **(c)** UMAP representation of chromatin accessibility in log_2_(CPM) of 31,295 dACRs across wild-type plants and *met1* mutants. **(d)** Heatmap of z-scaled values of 5,439 dACRs with coherent behavior as identified by k-means clustering (k-group 4 in Figure S11), with mean accessibility indicated on the right. TMM, trimmed mean of M-values. CPM, counts per million.

Genome wide, the chromatin of *met1* mutants was more accessible than chromatin of wild-type plants, and this difference was particularly pronounced for a subgroup of 5,439 dACRs, identified by k-means clustering **(Figure 4d, Figure S11). More than** three quarters of the dACRs in this subgroup overlapped with positions of TE sequences, and only 3.5% of these dACRs overlapped with Non-TE genes. Across all accessions, while approximately half (46%) of CG-DMRs overlapped in position with dACRs, most dACRs conversely (95%), did not overlap with DMRs, indicating that the vast majority of dACRs appeared due to *trans* effects of methylation changes in the genome.

To see whether epigenetic profiles at TEs, which included both TE genes and other TE sequences, differed from Non-TE genes, we averaged methylation levels (all contexts) and chromatin accessibility levels for all *met1* mutants and wild-type plants across 31,189 TEs and 29,699 Non-TE genes including 1 kb flanking sequences **(Figure S12)**. Genome-wide chromatin accessibility patterns in *met1* mutants mirrored cytosine methylation levels, with increases in accessibility accompanied by decreases in methylation for most accessions. This pattern was much more pronounced in magnitude over TEs **(Figure S12a)** than in Non-TE genes **(Figure S12b)**. These observations once again demonstrate that TEs are highly sensitive to the absence of MET1, in agreement with TE genes being highly upregulated in *met1* mutants.

Finally, we asked how much of the expression changes at DEGs were explained by these epigenetic alterations. We focused on Est and Com-1, the two accessions with the highest and lowest number of DEGs. While TE-DEGs behaved similarly in the two accessions, Non-TE-DEGs showed contrasting patterns, with accessibility differences between *met1* mutants and wildtypes being on average much smaller in Est than in Com-1 **(Figure S12c)**, confirming a complex relationship between methylation, chromatin accessibility and gene expression changes at Non-TE-DEGs, which in turn leads to very different numbers of Non-TE-DEGs in different accessions.

### Non-TE-DEGs show varying methylation and chromatin accessibility profiles across accessions

Observing that differential gene expression can be accompanied by epigenetic changes, we quantitatively assessed mutual relationships of cytosine methylation and chromatin accessibility with gene expression, having analyzed all of the three factors from the same individuals across our collection of *met1* mutants and wildtypes.

To investigate whether the presence or absence of methylation within- or in proximity to a gene could influence its expression level, we examined how DEGs were regulated in the presence of a DMR. We first defined a consensus set of 7,132 DEGs from all 18 accessions (**Methods, Figure S13**), containing TE-DEGs (1,401) and Non-TE-DEGs (5,731) and intersected their genomic coordinates with 1,569 CG-DMRs with sufficient methylation calls across all samples that were identified from all mutant and wild type samples (**Methods**).

While only a small fraction of consensus DEGs were located close to a CG-DMR (21% of TE-DEGs and 7% of Non-TE-DEGs), the converse was also true: only a minority of CG-DMRs was found next to DEGs (21% next to TE-DEGs and 28% to Non-TE-DEGs). Local differences in CG methylation are thus neither necessary nor sufficient for differences in gene expression between wild-type and *met1* mutant plants. At the loci where both expression and CG methylation were altered, we examined expression counts and methylation levels in homozygous *met1* mutants and in corresponding wild-type plants from 17 accessions (**Figure S14**). As a control, we randomly sampled Non-DEGs (genes that were not differentially expressed; **Methods**) that positionally overlapped with DMRs.

Most TE-DEGs with CG-DMRs in either their extended gene bodies **(Methods, Figure S14, Figure S15c)** or 1.5 kb up- or downstream sequences (’cis’ regions) **(Methods, Figure S14, Figure S16c)** had lost CG methylation in *met1* mutants. These TE-DEGs were upregulated in all *met1* mutants, albeit to a different extent in each accession. Non-TE-DEGs, which were already observed to be highly variable in their differential expression levels, exhibited a wide gradient of methylation differences between *met1* mutants and the respective wild-type parents (ranging from -100% to +10%). This was observed both when DMRs were located in extended gene bodies **(Figure S15a)** and in 1.5 kb up- or downstream sequences **(Figure S16a, Figure S17c)**. In both cases, one group of Non-TE-DEGs was highly upregulated and had highly reduced methylation in *met1* mutants, suggesting that methylation could have a strong effect on gene regulation at these genes. There was also another group of more moderately up- or down-regulated Non-TE-DEGs with negligible methylation changes in *met1* mutants. On examining the accession-of-origin of each of these Non-TE-DEGs, we found that the same gene in different accessions could be found in either of the groups identified above, indicating considerable epigenetic plasticity across accessions.

Consequently, we asked whether parental methylation and expression state can predict epigenetic and transcriptional response in *met1* mutants. DEGs classified in the top quintiles of wild-type CG methylation levels (**Methods**) were more likely to increase in expression than DEGs from the bottom quintiles, which were similarly likely to be up- or downregulated in *met1* mutants **(Figure 5c, 5g)**. In terms of expression changes, we found that DEGs from the lowest quintile of expression counts in the wild-type parents were the ones that were the most upregulated in *met1* mutants **(Figure 5a, 5e)**. These observations are consistent with high levels of CG methylation in the wild type state serving to silence genes.

**Figure 5.**
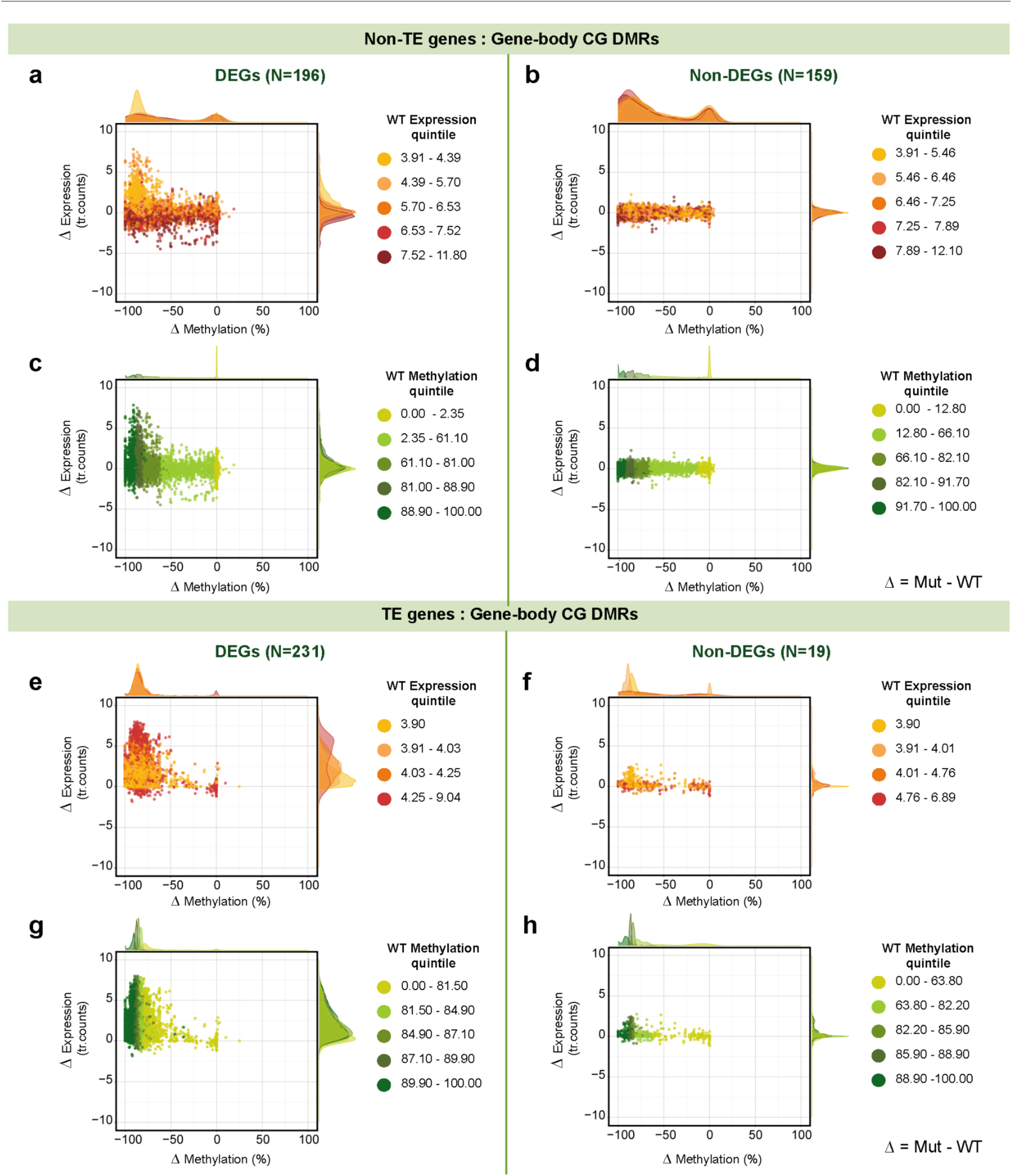
CG-DMRs in gene-bodies of Non-TE-DEGs and TE-DEGs. Differences in CG methylation between *met1* mutants and wild-type plants plotted against differences in gene expression. Dots are colored by wild-type expression quintiles **(a,b,e,f)** and wild-type methylation quintiles **(c,d,g,h)** with density distributions shown on top and left. Expression levels are represented as transformed read counts (tr. counts) and methylation levels as % CG methylation in CG-DMRs.

In *A. thaliana*, many constitutively active genes are marked by gene-body CG methylation (gbM) [45] and the methylation levels of these genes are known to vary in tandem with their differential expression across *A. thaliana* accessions [22, 23]. Most genes with gene body methylation among Non-TE-DEGs (’gbM like’ genes; **Methods)** were downregulated in *met1* mutants **(Figure S18).** Incidentally, genes that were highly methylated and lowly expressed in wildtypes (exhibiting TE methylation characteristics in the CG context or ’teM like’ genes; **Methods**) were always upregulated in *met1* mutants, similarly to TE-DEGs **(Figure S19)**. The two groups of genes, with gene body or TE type methylation in the CG context, did not overlap.

We next carried out a similar quantitative analysis to examine the relationship between differential expression changes and differential chromatin accessibility at DEGs (**Methods**, **Figure S20**). Consistent with the much larger number of dACRs compared to DMRs, over 90% of all consensus DEGs (**Methods**, **Figure S13**) carried dACRs within 1.5 kb upstream and downstream of their gene body. While chromatin accessibility increased almost proportionally with expression in *met1* mutants for TE-DEGs, increased accessibility could be associated with, but did not necessitate an increase in gene expression for Non-TE-DEGs (**Figure S21-S22**). Accessible chromatin is well known to favor gene transcription by facilitating transcription-factor binding [46, 47] although there is also evidence that inaccessible regions can occur in some long and highly transcribed genes [48]. Together, these observations point to complex interactions between gene expression and chromatin accessibility levels in *cis*, especially at Non-TE-DEGs.

### *MET1* can have indirect effects on the expression of Non-TE genes

Since variation in the response of gene expression, methylation and chromatin accessibility to loss of *MET1* was most apparent for Non-TE-DEGs, we focused on these genes to explore the nature of their epigenetic plasticity. Among 5,731 consensus Non-TE-DEGs that varied in their expression response across accessions (**Figure 6a**), a majority (87%) had differentially accessible chromatin regions (dACRs) in their vicinity, but lacked a *cis* CG-DMR (’*cis*’ here including the gene-body). Conversely, a small minority of Non-TE-DEGs, only 12 (0.2%), had *cis* CG-DMRs but no nearby dACRs, and 292 (5%) had neither a nearby dACR nor CG-DMR. Closer inspection of examples from these categories (**Figure 6a**) showed that the relationship between altered gene expression, CG methylation, and accessibility varied substantially in *met1* mutants in different accessions.

**Figure 6.**
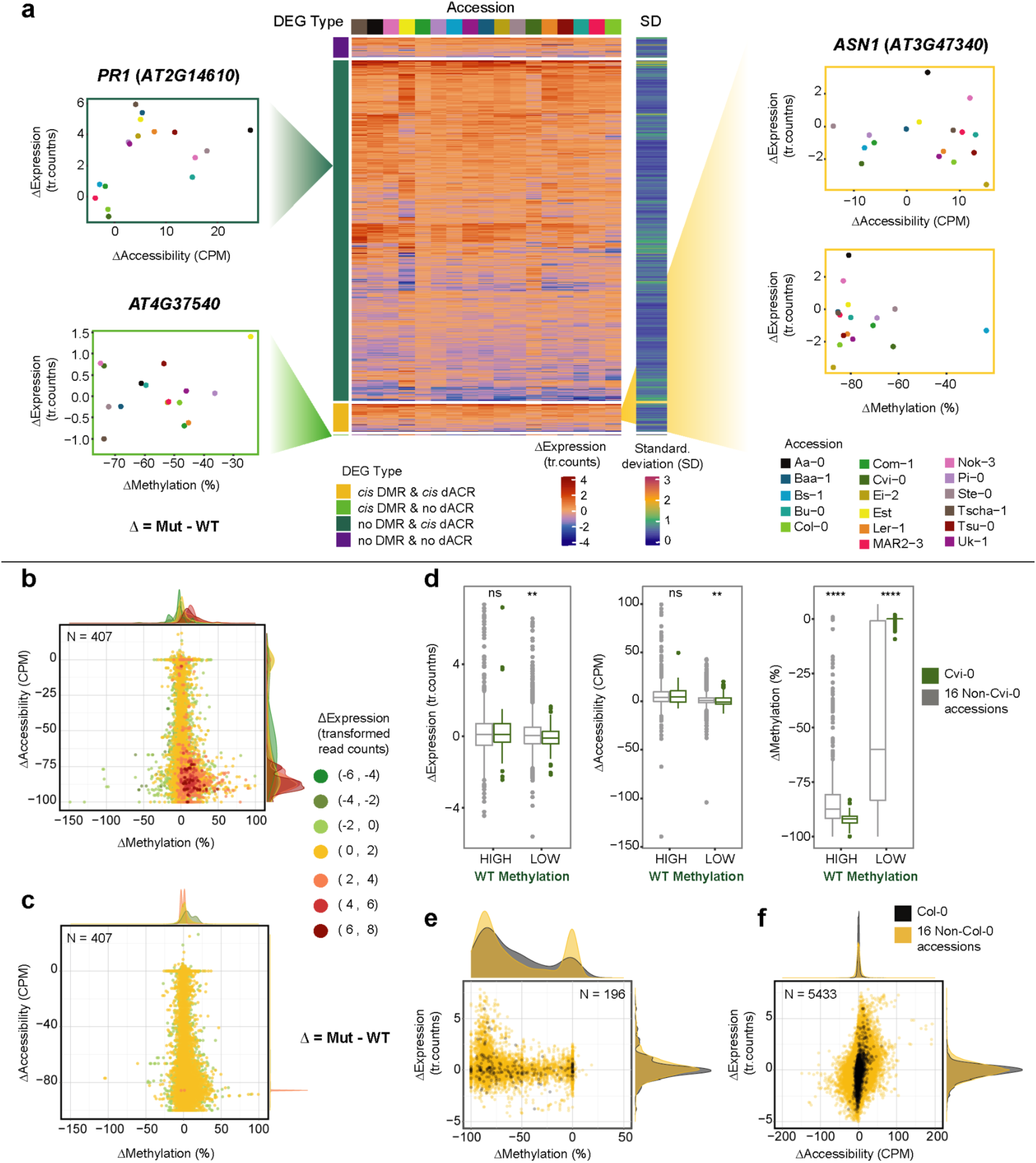
Non-TE-DEGs in *met1* mutants can have different epigenetic states in different accessions. **(a)** Heatmap of expression changes across 5,731 Non-TE-DEGs in 17 accessions, with an adjacent heatmap showing variance expressed as standard deviation (SD) across accessions, and scatterplots of changes in expression and accessibility in representative genes from three different DEG categories (based on overlap with *cis* CG-DMRs and dACRs). **(b)** Scatterplot of changes in chromatin accessibility and methylation in Non-TE-DEGs across 17 accessions. Colors and density distributions represent custom bins of expression changes. **(c)** Scatterplots similar to **(b)** for Non-TE genes. **(d**) Boxplots showing *MET1*-dependent changes in chromatin accessibility, gene expression and CG methylation of genes that are weakly (’LOW’) or highly (’HIGH’) methylated in wild-type Cvi-0. The same genes are compared for Cvi-0 (dark green) and 16 other accessions (gray). **(e)** Scatterplot of changes in methylation and expression in Non-TE-DEGs with gene-body CG-DMRs, colored by DEGs specific to Col-0 (black) against the same genes in other accessions (yellow). **(f)** Scatterplot of changes in chromatin accessibility and expression in Non-TE-DEGs carrying *cis* dACRs, colored by DEGs specific to Col-0 (black) against the same genes in other accessions (yellow). Expression levels are represented as transformed read counts (tr. counts); chromatin accessibility levels as TMM (trimmed mean of M-values) normalized values in counts per million (CPM), and methylation levels as % CG methylation.

To focus on close-range interactions between changes in methylation, chromatin accessibility and gene expression at the same locus, we examined Non-TE genes associated with both *cis* dACRs and CG-DMRs - comprising 407 DEGs, which we compared with 407 randomly sampled Non-DEGs (**Methods**). In general, we found that genes with strongly reduced CG methylation in *met1* mutants had a tendency to show drastic changes in chromatin accessibility, becoming either much more accessible or much less. For both DEGs and Non-DEGs **(Figure 6b, 6c)**, there were many different combinations of methylation and accessibility states, and these did not cluster by degree of expression change. This suggested the presence of multiple epigenetic states, both for different genes in the same accession and for the same gene in different accessions. One group of genes stood out from the rest: this group, which accounted for 10% of all unique DEGs, was upregulated in *met1* mutants, marked by highly reduced methylation that was paralleled by highly increased accessibility, consistent with methylation changes being directly responsible for induction of expression (**Figure 6b**).

We next examined Non-TE genes as a group for accession-specific epigenetic patterns, first comparing the methylation levels of these genes in wildtypes and mutants. Cvi-0 had the highest fraction of genes with minimal reduction in methylation in *met1* mutants **(Figure S23a)**. This was explained by Cvi-0 wild type having already many genes with low methylation levels, limiting the extent of any further reduction in methylation **(Figure S23a)**. We asked how genes with low CG methylation in Cvi-0 wild type, that is, genes with methylation levels that could not be reduced much further by loss of *MET1*, fared in other accessions. While the average reduction in methylation level was greater in other accessions, as expected, changes in accessibility and expression level after inactivation of *MET1* were very similar in magnitude when compared to Cvi-0 (**Figure 6d**). This observation provides further support for genome-wide hypomethylation indirectly affecting the expression of many genes. Finally, comparing the relationship between *MET1-*dependent changes in methylation, chromatin accessibility and gene expression in the reference accession Col-0 with changes in other accessions confirms that Col-0 is not particularly representative of *A. thaliana* accessions at large (**Figure 6e, f**).

### *met1* mutants express signatures of known epialleles

Many of our *met1* mutants had strong methylation and expression changes at well-known loci sensitive to epigenetic regulation, such as *FWA* [42] **(Figure S23c,d)***, SDC* [44] **(Figure S24)**, the *PAI* genes [49], *IBM1* [50], *SNC1* (with alleles similar to the *bal* variant of *SNC1*) [51], the *ROS1* demethylase, which is known to function as a methylation sensor [11,52,53] **(Figure S23b),** *AG* [4] **(Figure S25),** and *SUP* (with alleles similar to the *clark-kent* variant of *SUP*) [54] **(Figure S26)**. New epialleles at several of these loci have been reported before in Col-0 and C24 *met1* mutants [3, 55]. We observed a variety of methylation patterns at these loci depending on the accession of origin, with considerable differences in chromatin accessibility and gene expression across accessions; several examples at the *FWA* locus are shown in **Figure S23c,d**. As seen before [11], some epialleles arose only in second-generation *met1* mutants **(Figure S25)**, suggesting that epialleles continue to accumulate in the absence of *MET1* during inbreeding.

### *met1* mutants vary in phenotypes and segregation distortion in natural accessions

Compared to their wild-type parents, homozygous *met1* mutants were dwarfed, flowered late and had altered rosette leaf architecture - although to different degrees in each accession **(Figure 7a, Figure S27-S28)**. *met1* mutants also suffered from silique abnormalities, in some cases affecting fertility (**Figure 7b-d**). Additionally, we observed during the preparation of ATAC-seq libraries that *met1* mutant nuclei had lower levels of endopolyploidy (**Figure S29**). Homozygous mutants were underrepresented in the progeny of heterozygous parents, consistent with reduced transmission of *met1* alleles, as described previously [6, 56]. To estimate the extent of segregation distortion, we grew up to 96 progeny of heterozygous mutants and genotyped all individuals. In all accessions, homozygotes were underrepresented **(Figure 8a, Table S3)**, from 2% in Aa-0 to 18% in Uk-1.

**Figure 7.**
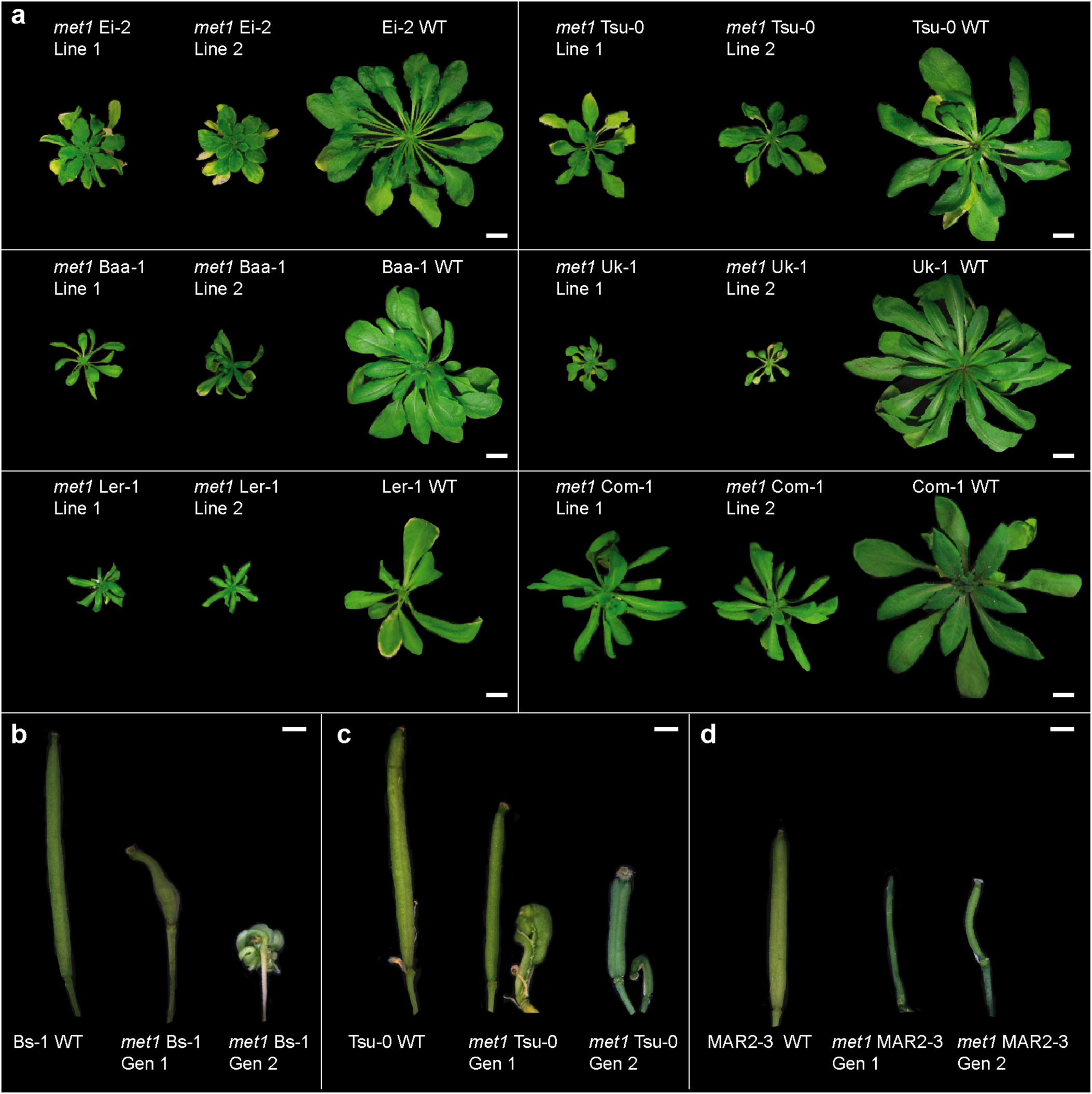
Rosette and silique morphology of *met1* mutants. **(a)** Representative images of two independently derived mutants and the corresponding wild type (WT) for six accessions at six weeks post germination; scale bars represent 1 cm. **(b-d)** Silique morphology in three accessions. Scale bars represent 1 mm.

**Figure 8.**
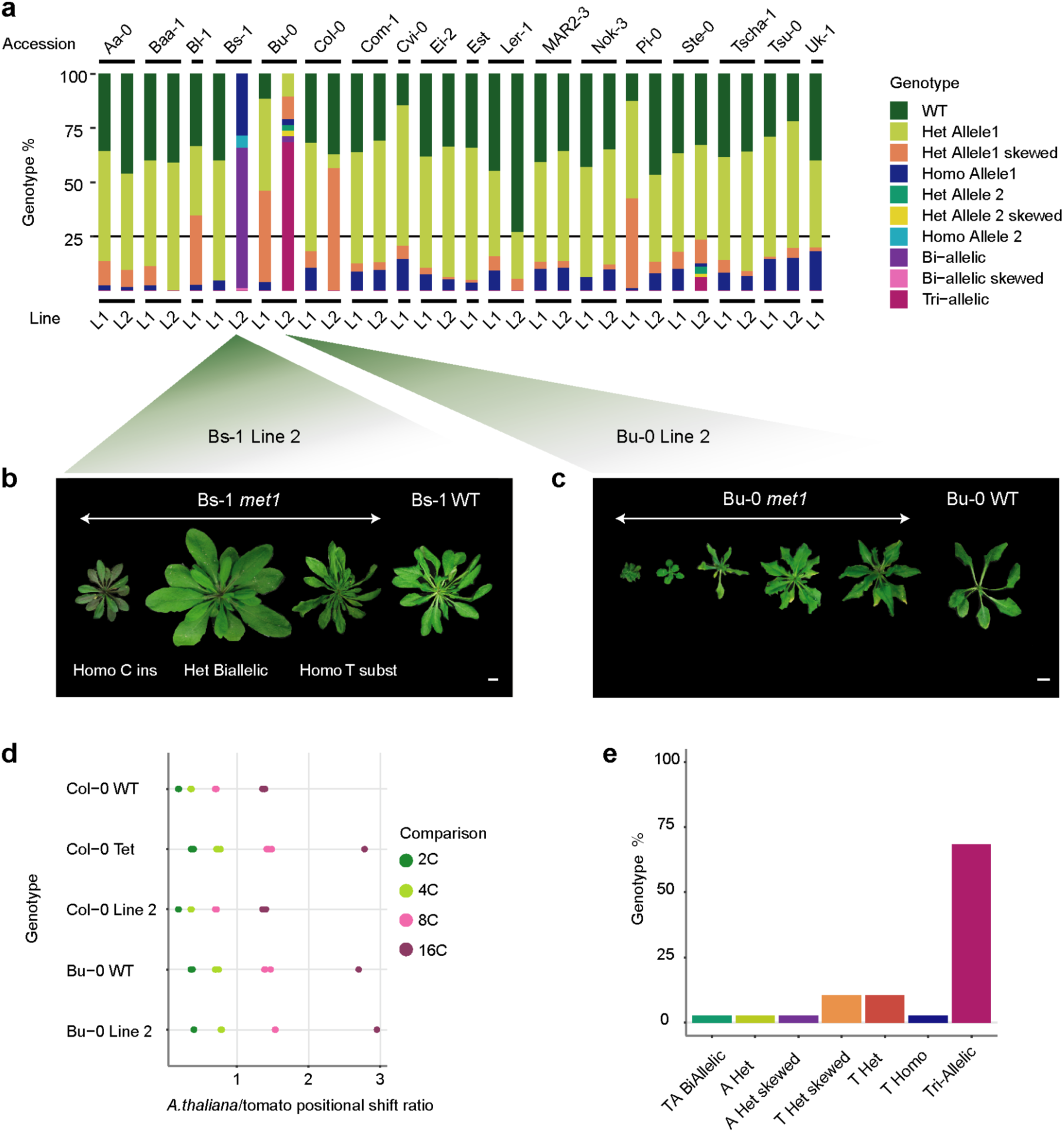
Segregation distortion in *met1* mutants. **(a)** Proportions of wild-type and *met1* mutant genotypes in progeny of heterozygous individuals. L1, L2, Line 1, Line 2. **(b)** Different phenotypes in *met1* Bs-1 Line 2. ’Ins’ refers to ’insertion’ and ’subst’ refers to ’substitution’. **(c)** Phenotypic diversity in Bu-0 Line 2. **(d)** Endopolyploidy peak position ratios (from flow cytometry profiles) in Bu-0 and Col-0 lines relative to tomato internal standard. ’Col-Tet’, Col-0 tetraploid line. **(e)** Fractions of segregating genotypes in *met1* Bu-0 Line 2 progeny. Scale bars in (b) and (c) = 1 cm.

Bs-1 Line 2 included two different types of homozygous mutants that were phenotypically distinct, with the frameshift allele, which we used for the genomic analyses described above, causing more severe phenotypic defects and being more strongly underrepresented in a segregating populations (**Figure 8b**). Bu-0 Line 2 and Ste-0 Line 2 segregated individuals with different combinations of two mutant alleles along with a wild-type allele, pointing not only a biallelic origin of the *met1* alleles but also a change in the generative ploidy level. Genotyping by sequencing also revealed heterozygous individuals with a skewed ratio of reads for the wild-type and mutant alleles, especially in Bu-0 Line 1, Pi-0 Line 1, Col-0 Line 2 and Bl-1 Line 1. Since heterozygotes of these four lines also exhibited phenotypic variation **(Figure 8c, Figure S30)**, we suspected that there was variation in ploidy. Both Bu-0 Line 1 and Line 2 as well as wild-type Bu-0 plants used in this study turned out to be tetraploid, possibly explaining the altered segregation ratios **(Figure 8d, 8e)**. However, other lines with skewed heterozygous allele frequencies were derived from diploid parents **(Figure S30)**, awaiting a more comprehensive explanation of genotypic and phenotypic variation in these lines.

## DISCUSSION

The genomes, methylomes and transcriptomes of different *A. thaliana* accessions can vary substantially [13, 16], and studying their interplay has often focused on TEs, which are silenced by DNA methylation [14, 19]. DNA methylation is also prevalent throughout euchromatin, both near and inside protein-coding genes that are not associated with TEs [23, 57]. MET1 is well known to establish methylation in the predominantly occurring CG-nucleotide context, and cause major phenotypic consequences when its function has been lost. The availability of a collection of *met1* mutants in several accessions enables the investigation of gene expression diversity associated with variation in the parental genomes and epigenomes. We analyzed first- and second-generation homozygous *met1* mutants, finding that the majority of qualitative and quantitative variation in gene expression across accessions arises from genes that are not associated with TEs (Non-TE-DEGs).

The absence of CG methylation can disturb the regulatory balance of hundreds to thousands of genes, and that it does so to a remarkably different extent in genetically diverse backgrounds. Moreover, the comparison of transcriptome diversity across accessions between wild-type and *met1* mutants shows that CG methylation by MET1 can mask underlying genetic diversity at some genes, but at the same time increase expression diversity at another, albeit smaller set of genes.

When analyzing a consensus set of DEGs from all accessions together, we systematically find that the effects of MET1 inactivation can be linked to the initial epigenetic state of the wild-type parent. Genes that are highly CG methylated, expressed at low levels, and have inaccessible chromatin in wild-type plants are the ones that are most likely to become highly expressed and have greatly increased chromatin accessibility in the corresponding *met1* mutants. This pattern is typical for most TE-DEGs, but also seen at some Non-TE-DEGs. From first principles, genes not associated with TEs are much more likely than those genes associated with TEs to change in expression due to indirect effects of *MET1*, and this is confirmed by Non-TE-DEGs being more variable in their methylation and accessibility changes in *met1* mutants. This is also associated with more variation in gene expression changes: While TE-DEGs are almost always upregulated, Non-TE-DEGs change in both directions in *met1* mutants.

Genomic regions that become more accessible in Col-0 *met1* mutants have been associated with multiple gene groups, classified by their expression changes [10]. In our study, we find that this association can be multi-layered, varying by gene, initial methylation level in wild type, and overall genetic background. Our most important conclusion is perhaps that CG methylation, which requires *MET1* activity, cannot be simply thought of as a factor that masks genetic differences or that increases expression diversity beyond genetic variation, but that its effects are highly context-dependent.

Our results show that the same gene across different wild-type backgrounds can not only exist in multiple epigenetic states, but also that it can vary in its regulatory response to genome-wide CG hypomethylation, thereby unveiling distinct associations between methylation, chromatin accessibility and gene expression. For example, the study by Zhong and colleagues [10] demonstrated how local changes in methylation were sufficient to alter chromatin accessibility at the *FWA* epiallele in the reference accession Col-0. Upon examining the *FWA* locus in homozygous *met1* mutants in different accessions, we find that the increases in accessibility can highly vary despite a similar degree of CG hypomethylation in each *met1* mutant, starting from different initial parental methylation levels **(Figure S23)**. Additionally, the relationship between accessibility changes and expression changes is non-linear for this gene **(Figure S23)**, suggesting that either variation in cis-regulatory sequences or trans regulators make an important contribution to *FWA* expression beyond methylation.

*FWA* is only one example from 11,675 unique DEGs in our set of *met1* mutants from 18 accessions (compared to 1,759 DEGs, when comparing only Col-0 wild type and *met1*). That only 291 DEGs among these 11,675 unique DEGs are universal across all accessions highlights the differential sensitivity of each genetic background to methylation-dependent changes in gene expression. Accession-specific DEGs include developmental and epigenetic regulators such as *ROS1*, *IBM1* and *SUVH3*, many of which feature well-examined epialleles. We note that because we only sampled leaves, we almost certainly have not discovered all genes that are sensitive to loss of *MET1*, and indeed many other tissues besides leaves are phenotypically affected in *met1* mutants.

Methylation and chromatin accessibility changes are clearly neither necessary nor sufficient for changes in gene expression in *met1* mutants, again revealing how methylation and chromatin architecture must interact with cis-regulatory sequences to effect gene expression [58]. It will be interesting to determine whether genes with altered methylation and chromatin accessibility are more likely to change in their expression during further cycles of propagation. In addition, deeper investigation of non-CG and residual CG methylation in *met1* mutants, catalyzed by other methyltransferases and compensatory pathways, may further improve our understanding of methylation-induced gene regulation.

A wide range of silique abnormalities and distorted segregation ratios observed in our *met1* mutants indicates that the absence of MET1 function may be detrimental for gametogenesis, fertilization or post-zygotic development, at least in some accessions. Among the accessions used in our study, Ler-1 and Tsu-0 carry a microRNA haplotype that impairs silencing of specific TEs in male gametes (Ler-0 *MIR845* haplotype [59]). Furthermore, epigenetic variation across Col-0, Ler-1 and Cvi-0 has been shown to impact seed development due to differential imprinting [60, 61]. For example, the comparatively higher fertility of *met1* mutants in Cvi-0 could be explained by the natural hypomethylation of the *HDG3* (*AT2G32370*) locus in wild-type Cvi-0 plants **(Figure S26)**, thereby reducing dramatic effects in seed size as observed in mutants of other accessions. Future studies on the epigenetic and epigenomic landscape of gametophytic, embryonic and endosperm tissue of our *met1* mutants will shed light on how hypomethylation in somatic cells can impact fertilization processes.

## CONCLUSION

The absence of *MET1* in *A. thaliana* has long been known to affect the chromatin landscape [7,8,10,11]. Here we show the value of studying accessions beyond the reference accession Col-0, finding not only that CG methylation, gene expression and chromatin accessibility can widely vary, but also that the impact of *MET1* can be much greater in other accessions, especially at genes that are not associated with TEs.

Our *met1* mutant collection provides future opportunities for investigating how epigenetic and epigenomic regulation at individual loci are fine-tuned across accessions to determine plant phenotype. Another opportunity will be to study the TE mobilization landscape in these mutants, and to ask whether sites with newly inserted or newly excised TEs have similar epigenetic states across different genetic backgrounds and how these in turn affect adjacent protein-coding genes [62]. Finally, while the resources and insights we have generated have already improved our understanding of the variation in epigenetic and epigenomic regulation that evolution has produced, high-quality genome assemblies of the studied accessions will make our resources even more valuable.

## METHODS

### CRISPR/Cas9 knockout of *MET1* in 18 *A. thaliana* accessions

Using a plant molecular cloning toolbox [28], a supermodule destination binary vector carrying a plant-codon optimized *Cas9* driven by a *UBQ10* promoter was cloned with a single guide-RNA (gRNA) targeting the *A. thaliana MET1* (*AT5G49160*) gene. The gRNA was designed using the CRISPR design tool in *Benchling* (www.benchling.com) targeting a 20 bp region in exon 7 of *MET1* (**Table S1**), which is the same exon where previously described *met1-3* mutants are known to harbor a T-DNA insertion [6]. This exon is present in the catalytic domain of the protein and harbors a motif that is a binding site for cytosine nucleotide substrates [63]. Eighteen early-flowering *A. thaliana* accessions were transformed with the above CRISPR construct by *Agrobacterium-*mediated floral dipping [64], carried out twice with a 7-10 day interval. Seeds of primary transformants (T_1_) were screened for the presence of the transgene by selecting for the mCherry fluorescence marker, and sown on soil. These T_1_ plants were subjected to heat treatment cycles for enhancing Cas9 activity [65]. Genotyped lines carrying a mutation in the gRNA-target region were propagated to the T_2_ generation after segregating the transgene (by selecting for non-mCherry seeds), followed by identification of lines carrying heritable heterozygous mutations. 1-2 heterozygous T_2_ lines per accession were further subjected to one or two more rounds of propagation to identify first generation homozygous plants in the segregating progeny.

### Genotyping transformants and identification of homozygous mutants

First- and second-generation homozygous mutants were genotyped either using Sanger sequencing of a 649 bp PCR-amplicon, or by amplicon-sequencing of a 152 bp PCR-amplicon **(Table S1)**, both covering the CRISPR guide-RNA target region. Most of the mutations identified in all mutants were single bp insertions or deletions that occurred within the first four bp from the 5’ end of the target region, and disrupted the open reading frame due to frameshifting. Candidate homozygous mutants in segregating T_3_ populations were first identified by visual phenotyping, followed by genotyping.

### Plant growth conditions and tissue collection for large-scale sequencing

Seeds were sterilized by treatment with chlorine gas for 4 hours, followed by stratification in the dark at 4**°**C for 4 days in 0.1% agar. All plants were grown in controlled growth chambers at 23 **°**C, long day conditions (16 h light/8 h dark) with 65% relative humidity under 110 to 140 μmol m^−2^ s^−1^ light provided by Philips GreenPower TLED modules (Philips Lighting GmbH, Hamburg, Germany) with a mixture of 2:1 DR/W LB (deep red/white mixture with ca. 15% blue light) and W HB (white with ca. 25% blue light), respectively, and watered at 2-day intervals.

Since homozygous mutants from several accessions had reduced fertility and did not set sufficient seeds for further propagation, sampling for all sequencing experiments was carried out in the first homozygous generation. Segregating populations of T3 mutant plants were grown from 1-2 lines per accession, and homozygous individuals were marked by their distinct phenotype (as identified in previous growth experiments) and later confirmed by Sanger sequencing of DNA used for BS-seq libraries. Wild-type plants from the same accessions were grown in parallel to all mutant lines.

At 25 days after germination, three homozygous individuals per parental line per accession were collected as separate biological replicates, along with three wild-type individuals. Sampling involved the collection of two sets of rosette leaves from the same individual plant. One set of leaves was immediately frozen in liquid nitrogen containers (and subsequently at -80°C) to be homogenized and split for Bisulfite-sequencing and RNA-sequencing analysis. The second set of leaves were collected for ATAC-sequencing analysis and were subjected to syringe-infiltration with 0.1% formaldehyde in phosphate-buffer saline, followed by 0.125 M glycine in phosphate-buffer saline, washed with autoclaved water and dried before storage at -80°C. All tissue sampling and fixation was carried out within a 30 minute time window.

### Bisulfite-seq library prep

Frozen leaf tissue from three biological replicates were mixed and grinded together. This powder was used for isolating genomic DNA using the DNeasy Plant Mini Kit (Qiagen). 100 ng of this genomic DNA was subsequently used to prepare Bisulfite libraries with the TruSeq Nano kit (Illumina, San Diego, CA, USA) according to the manufacturer’s instructions, with the modifications used in [66]. The libraries were sequenced in paired-end mode, with approximately 8.5 million 150 bp reads/library on an Illumina HiSeq3000 instrument.

### Processing of Bisulfite-seq data and DMR calling

Raw BS-seq reads were aligned using Bismark with default parameters [67] and mapped to the *A. thaliana* (TAIR10) reference genome. The bisulfite conversion efficiency for each sample was estimated by evaluating the fraction of positions correctly called as unmethylated in the chloroplast genome. It was consistently above 99.6% in all samples. The mapping efficiency for all samples varied between 40% and 65%, with an average of 49% (**Appendix 1**). Deduplicated bam files generated by Bismark were sorted and then processed using *MethylScore* (https://github.com/Computomics/MethylScore; [68]) in the CLIP cluster of the Vienna Biocenter (VBC), to identify DMRs (Differentially Methylated Regions) across all 73 samples (55 mutants and 18 wildtypes) with the following parameters: DMR_MIN_C=10 (minimum 10 cytosines in each DMR), DMR_MIN_COV=3X (minimum 3X coverage in each cytosine), MR_FREQ_CHANGE=20 (at least 20% of samples showing a change in MR frequency to be tested as a candidate DMR), CLUSTER_MIN_METH_DIFF=20 (which sets a 20% cutoff for methylation difference between clusters in the CG, CHG and CHH contexts). All other parameters were based on default settings.

DMR coordinates were intersected with individual genome-wide cytosine methylation levels based on the *genome_matrix* file generated by MethylScore. A total of 2,836 All-C DMRs (in all 3 contexts), 2,388 CG DMRs, 350 CHG DMRs and 1,023 CHH DMRs were called across 73 samples (55 mutants and 18 wildtypes) (**Appendix 2**). In some cases, a DMR was enriched for more than one context, resulting in partial redundancies. Subsequently, the three context-specific DMRs (CG, CHG, CHH) were evaluated for context-specific average methylation levels by intersecting with sample-specific cytosine methylation data. This was achieved using the *bedtools* software (v2.26.0) with the following command:

~~~
*bedtools map -a DMR_coordinates.bed -b methylated_cytosines_sampleX.bed -c 5 -o mean - nonamecheck -null “NA” -g TAIR10genomesize > DMR_Methavg_sampleX.bed*
~~~

Although the DMR calling was performed with a three-read cutoff for each cytosine in a DMR, there remained some samples which did not have sufficient coverage. Therefore, we retained the same DMRs, but calculated average methylation by lowering the cutoff to 2 reads per cytosine. For downstream analysis, DMRs with a maximum of 7 NAs (insufficient coverage) out of 73 samples were retained. This resulted in 1,966 AllC DMRs, 1,569 CG DMRs, 207 CHG DMRs and 614 CHH DMRs (**Appendix 2**), which were used for intersecting with dACRs and DEGs.

### Intersections between DMRs, transposable elements and Non-TE genes

DMR positions were intersected with positions of TAIR10 transposable elements and non-TE genes (TAIR10 genes that are not associated with the term “transposable-element-gene”) using *bedtools intersect*, with a minimum overlap of 1bp (**Table S2**).

### Nuclei isolation for ATAC-seq

For ATAC-seq analyses, each of the biological replicates was processed individually. Fixed tissue was chopped finely with 500 µl of General Purpose buffer (GPB; 0.5 mM spermine•4HCl, 30 mM sodium citrate, 20 mM MOPS, 80 mM KCl, 20 mM NaCl, pH 7.0, and sterile filtered with 0.2 µm filter, followed by the addition of 0.5% of Triton-X-100 before usage). The slurry was filtered through one-layered Miracloth (pore size: 22-25 µm), followed by filtration twice through a cell-strainer (pore size: 40 µm) to collect nuclei.

### Fluorescence-activated cell sorting (FACS) for ATAC-seq

Liberated nuclei were sorted with a MoFlo XDP (Beckman Coulter) instrument outfitted with a 488 nm (elliptical focus, 100 mW) and a 375 nm (spherical focus, 35 mW) laser for scatter and DAPI emission, respectively. Nuclei were sorted with a 70 µm cytonozzle, sheath PBS [pH 7.0] at psi 30.5/30.0 sample/sheath, purify 1 drop, triggered off the DAPI emission (465/30nm). The 2C endoreduplicated population was identified as the first clear DAPI emitting population over scatter debris. DAPI emission was utilized to reduce further contaminating debris, followed by a clean-up utilizing 530/34 emission from the 488 nm laser. 488 nm channels: SSC (488/6), FL1 (520/34). 375 nm channels: FL8 (405/30), FL9 (465/30), FL10 (542/27). See **Figure S29** for the gating scheme. Approximately 20,000 DAPI stained nuclei were sorted using fluorescence-activated cell sorting (FACS) as two technical replicates. For samples from dwarfed mutant lines where leaf tissue was scarce, approximately 8,000 nuclei were sorted per technical replicate.

### ATAC-seq library prep

Sorted nuclei were heated at 60°C for 5 minutes, followed by centrifugation at 4°C (1,000 g, 5 minutes). The supernatant was removed, and nuclei were resuspended with a transposition mix (1 µl homemade Tn5 transposase, 4 µl of 5X-TAPS-DMF buffer and 15 µl autoclaved water) followed by a 37°C treatment for 30 minutes. 200 µl SDS buffer and 8 µl 5 M NaCl were added to the reaction mixture, followed by 65°C treatment overnight. Nuclear fragments were then cleaned up using Zymo PCR column-purification (DNA Clean and Concentrator). 2 µl of eluted DNA was subjected to 14 PCR cycles, incorporating Illumina indices, followed by a 1.8:1 ratio clean-up using SPRI beads.

Genomic DNA libraries (10 ng input from the DNA extracts used for BS-seq-library prep) were prepared using a similar library prep protocol starting with Tn5 enzymatic digestion (0.5 µl homemade Tn5 transposase, 4 µl of 5X-TAPS-DMF buffer and autoclaved water made up to a final reaction volume of 20 µl including the DNA template). Digested gDNA was immediately column-purified, followed by PCR (2 µl of eluted DNA was used as template for 11 PCR cycles) incorporating Illumina indices, followed by a 1.6:1 ratio clean-up using SPRI beads.

### Processing of ATAC-seq libraries and peak-calling

Libraries were sequenced on an Illumina HiSeq3000 instrument with 2 x 150bp paired-end reads. Each technical replicate derived from nuclei sorting was sequenced at approximately 7 million paired-end reads per library. The reads were aligned as two single-end files to the TAIR10 reference genome using *bowtie2* [default options], filtered for the SAM flags 0 and 16 (only reads mapped uniquely to the forward and reverse strands), converted separately to bam files. The bam files were then merged, sorted, and PCR duplicates were removed using *picardtools.* The sorted bam files were then merged with the corresponding sorted bam file of a second technical replicate (samtools merge --default options) to obtain a final average of 11 million mapped reads for each biological replicate. Genomic DNA libraries were similarly aligned, with an average of 4.5 million mapped reads per library (**Appendix 1**). Peak calling was carried out for each biological replicate using *MACS2* (**Appendix 2**) using the following parameters:

*macs2 callpeak -t [ATACseqlibrary].bam -f BAM --nomodel --extsize 147 --keep-dup=all -g 1.35e8 -n [Output_Peaks] -B -q 0.01*

After peak calling, every peak set was further filtered based on their respective q-values in the MACS2 peaks.xls files, retaining peaks with q <= 0.001, thereby reducing the false positives when all 158 samples were subsequently tested together. This additional filtering step was carried out separately after MACS2 calling to minimize the effect on peak size based on the q-value.

Filtered peak files and .bam alignment files from a total of 158 samples (104 mutant samples plus 54 wild-type samples) were processed with the R package DiffBind to identify consensus peaks which overlapped in at least two out of three biological replicates per group, and represented peaks unique to at least one group (FDR adjusted p-value <0.01). To normalize peak accessibility counts with the background probability of Tn5 integration biases in the genome, .bam files of the control gDNA libraries were also provided in the DiffBind sample sheet (thereby ensuring that peak accessibility counts were normalized to controls).

A total of 35,049 consensus peaks were identified, with accessibility scores in each peak per sample evaluated in counts per million (CPM) after TMM (Trimmed Mean of M-values) normalization. Except for three out of 158 ATAC-seq libraries, FRIP (Frequency of reads in peaks) scores relative to the Consensus peak set was between 0.2 - 0.31 for all samples, reflecting the average representation of sample-specific peaks in the consensus dataset. After removing peaks which occurred in Chloroplast and Mitochondrial genomes, 34,993 peaks remained.

### Metaplot generation

From 34,993 consensus ATAC-peaks (pre-filtered), accessibility values for each peak region were derived for all samples, and converted into bigWig files using the *bedGraphtobigWig* command (UCSC software).

Similarly, cytosines in all-contexts and their corresponding methylation levels were derived for each Bisulfite library, and converted to bigWig files. Metaplots were generated using the deepTools (3.5.0) package (https://deeptools.readthedocs.io/en/develop/index.html), first with the *computeMatrix* function to evaluate the mean value of the epigenetic factor tested (methylation in % or chromatin accessibility in CPM) across 10 bp non-overlapping bins, within 1000 bp upstream and downstream of a given set of reference regions (TAIR10 Transposable elements/TAIR10 Non-TE protein coding genes). The output bed file from this command was subsequently used to generate metaplots using the *plotProfile* function.

### RNA extraction and RNA-seq library prep

RNA from each biological replicate was extracted individually using a column-based protocol adapted from [69]. RNA quality was validated with the Nanodrop spectrophotometer, and normalized to 500 ng in a 50 µl volume. Normalized RNA was subsequently used for mRNA library prep using an in-house custom protocol adapted from Illumina’s TruSeq library-prep, with details provided in [70].

### Mapping and identification of DEGs

RNA-seq libraries were sequenced at an average coverage of 8 million 150 bp single-end reads per library using HiSeq3000. Reads of the same sample from multiple sequencing lanes of the same flow cell were merged together, and 9 samples with > 12.5 million total reads were subsampled (using different seeds) to 80% using seqtk (v.2.0-r82-dirty, https://github.com/lh3/seqtk) with the following command:

~~~
*seqtk sample -sX <MERGED_FASTQ> 0.80 > subsampled_output.fastq*
~~~

All samples were aligned using *bowtie2* to the TAIR10 reference genome, prepared using the *rsem-prepare-reference* function of the RSEM software. Aligned bam files were sorted and indexed using *samtools* V1.9 (mapped reads given in **Appendix 1**). Gene transcript counts for each sample were estimated using *rsem-calculate-expression*. From each sample, chloroplast genes, mitochondrial genes, and rDNA cluster genes were excluded from downstream analyses. Twelve genes with excessive read counts across all samples were also excluded (**Appendix 1**).

Transcript counts per sample and corresponding metadata were then imported using the R packages “tximport” and “tximportData” for creating a DESeq object (R package “DESeq2”).

Identification of DEGs (differentially expressed genes) between 18 accessions for *met1* mutants and wildtypes: Two DESeq objects containing 104 samples of *met1* mutants and 54 samples of wildtypes were generated separately. After filtering genes with low read counts, a common set of 19,473 genes were retained in both objects, and the DESeq function was applied under a one-factor model (∼Accession) and default parameters (nbinomWald test). DEGs were identified from the DESeq output as those genes with a p value < 0.01 and |log_2_ FoldChange| >1.

Identification of DEGs between *met1* mutants and wildtypes: The DESeq function was applied to the object under the two-factor interaction model ∼*genotype + accession + genotype:accession* (where *genotype ==* wild-type *or mutant*) using default parameters (nbinomWald test). To obtain DEGs between wild-type and mutant genotypes across all accessions, a contrast was performed set to *genotype*, retaining only those genes with a p value< 0.01 and |log_2_ FoldChange| >1. For identifying accession-specific DEGs, similar contrasts were performed, nested within each accession.

A consensus set of 7,132 DEGs were derived from all 18 accession specific contrasts and the all-mutants-against-all-wild-type contrast, retaining only DEGs that occurred in at least two of all 19 contrasts. These were further classified as 1,401 TE-associated DEGs (TE-DEGs) and 5,731 Non-TE-DEGs based on the TAIR10 gene annotation (“TE genes” (3903) defined as those with the term “transposable-element-gene”, as described in https://www.arabidopsis.org/portals/genAnnotation/gene_structural_annotation/annotation_data.jsp. All remaining 29699 protein-coding genes were called “Non-TE genes”).

### Weighted gene co-expression network analysis

Transformed and normalized RNA-seq counts (vsd counts) were extracted for 10,151 unique Non-TE-DEGs identified from all 19 contrasts, across 158 samples. This matrix was then used to generate a weighted gene co-expression network using the R package WGCNA [71]. A soft threshold power of 4 was chosen, and subsequently used for constructing the TOM (topological overlap matrix) with minModuleSize set to 30 and mergeCutHeight set to 0.25. This resulted in the generation of 9 distinct gene modules. Module eigengenes were examined for correlation to the sample genotype (1 for mutant and 0 for wild-type). The highest eigengene significance (0.9) was observed for Module ’D’ with 814 genes (**Appendix 2**).

### Generation of consensus datasets

#### Consensus DEGs

These were generated by including DEGs from a total of 19 contrasts - 18 pairwise contrasts (*met1* mutants vs wild-type plants) in each accession, and a contrast between all *met1* mutants and all wild-type plants. This set of DEGs was filtered to retain only DEGs that occurred in at least 2 out of the total 19 contrasts, to obtain a final set of 7,132 Consensus DEGs. These were further classified as 1,401 TE-associated DEGs (TE-DEGs) and 5,731 Non-TE-DEGs (protein-coding) (**Appendix 2**). The metric for evaluating expression levels in DEGs was chosen as the variance stabilized transformed read counts (vsd counts) generated by the DESeq2 package for each of the 158 RNA-seq libraries (**Appendix 2**). We refer to these as transformed read counts in all figures.

#### Consensus dACRs

ATAC-peaks found across all samples (34,993) were also filtered in two steps. First, peaks that showed similar accessibility between both mutant lines relative to the WT line in each accession were retained. The second round of filtering retained peaks that showed an accessibility change between mutants and wild-type plants in at least two out of 18 accessions. This resulted in 31,295 filtered consensus ATAC-peaks, which we refer to as differential ACRs (dACRs) (**Appendix 2**). Chromatin accessibility levels were measured in counts per million (CPM) after TMM (trimmed mean of M-values) normalization generated by the DiffBind package. K-means clustering of consensus dACRs was carried out using the functions *kmeans()* in the R *stats* package, with k = 5.

#### Consensus DMRs

From the complete set of context-specific DMRs identified, DMRs with a maximum of 7 NA values (insufficient coverage) out of 73 samples were retained. This resulted in 1,966 All-C-DMRs, 1,569 CG DMRs, 207 CHG DMRs and 614 CHH DMRs (**Appendix 2**). Since we were primarily interested in understanding the effects of MET1 on genome-wide methylation changes, we considered only CG-DMRs and All-C-DMRs for intersecting with other features. While both groups contained some redundant DMRs, the methylation level evaluated for each DMR was evaluated differently (for C-DMRs, methylation level evaluated only for CG context; for All-C-DMRs, methylation level evaluated for all Cs in CG, CHG, CHH contexts). Methylation levels in DMRs were measured as arithmetic mean over methylation percentage of all context-specific and sample-specific cytosines within the assigned chromosomal region.

### Generation of feature intersections between DEGs, DMRs and dACRs

#### DMR - DEG intersections

DMRs occurring at the extended gene body (between 100bp upstream of the TSS and 100bp downstream of the TTS of a gene) were called ’gene-body DMRs’, while those that occurred within 1.5kb upstream or downstream of the TSS/TTS respectively were named ’cis DMRs’. Several DEGs had multiple DMRs associated with them, and therefore we retained only one DMR for each DEG, which showed the largest difference in methylation level between mutant and WT for each mutant genotype, thereby aiming to represent only the strongest methylation signals that could explain gene expression differences. To determine the extent at which MET1-induced CG methylation could influence gene expression, and compare it with methylation in all contexts, we intersected DEGs (TE-DEGs and Non-TE-DEGs separately) with CG-DMRs. For each DEG, we measured changes in gene expression levels between *met1* mutants and wild-type plants, and corresponding changes in methylation levels of their closest DMRs. For generating scatter-plots, we divided wild-type methylation/expression levels of all examined genes into quintiles and colored points based on these quintiles.

#### dACR - DEG intersections

Similarly, we analyzed the dACRs closest to each DEG. Since a large majority of dACRs occurred in proximity to the transcription start site, we grouped all dACRs occurring either over the gene body or within 1.5 kb upstream or downstream of the TSS/TTS respectively, under a single ’cis’ category. For each DEG, a single dACR which showed the largest difference in accessibility between mutant and WT for each mutant genotype was retained. We next measured changes in gene expression levels between *met1* mutants and wild-type plants, and corresponding changes in accessibility levels of their *cis* dACRs. For generating scatter-plots, we divided wild-type accessibility/expression levels of all examined genes into quintiles and colored points based on these quintiles.

#### Intersections of DMRs and dACRs with Non-DEG genes

As a control for the DEGs, we generated similar feature intersections for Non-DEGs as well. From a total of 1,678 TE genes and 19,979 Non-TE genes analyzed using DESeq2, we identified 277 TE genes and 14,248 Non-TE genes which were not classified as ’Consensus DEGs’, and we subsequently referred to these as ’Non-DEGs’. To ensure that the number of Non-DEGs analyzed were comparable to the number of DEGs in each category, 5,731 Non-TE genes were randomly subsampled from the total set of 14,248 Non-TE (Non-DEG) genes. However, only 250 TE genes were randomly subsampled from the total set of 277 TE (Non-DEG) genes, since the total number of TE-DEGs (1,401) exceeded the number of TE (Non-DEG) genes.

#### DMR - dACR - DEG intersections

For simplicity, the above class of three-way intersections was only carried out for CG DMRs and Non-TE genes. In short, Non-TE-DEGs carrying CG DMRs in both the extended gene-body and in *cis* were combined together. These combined DEGs were then filtered to identify only those which carried a dACR in *cis*. This resulted in a final set of 407 DEGs which carried both a CG DMR and a dACR in *cis*. Similarly, when a control set of Non-TE Non-DEG genes were used to generate similar intersections, a final set of 407 genes carrying both a CG DMR and a dACR in *cis* were identified.

### gbM-like and CG teM-like genes

To follow conventional definitions of gene body methylation (gbM), only Non-TE-DEGs with CG-DMRs overlapping the gene body were considered. These criteria were satisfied by 196 DEGs across homozygous *met1* mutants in 17 accessions (since one accession, Bl-1, did not have homozygotes). From this set of genes, we identified 91 gbM-like genes, which had >80% methylation in the wild-type parent, and transformed expression counts in wild type >=6.53, which represented genes in the top two quintile range of the distribution of wild-type expression levels for all 196 genes.

Next, we identified 51 CG teM-like genes that exhibited >80% CG methylation in the wild-type state, and transformed expression counts in wild-type state being <=4.39, which represented genes in the lowest quintile range of the distribution of wild-type expression levels for all 196 genes.

Metadata (transformed expression counts and methylation levels in *met1* and mutants wild-type plants) for gbM-like and CG teM-like genes were then extracted from the total set of 196 genes (thereby representing the same genes in all accession backgrounds) and used for generating visual plots.

### Gene Ontology Enrichment and Visualization

GO Enrichment was carried out using agriGO (http://bioinfo.cau.edu.cn/agriGO), with the Singular Enrichment Analysis (SEA) analysis tool and Arabidopsis (TAIR10) gene model as the reference. Graphical results of significant GO terms were generated in agriGO. The GO terms were further visualized with ReviGO (http://revigo.irb.hr).

### Segregation distortion analyses

#### Experimental design

To accurately estimate the extent of this segregation distortion in mutants of various accessions, we grew a maximum of 96 segregating progeny from heterozygous parent lines (2 mutant lines per accession) and genotyped them individually using amplicon-sequencing of the *MET1* locus, amplifying a 150 bp region around the CRISPR/Cas9-induced frameshift mutations.

#### Amplicon-seq library preparation, sequencing and genotyping

Amplicon-seq libraries were prepared according to the CRISPR-finder system [72], where amplicons from multiple 96-well plates can be pooled together for high-throughput sequencing by incorporating frameshifted primers and TruSeq adapters with 96 barcodes. The amplicons were designed as 152 bp sequences spanning the gRNA target site in the *MET1* gene locus **(Table S1)**. A total of 2,788 individual samples were sequenced at an average coverage of 12,000 reads per sample on a HiSeq3000 instrument with 2 x 150 bp paired-end reads.

Sequenced read pairs were first merged using *FLASH* (*Fast Length Adjustment of Short reads*) (https://ccb.jhu.edu/software/FLASH/), followed by demultiplexing based on plate-specific frameshifted primers (see **Table S4**) using the *usearch10 fastx_truncate* function. Only samples with >=80 reads were retained for downstream processing. For all samples within a plate (i.e., segregating progeny), amplicon-reads per individual were counted for the ratio of wild-type alleles to mutant alleles, to estimate whether the genotypes were Homozygous for the mutant allele (wild-type reads <= 15%), heterozygous (wild-type reads >= 42% or <= 58%) or wild type (wild-type reads >= 90%). A fourth genotypic classification, “skewed heterozygous” was made for individuals where the read ratio between the wild-type and mutant alleles were either 0.15 - 0.42 or 0.58 - 0.90 (i.e., if either one of the alleles were more represented than the other, but not approximately equal in counts).

For Bu-0 Line 2, Ste-0 Line 2 and Bs-1 Line 2 samples that had more than one mutant *MET1* allele, additional genotypic categories were specified: homozygous allele 2, heterozygous allele 2, skewed heterozygous allele 2, bi-allelic, skewed bi-allelic and tri-allelic **(Table S3)**.

### Cytometric ploidy analysis

Cytometric determination of generative ploidy levels was conducted on a CytoFlex (Beckman Coulter) outfitted with a 488 nm laser, 10 µL min^-1^ flow rate. Nuclei were freshly liberated by chopping into cold General-purpose Buffer [73], filtered through 40 µm mesh, and stained with 50 µg mL^-1^ propidium iodide and 50 µg mL^-1^ RNase for 10 minutes at 20°C. The 2C endoreduplication population was identified as the first clear PI emitting population over scatter debris. The 2C nuclei of *Solanum lycopersicum* (var. Moneymaker) provided by the Zentrum für Molekularbiologie der Pflanzen (ZMBP) Cultivation Facility and *Capsicum annuum* provided by Annett Strauss (ZMBP) were used as internal standards to determine the relative Arabidopsis generative ploidy levels.

## DATA AVAILABILITY

Raw Data Availability: SRA links for raw sequencing data have been deposited at ENA (European Nucleotide Archive, https://www.ebi.ac.uk/) under the accession numbers PRJEB53354 (RNA-seq data), PRJEB54034 (ATAC-seq data), PRJEB54036 (Bisulfite-Seq data) and PRJEB54071 (Amplicon-seq data).

## ACKNOWLEDGEMENTS

We are grateful to Chang Liu for guidance with ATAC-seq experiments and analyses. We thank Anjar Wibowo for advice and valuable discussions on the propagation of *met1* mutants and the experimental setup of the project during its initial stages. Claude Becker, Isaac Rodriguez and Patrick Hüther (GMI Vienna and LMU Munich) kindly provided access to and answered our queries about the ’MethylScore’ pipeline for DMR calling before publication. We thank current and former members of the Weigel lab for helping with tissue collection and nuclei extraction. Large scale nuclei extractions, and preparation of large scale amplicon-seq libraries benefitted from 3D printed custom tools designed by Dr. Ulrich Lutz (Custom Lab Institute). Finally, we acknowledge valuable feedback from Anjar Wibowo, Leandro Quadrana and Vincent Colot. This study was supported by the Max Planck Society.

## AUTHOR CONTRIBUTIONS

Study design: T.S. and D.W. All experimental work except flow cytometry: T.S. Flow cytometry: K.W.B. Data analysis: T.S. Data interpretation: T.S., W.Y., H-G.D., R.S., D.W. Drafting of initial manuscript: T.S. Editing and finalizing of manuscript: T.S., R.S., W.Y., K.W.B., H-G.D., A.C.G, D.W.

## SUPPLEMENTARY FIGURES

**Figure S1.**
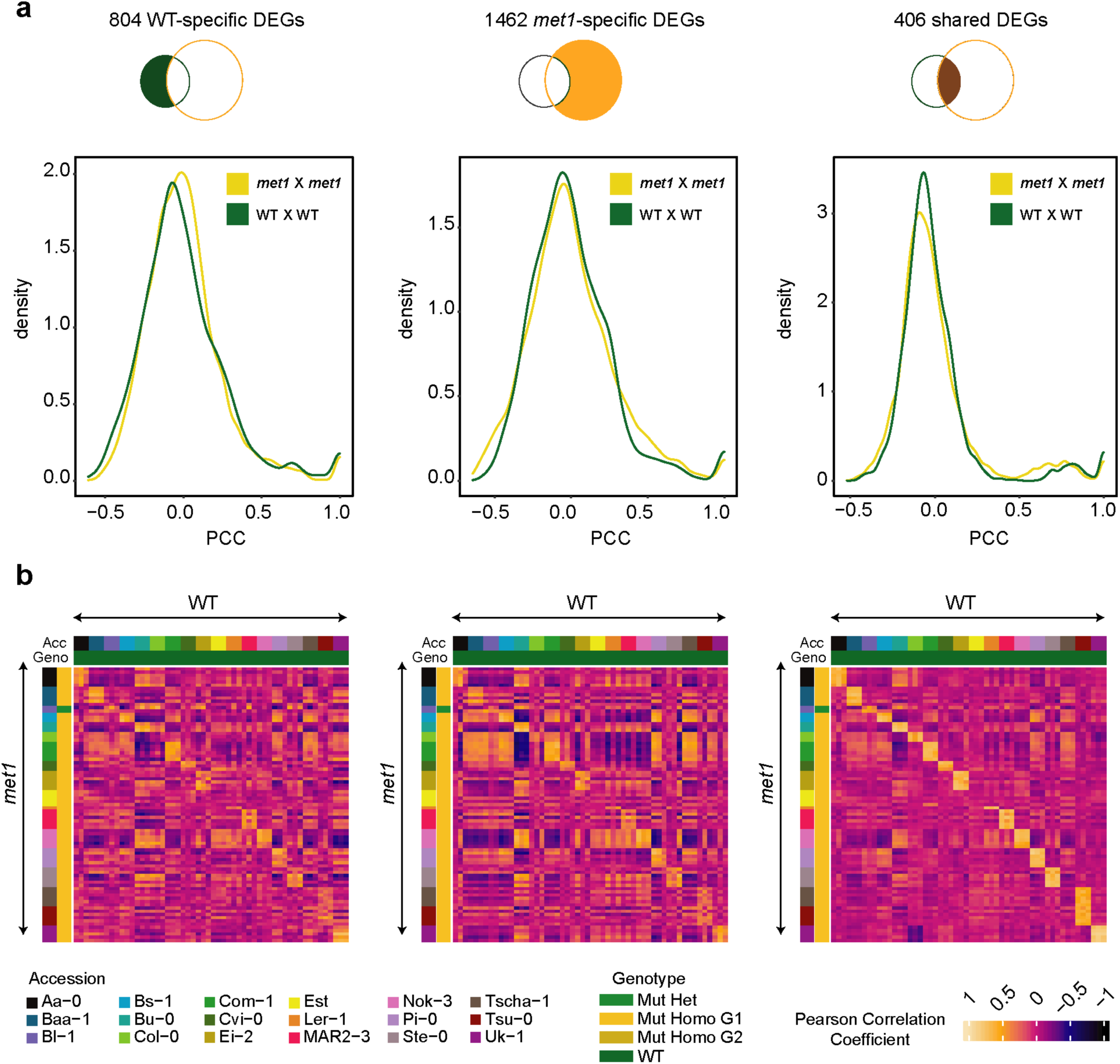
Transcriptome changes in *met1* mutants are gene- and accession-specific. (a) Density distributions of Pearson Correlation Coefficients among *met1* mutants of different accessions and wildtypes of different accessions respectively, for three different groups of DEGs. (b) Heatmaps representing Pearson Correlation Coefficients for *met1* mutants against wildtypes for three different groups of DEGs.

**Figure S2.**
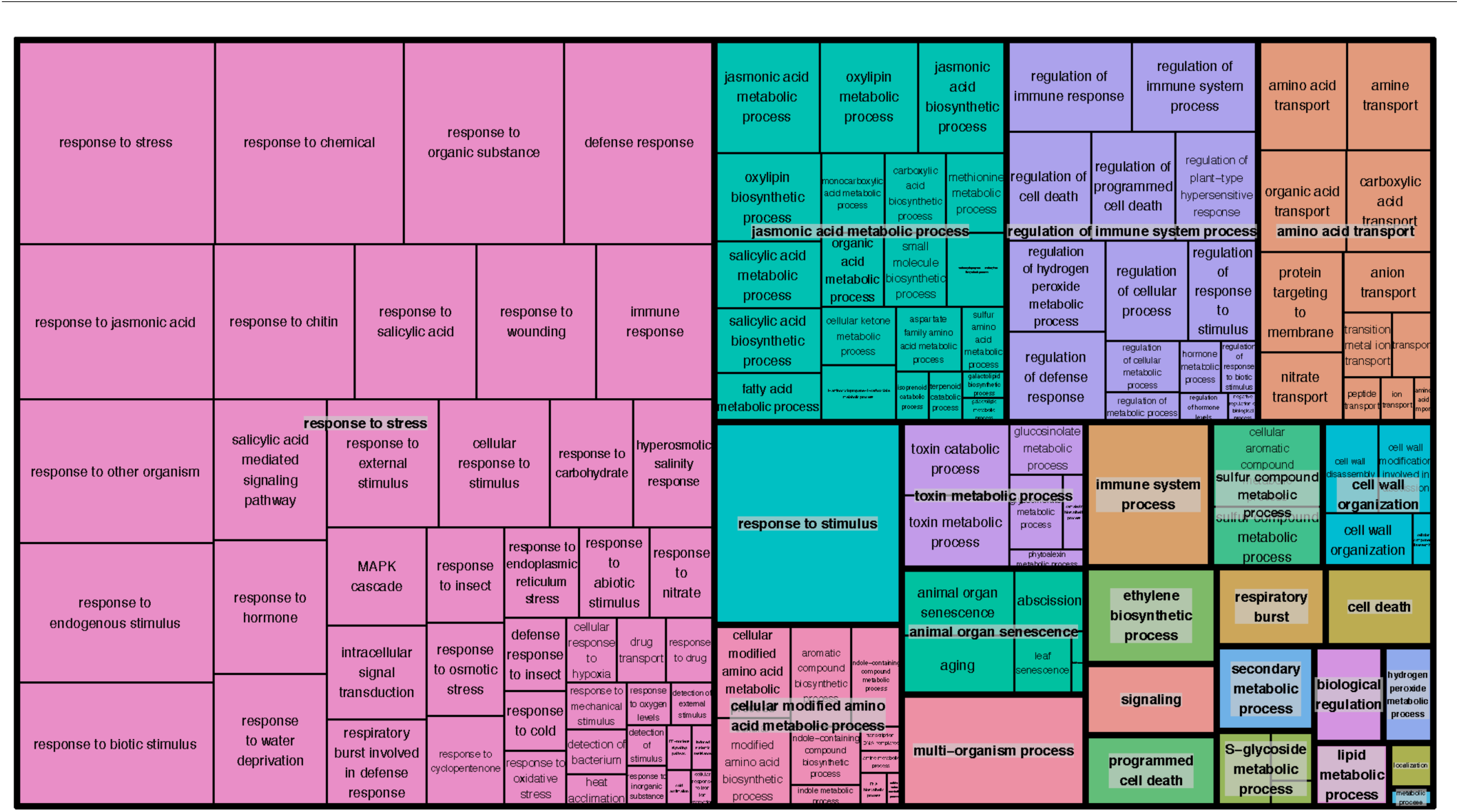
Gene Ontology enrichment for 2,013 Non-TE-DEGs identified in a contrast between all *met1* mutants and all wild-type samples.

**Figure S3.**
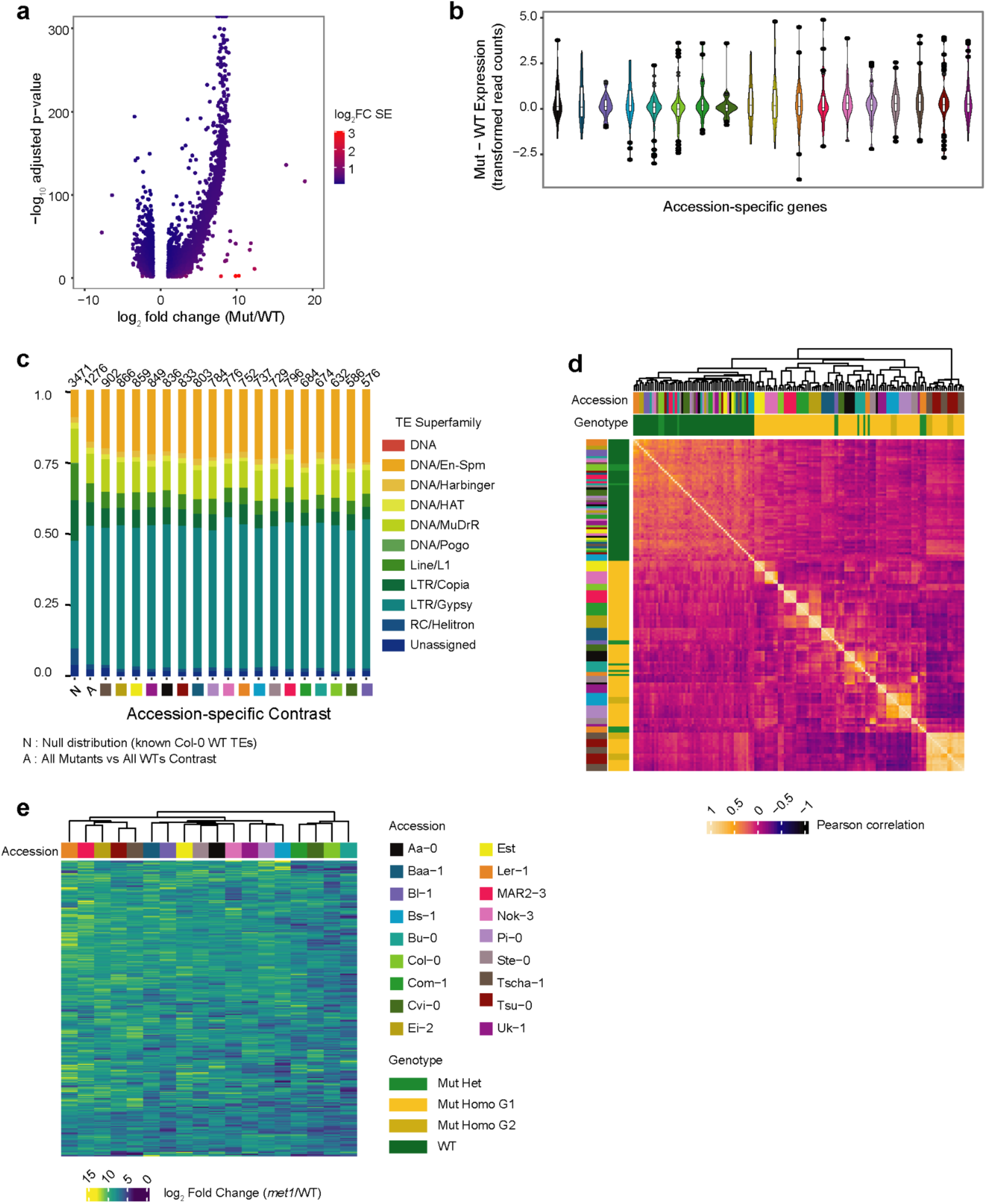
Accession-specific variation of differentially expressed genes (DEGs) in *met1* mutants. **(a)** Volcano plot colored by standard error (SE) of log_2_ fold change (FC) in all-mutants-against-all-wild-type DEGs. **(b)** Thirty random genes examined for accession-specific variation in expression changes (measured as transformed read counts). **(c)** Distribution of TE superfamilies in TE-DEGs across 19 contrasts, and a null distribution of all TE genes (denoted by ’N’). **(d)** Correlation between 291 universal DEGs across 158 RNA-seq libraries (104 *met1* mutant and 54 wild-type samples). **(e)** Heatmap showing log_2_ fold change in expression (Mut/WT) of 276 universal TE-DEGs. Color code of accessions in (b), (c) and (d) follows the same legend shown in (e).

**Figure S4.**
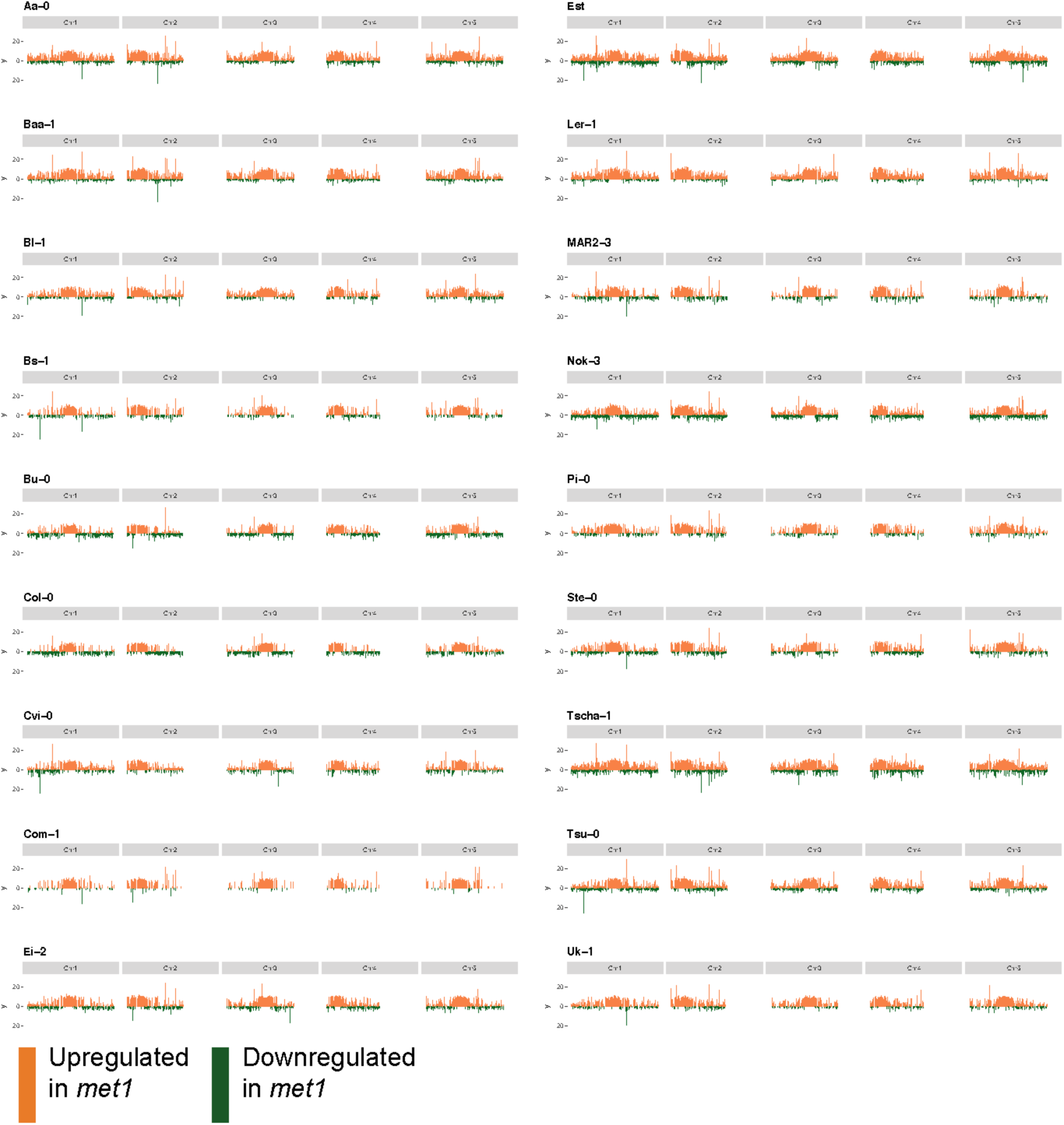
Accession-specific variation in the chromosomal distribution of up- and down-regulated DEGs.

**Figure S5.**
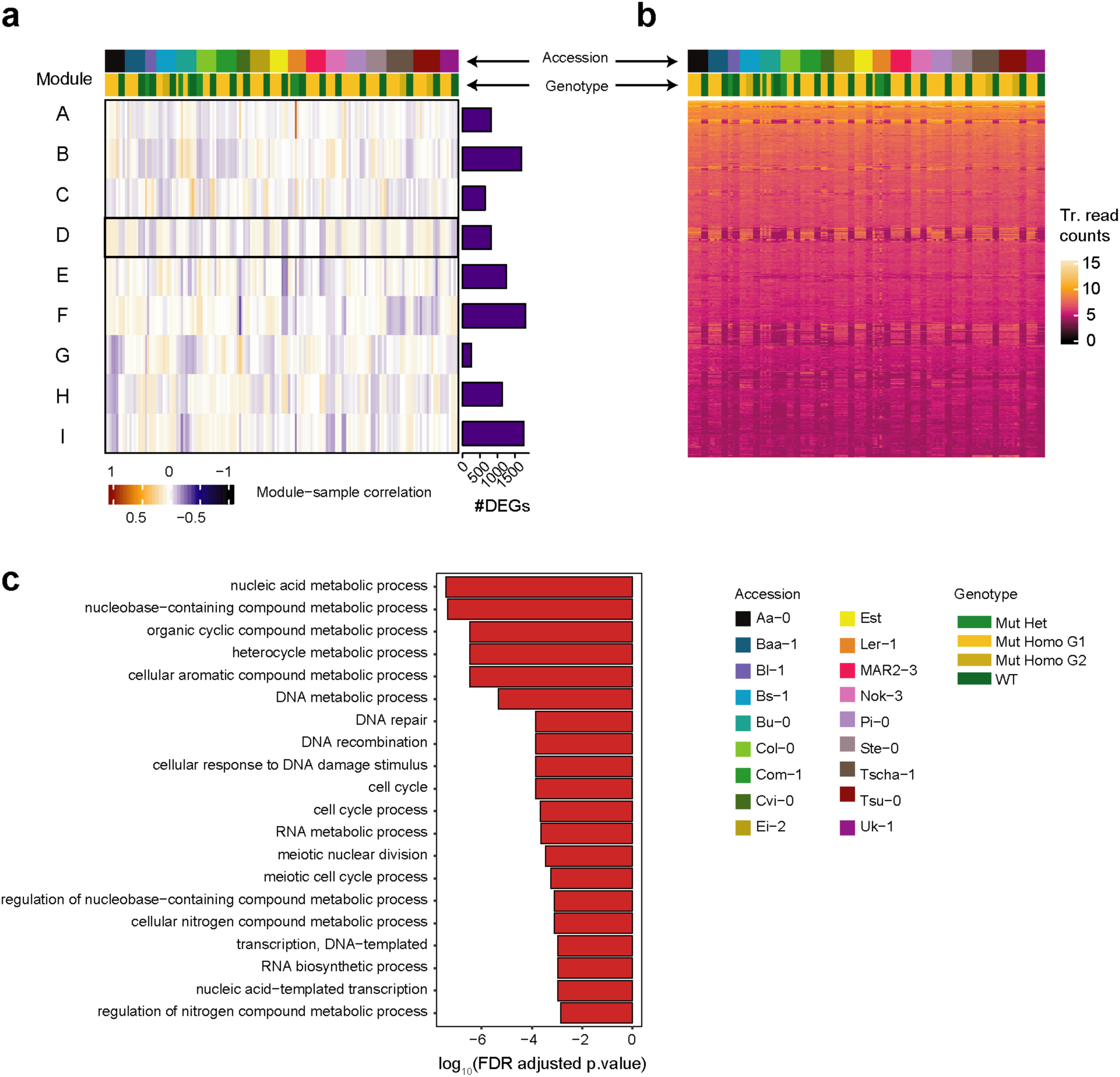
Gene network analysis on unique Non-TE-DEGs across all 19 contrasts. **(a)** Heatmap showing module-sample correlation levels of 10,151 unique Non-TE-DEGs from 19 contrasts, across 158 samples. The rows represent 9 modules (labeled A - I) based on weighted gene co-expression network analysis, with module ’D’ highlighted due to its high association with sample genotype. The marginal barplots along the vertical axis indicate the number of genes in each module. **(b)** Heatmap of transformed read counts across 158 RNA-seq libraries for 814 genes in module ’D’. (c) Results of the top 20 most significant Gene Ontology enrichment terms for 814 genes in module ’D’.

**Figure S6.**
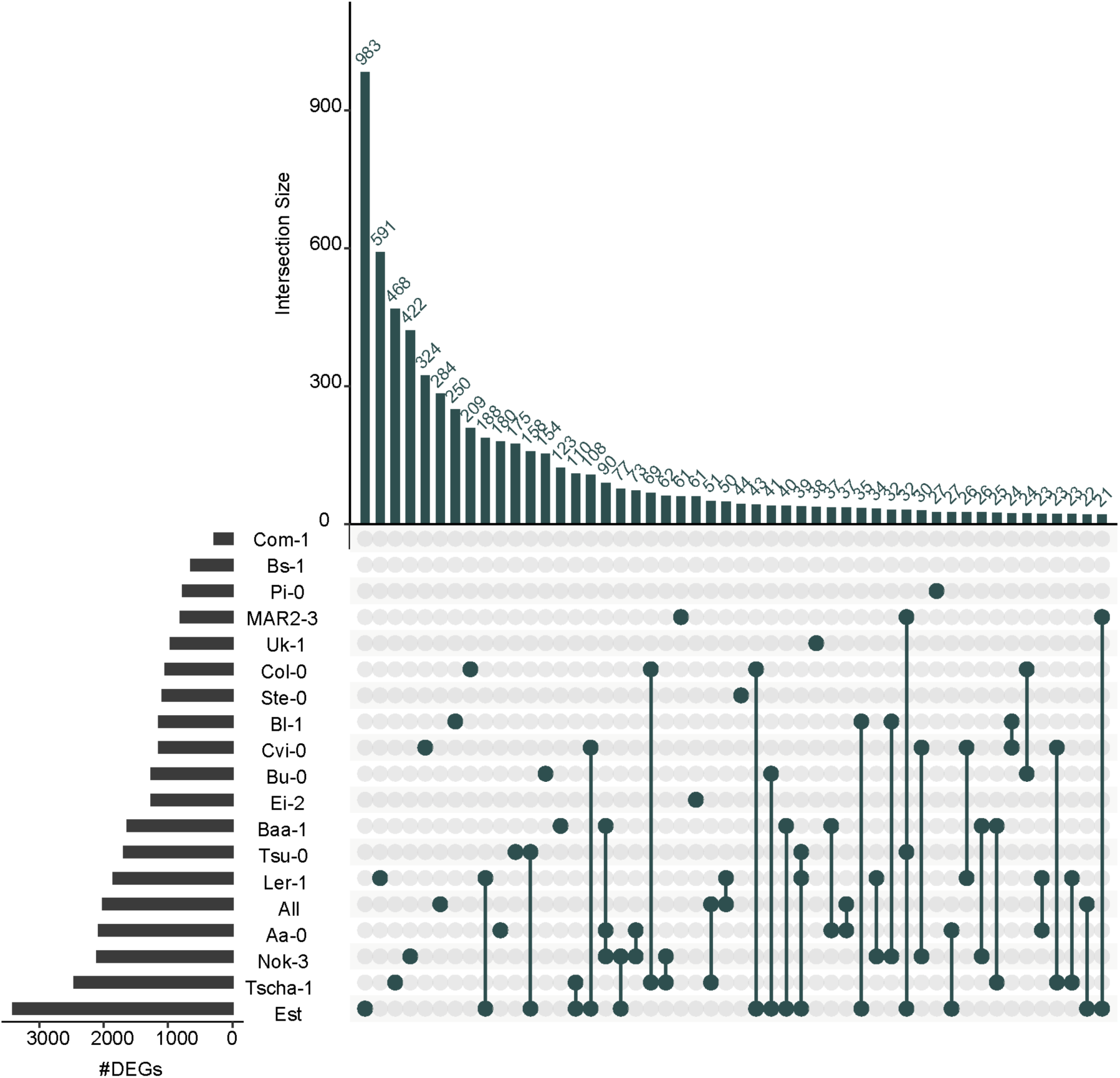
Comparisons and intersections between accession-specific DEGs. Upset plot of the top 50 all-pairwise intersections between Non-TE-DEGs in 19 accession-specific contrasts.

**Figure S7.**
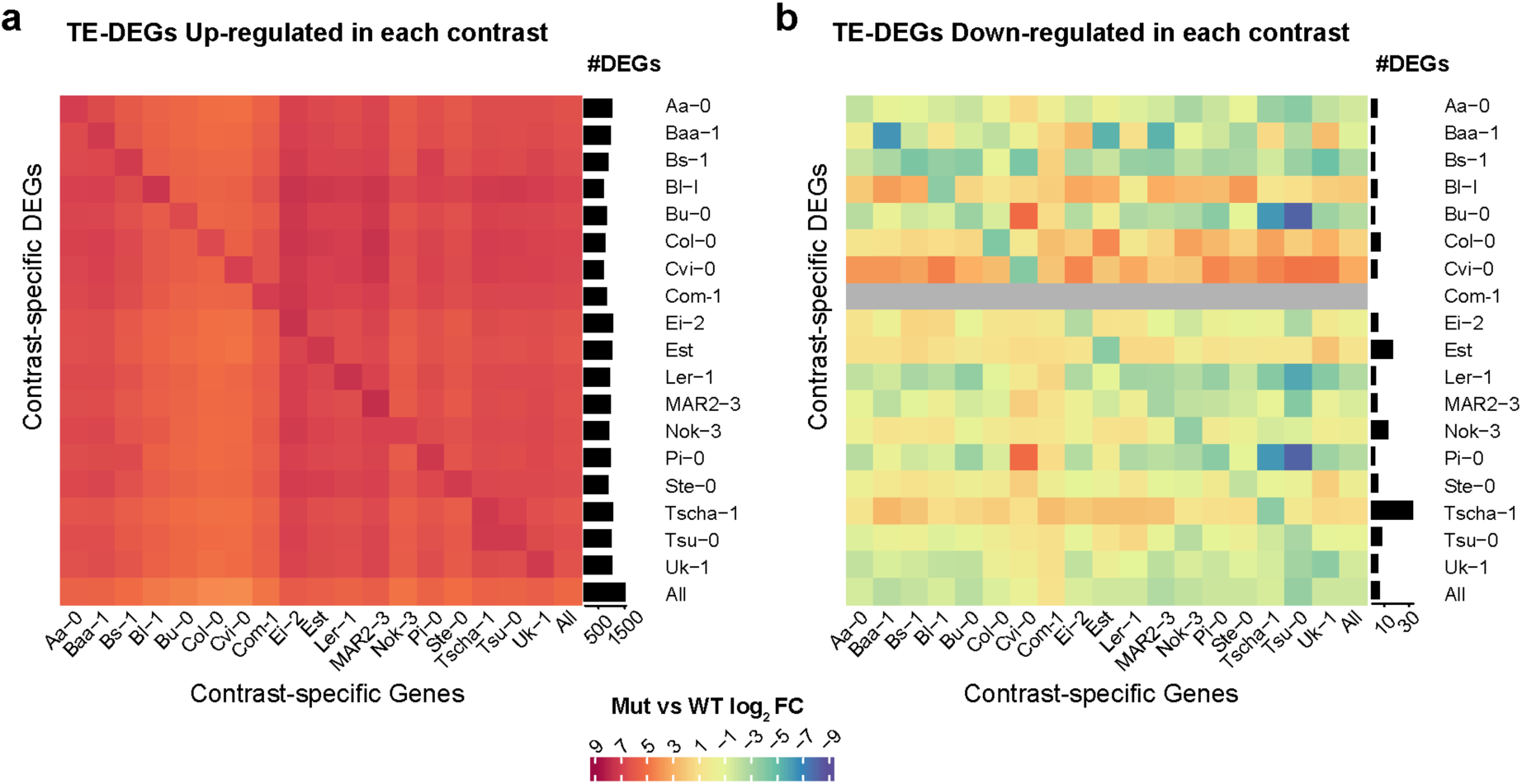
Quantitative comparisons between TE-DEGs across accession-specific contrasts. Heatmaps showing average log_2_ fold change (Mut/WT) for all TE-DEGs which are **(e)** upregulated and **(f)** downregulated in each accession-specific contrast, measured for the same genes across all other contrasts. Marginal barplots indicate the number of TE-DEGs in every accession-specific contrast. Com-1 did not exhibit any downregulated TE-DEGs and therefore the corresponding heatmap tiles in (b) are colored grey.

**Figure S8.**
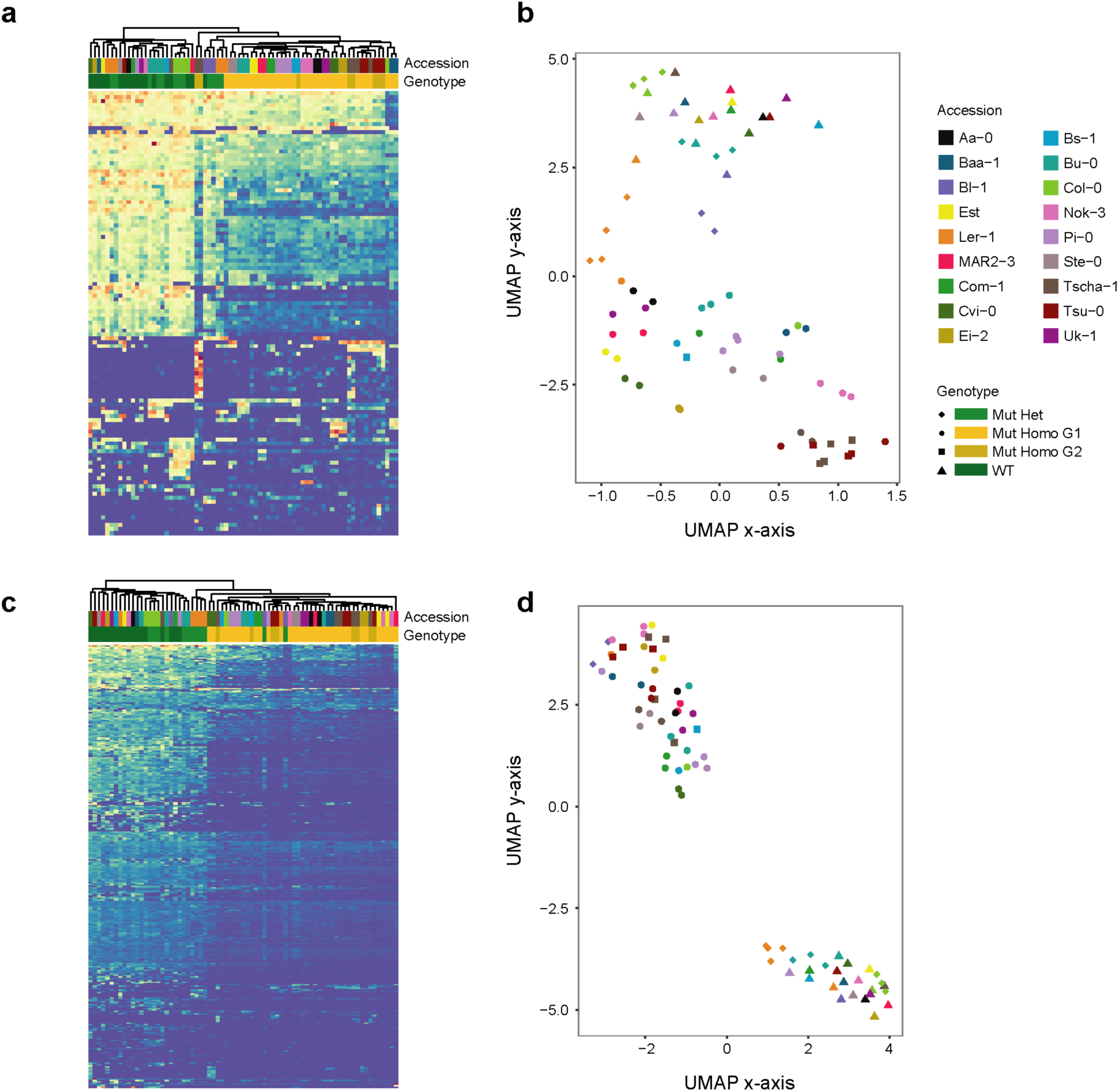
Differential methylation in non-CG contexts in *met1* mutants and wild-type individuals. **(a)** Heatmap and **(b)** UMAP visualization of CHG methylation levels in 73 samples (55 mutants and 18 wild-type plants) across 114 CHG-DMRs (from a total of 350 CHG-DMRs). **(c)** Heatmap and **(d)** UMAP visualization of CHH methylation levels in 73 samples (55 mutants and 18 wild-type plants) across 334 CHG-DMRs (from a total of 1,023 CHG-DMRs).

**Figure S9.**
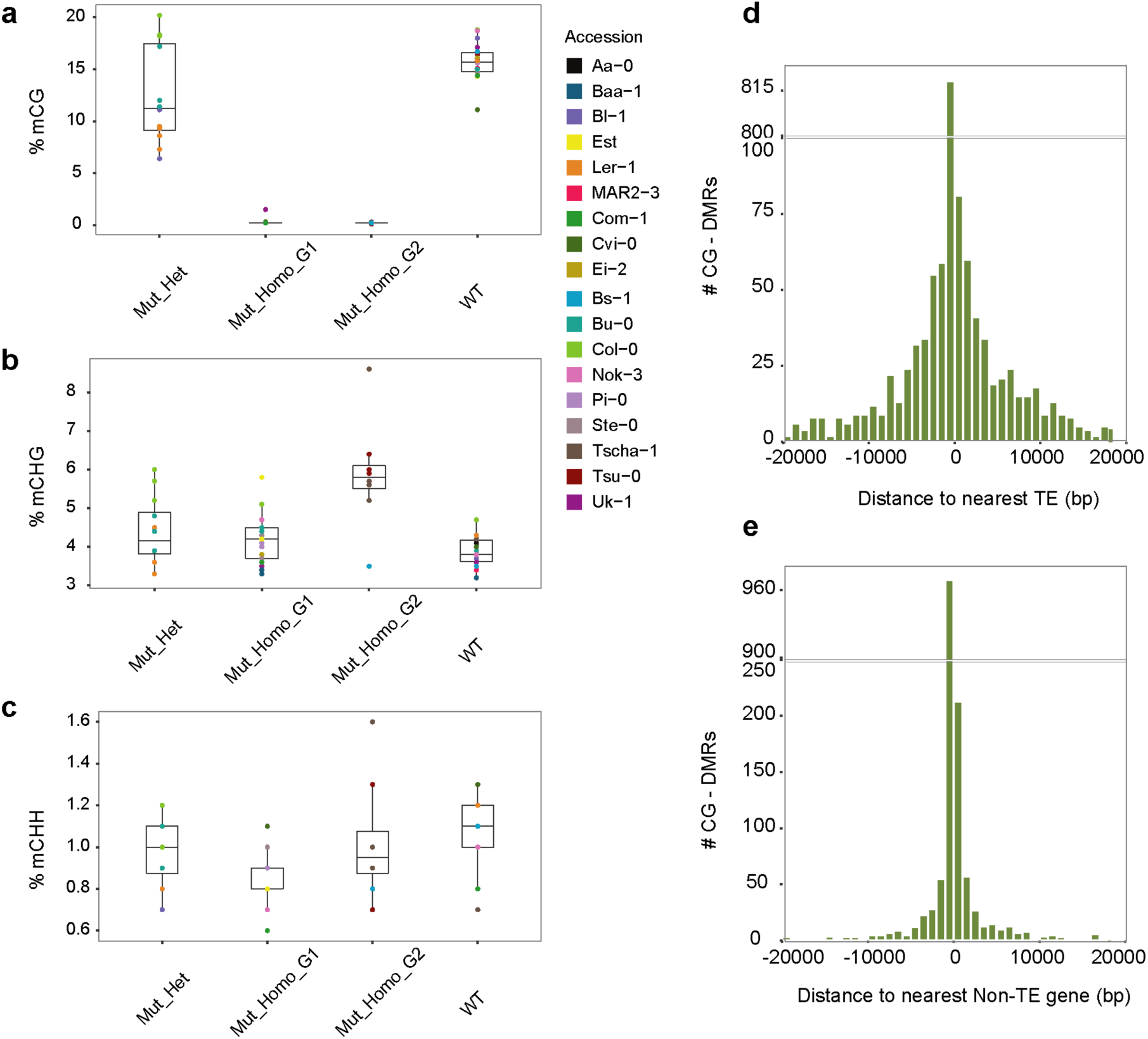
Altered genome-wide methylation levels in *met1* mutants compared to wild-type individuals. Genome-wide methylation levels of 73 samples (55 *met1* mutants and 18 wild-type plants) in the **(a)** CG, **(b)** CHG and **(c)** CHH contexts. Histogram showing distance of CG-DMRs to **(d)** nearest TE and **(e)** nearest Non-TE gene.

**Figure S10.**
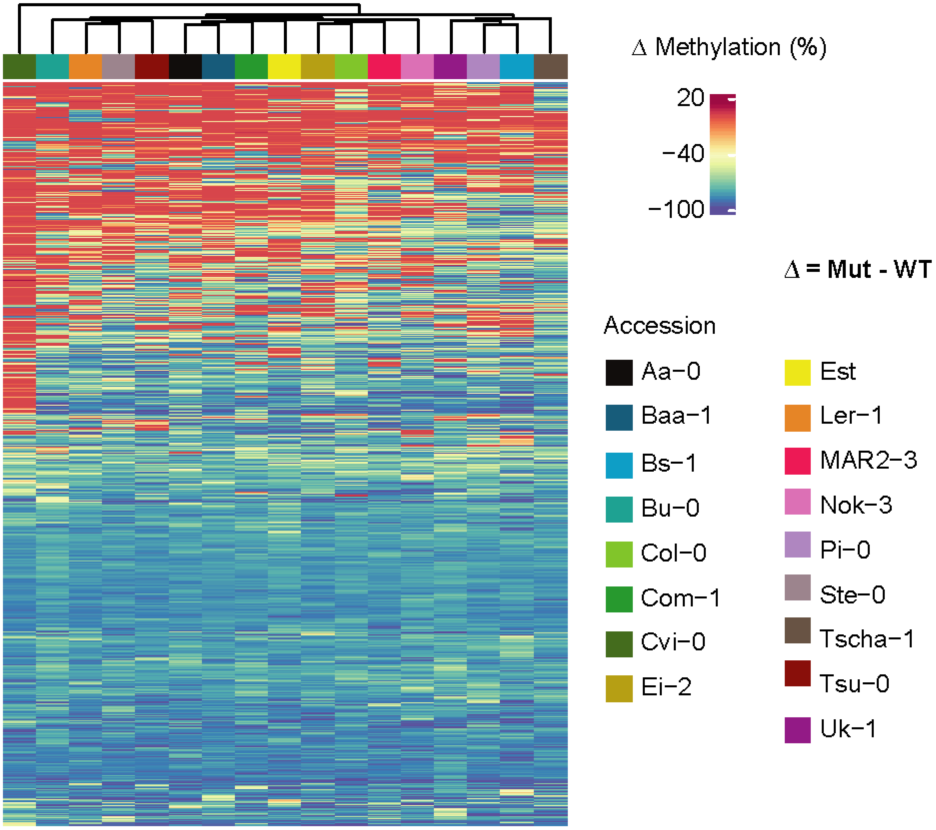
Heatmap of differences in CG methylation between first generation homozygotes and wild-type samples for 17 accessions across 749 CG DMRs.

**Figure S11.**
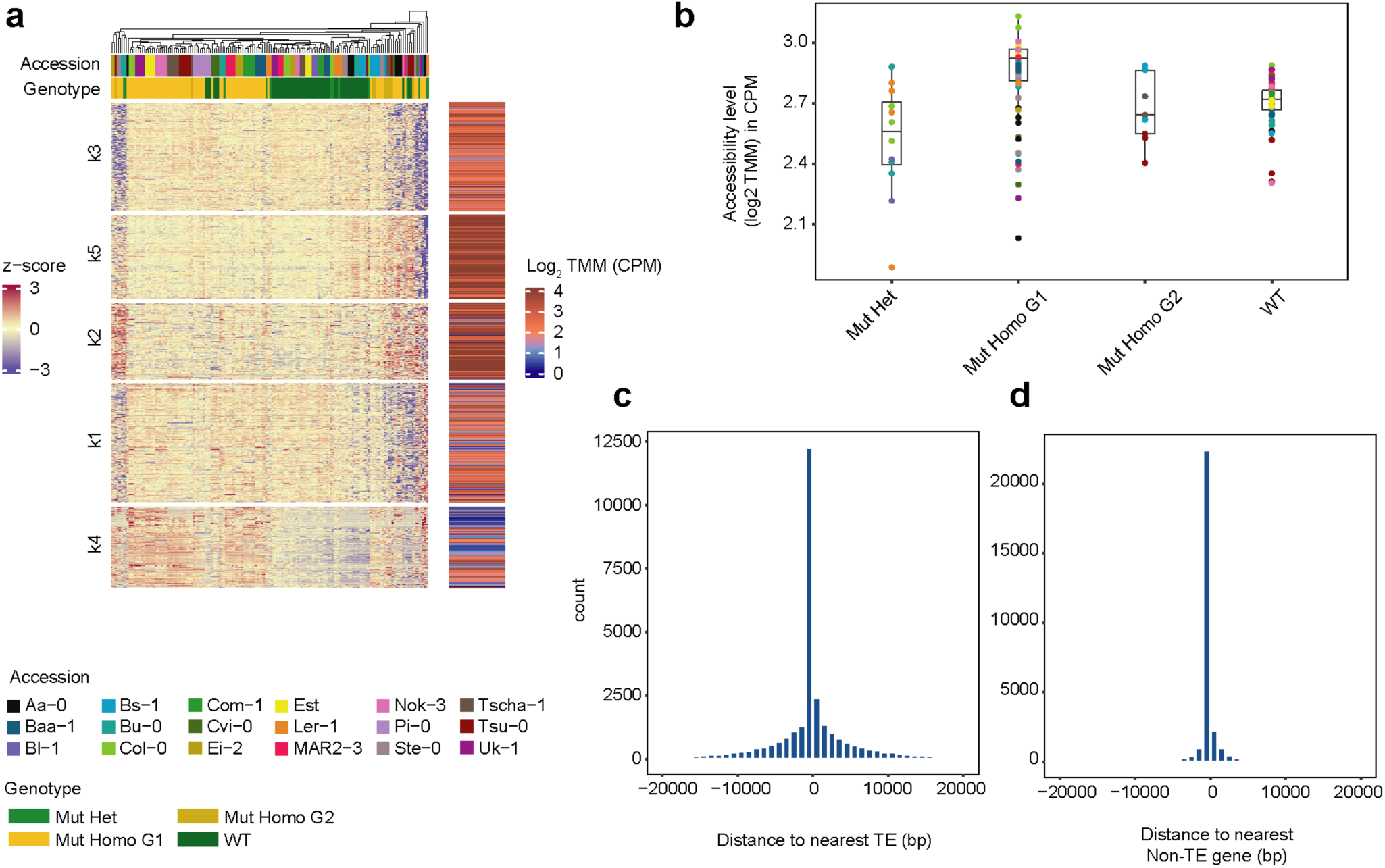
Altered genome-wide chromatin accessibility in *met1* mutants compared to wild-type individuals. **(a)** Heatmap of 31,295 z-scaled dACRs across 158 ATAC-seq libraries (104 *met1* mutant and 54 wild-type samples) and **(b)** histograms showing distance of dACRs to nearest TE and **(c)** nearest Non-TE protein coding gene.

**Figure S12.**
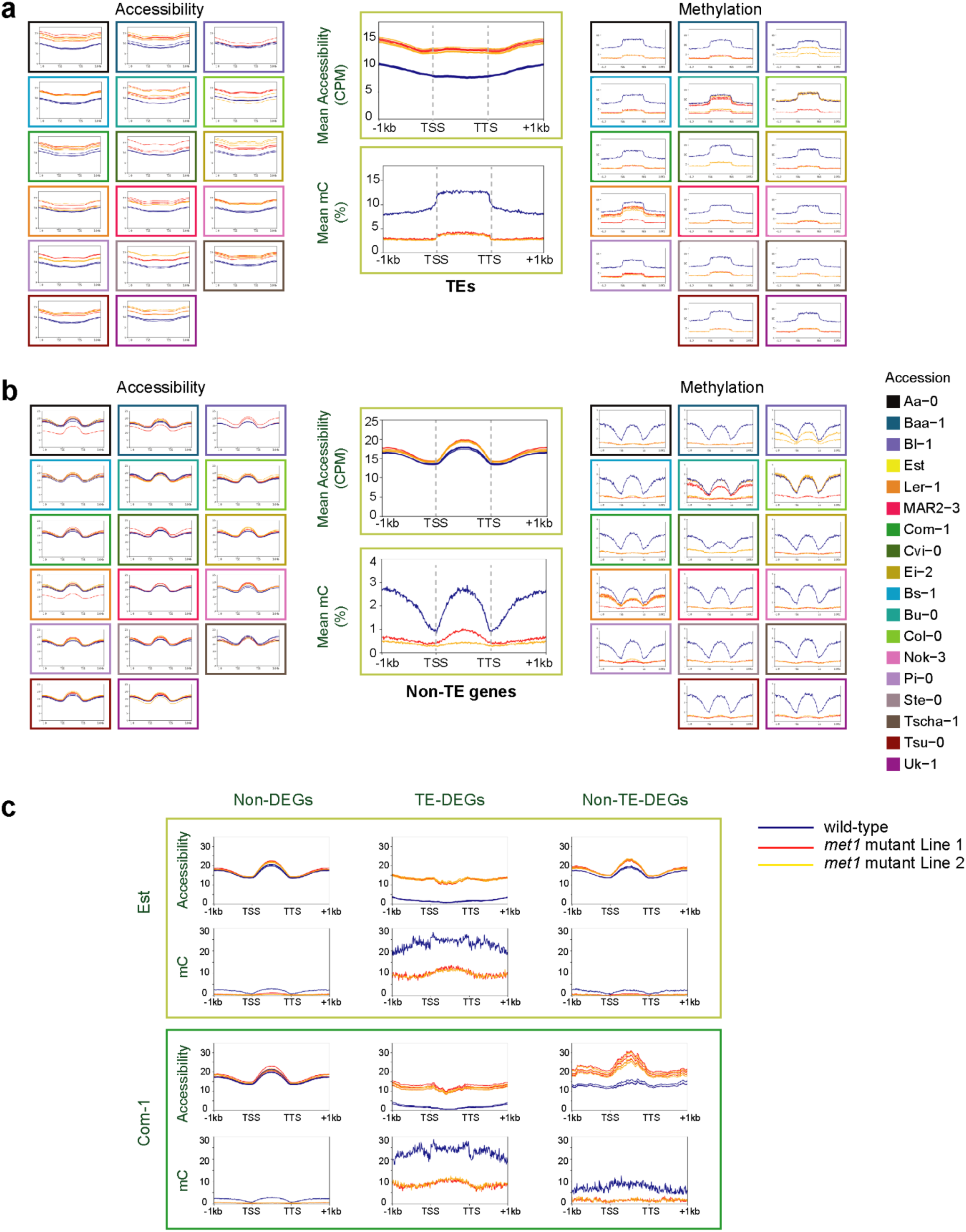
*met1* mutants have more accessible chromatin and are hypomethylated over TEs and genes. Metaplots of mean chromatin accessibility across all **(a)** TAIR10 TEs and **(b)** Non-TE protein-coding genes in 18 accessions. Boxes are color-coded by accession, and data are colored by genotype. **(c)** Metaplots of chromatin accessibility and methylation levels for Est and Com-1, across Non-DEGs, TE-DEGs and Non-TE-DEGs. TSS and TTS denote transcription start site and transcription termination site respectively. Methylation levels are represented as % cytosine methylation (all-contexts) and accessibility levels are represented as TMM normalized values in counts per million (CPM).

**Figure S13.**
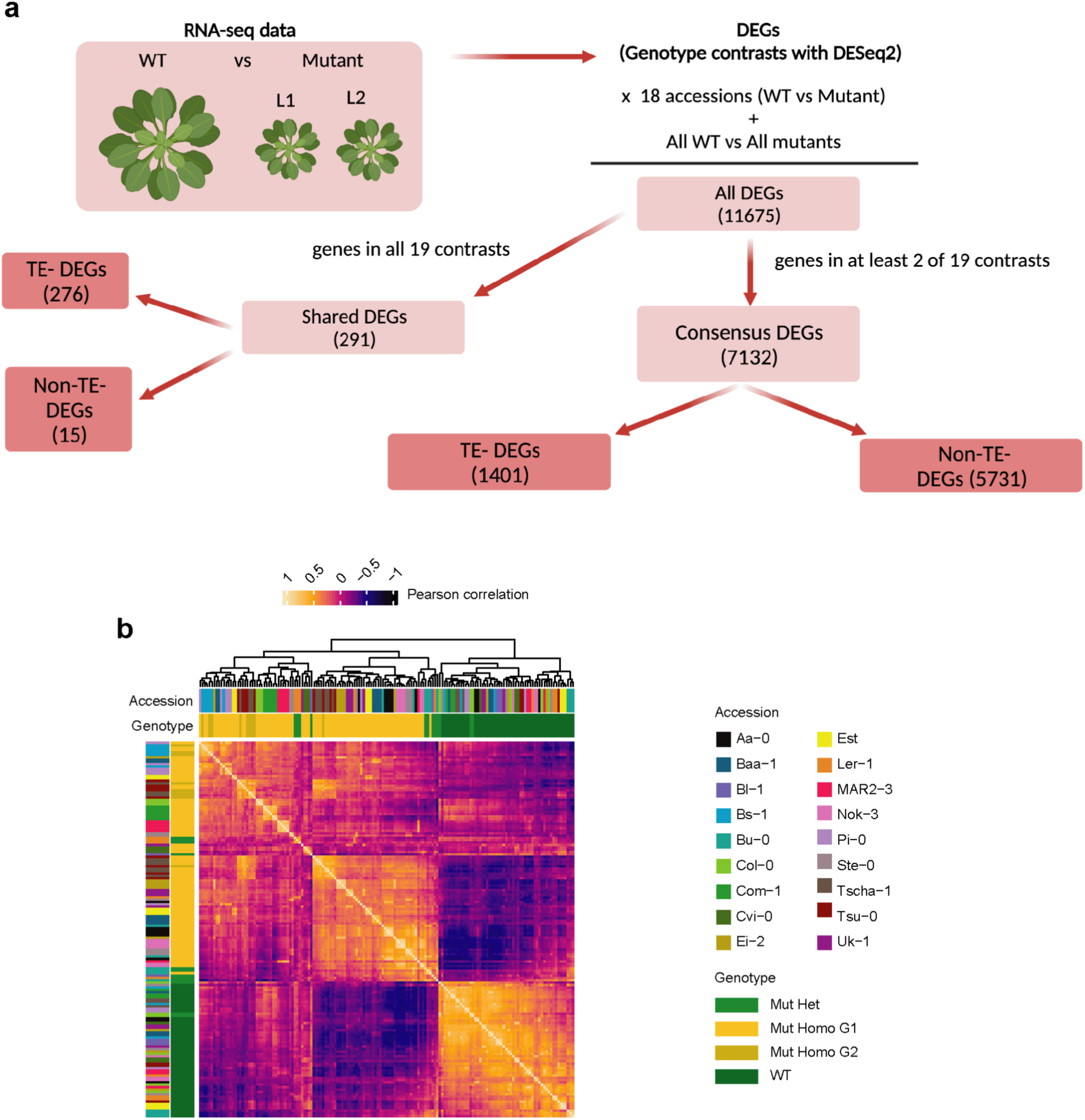
**(a) Diagram of generating consensus DEGs from RNA-seq data. (b)** Correlation between 7132 consensus DEGs across 158 RNA-seq libraries (104 *met1* mutant and 54 wild-type samples). Genotypes represented are wildtypes (’WT’), heterozygous *met1* mutants (’Mut Het’), first generation homozygous *met1* mutants (’Mut Homo G1’) and second generation homozygous *met1* mutants (’Mut Homo G2’).

**Figure S14.**
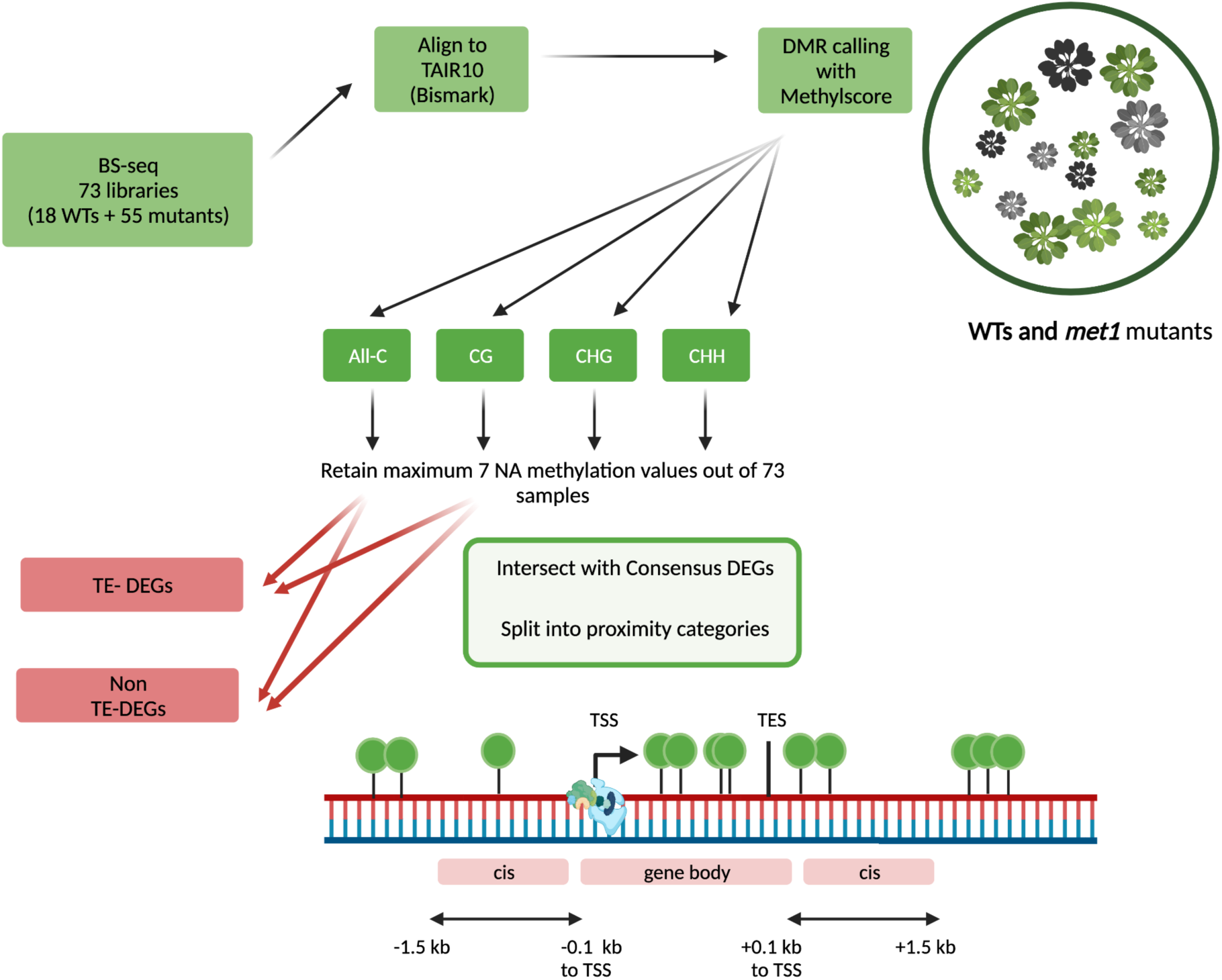
Diagram of generating DMRs from BS-seq data and intersections with consensus DEGs.

**Figure S15.**
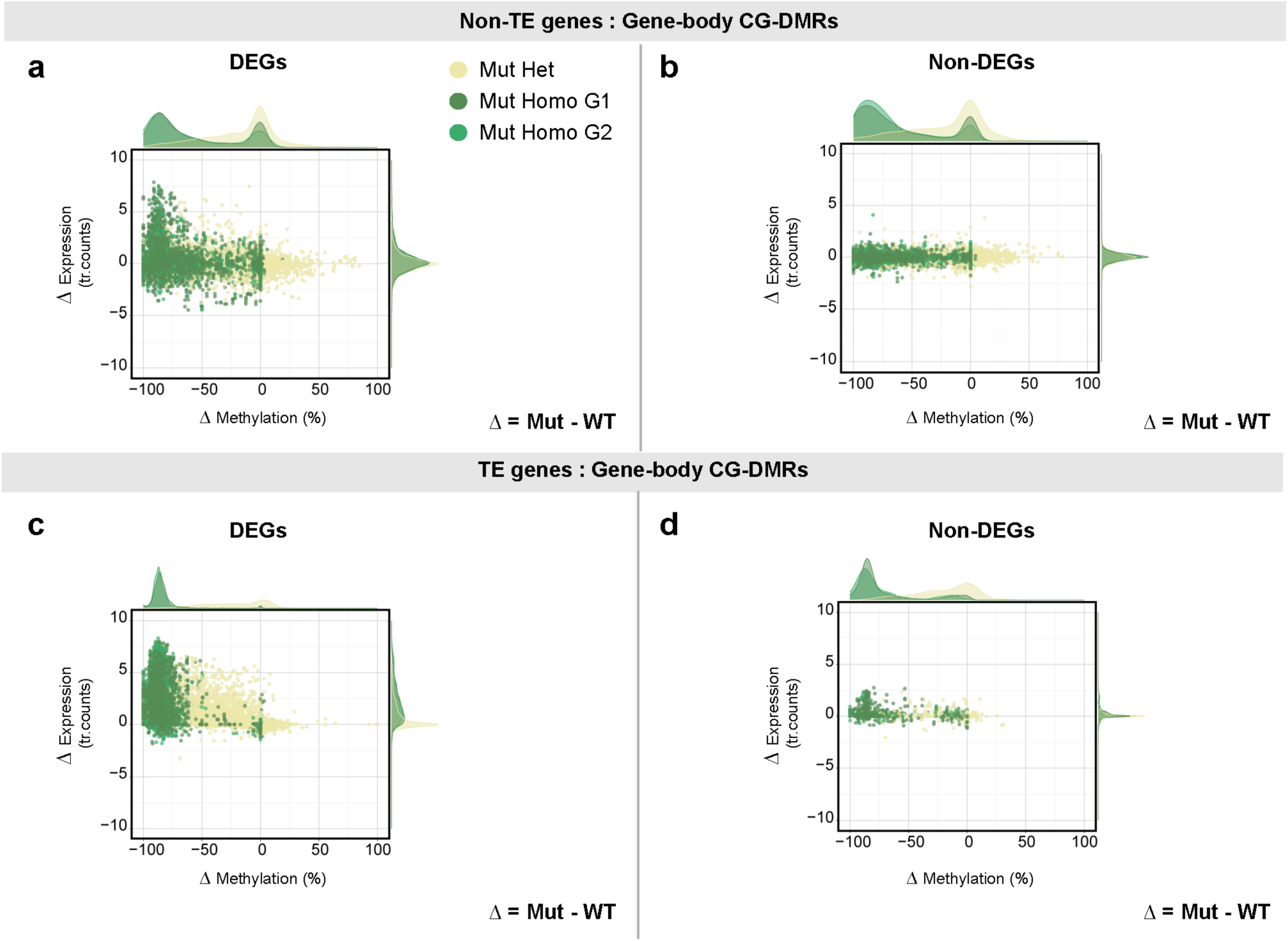
DMRs in gene-bodies of Non-TE genes (a-b) and TE genes (c-d). Scatter plots showing differences in CG methylation between *met1* mutants and wild-type plants against differences in gene expression. Dots in the scatter plot are colored by genotype of *met1* mutants; wildtypes (’WT’), heterozygous *met1* mutants (’Mut Het’), first generation homozygous *met1* mutants (’Mut Homo G1’) and second generation homozygous *met1* mutants (’Mut Homo G2’) with x- and y-axis density distributions of each genotype. Expression levels are represented as transformed read counts (Methods) and methylation levels are represented as % CG methylation.

**Figure S16.**
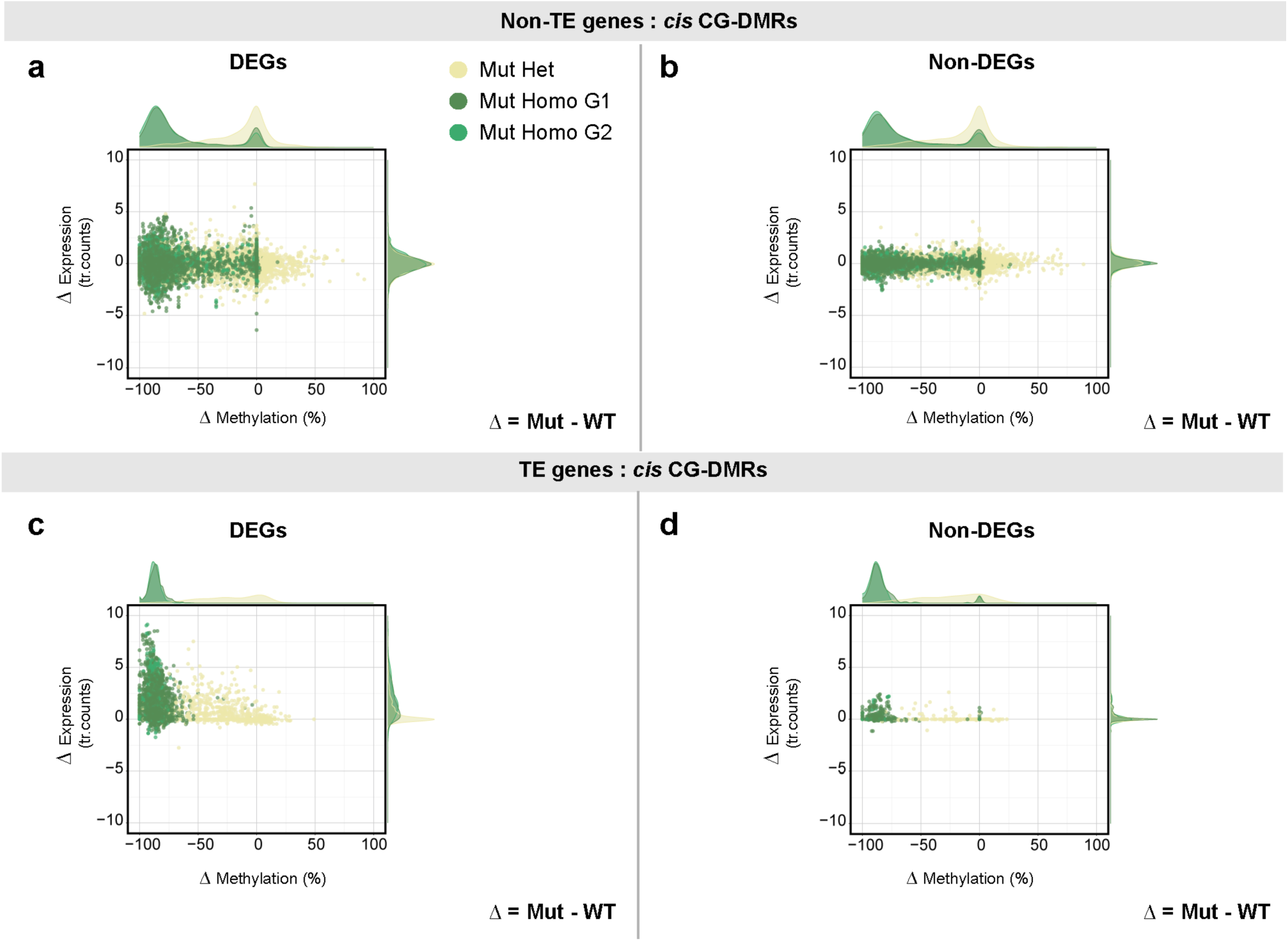
DMRs in *cis* to Non-TE genes (a-b) and TE genes (c-d). Scatter plots showing differences in CG methylation between *met1* mutants and wild type plants against differences in gene expression. Dots in the scatter plot are colored by genotype of *met1* mutants; wildtypes (’WT’), heterozygous *met1* mutants (’Mut Het’), first generation homozygous *met1* mutants (’Mut Homo G1’) and second generation homozygous *met1* mutants (’Mut Homo G2’) with x- and y-axis density distributions of each genotype. Expression levels are represented as transformed read counts and methylation levels are represented as % CG methylation.

**Figure S17.**
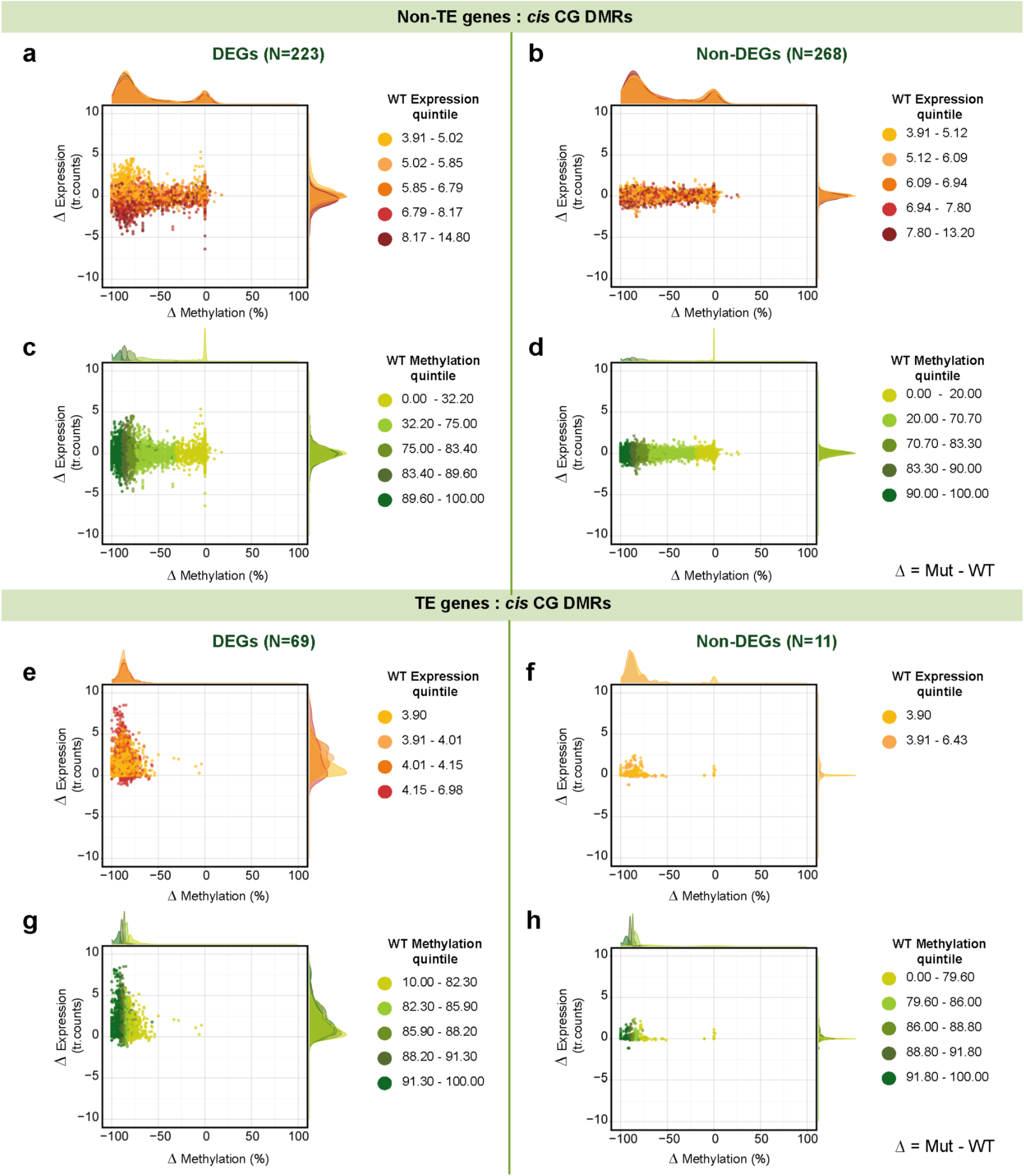
CG-DMRs in *cis* to Non-TE-DEGs (a-d) and TE-DEGs (e-h). Scatter plots showing differences in CG methylation between *met1* mutants and wild-type plants against difference in gene expression. Dots in the scatter plot are colored by wild-type expression quintiles **(a,b,e,f)** and wild-type methylation quintiles **(c,d,g,h)** with x- and y-axis density distributions of each expression/methylation quintile. Expression levels are represented as transformed read counts and methylation levels are represented as %CG methylation in CG-DMRs.

**Figure S18.**
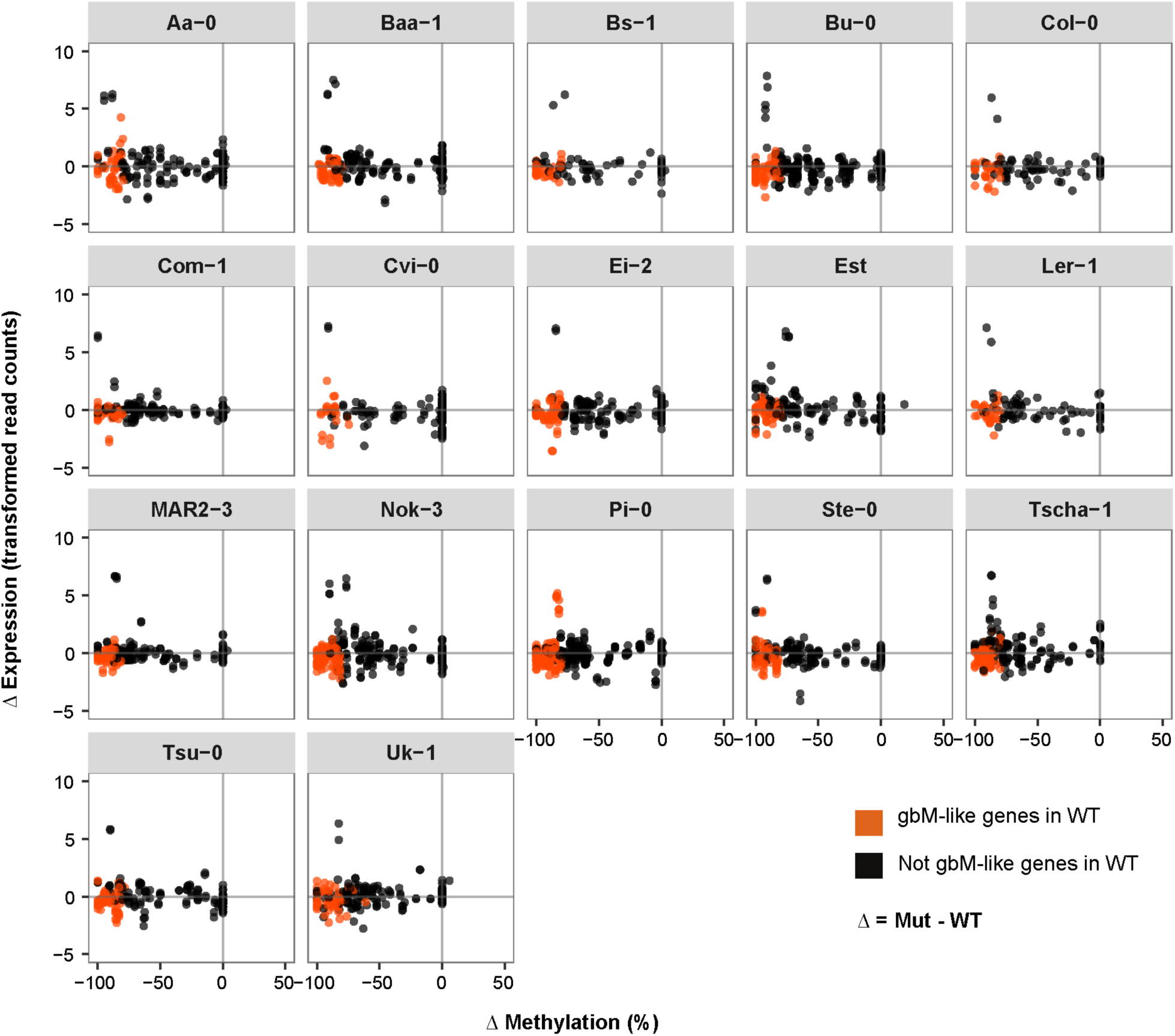
Methylation changes and associated gene expression changes for 91 gbM-like genes across 17 accessions. Orange colored dots represent gbM-like genes and black dots represent the same genes which are not gbM-like in other accessions. ΔMethylation represents *met1* - wild-type methylation, measured in % CG methylation level. ΔExpression represents *met1* - wild-type gene expression levels, measured in transformed read counts.

**Figure S19.**
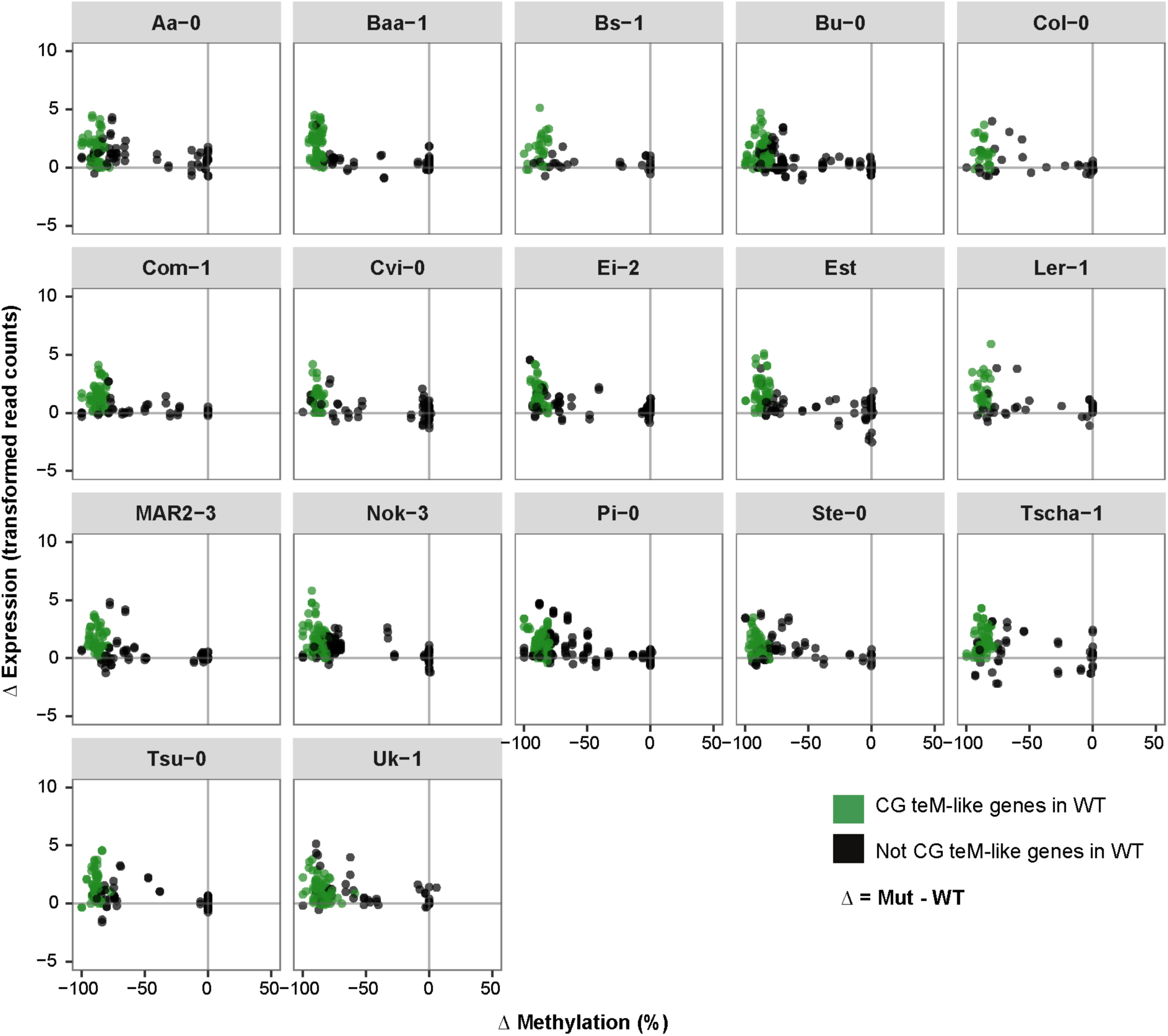
Methylation changes and associated gene expression changes for 57 CG teM-like genes across 17 accessions. Green colored dots represent CG teM-like genes and black dots represent the same genes which are not CG teM-like in other accessions. ΔMethylation represents *met1* - wild-type methylation, measured in % CG methylation level. ΔExpression represents *met1* - wild-type gene expression levels, measured in transformed read counts.

**Figure S20.**
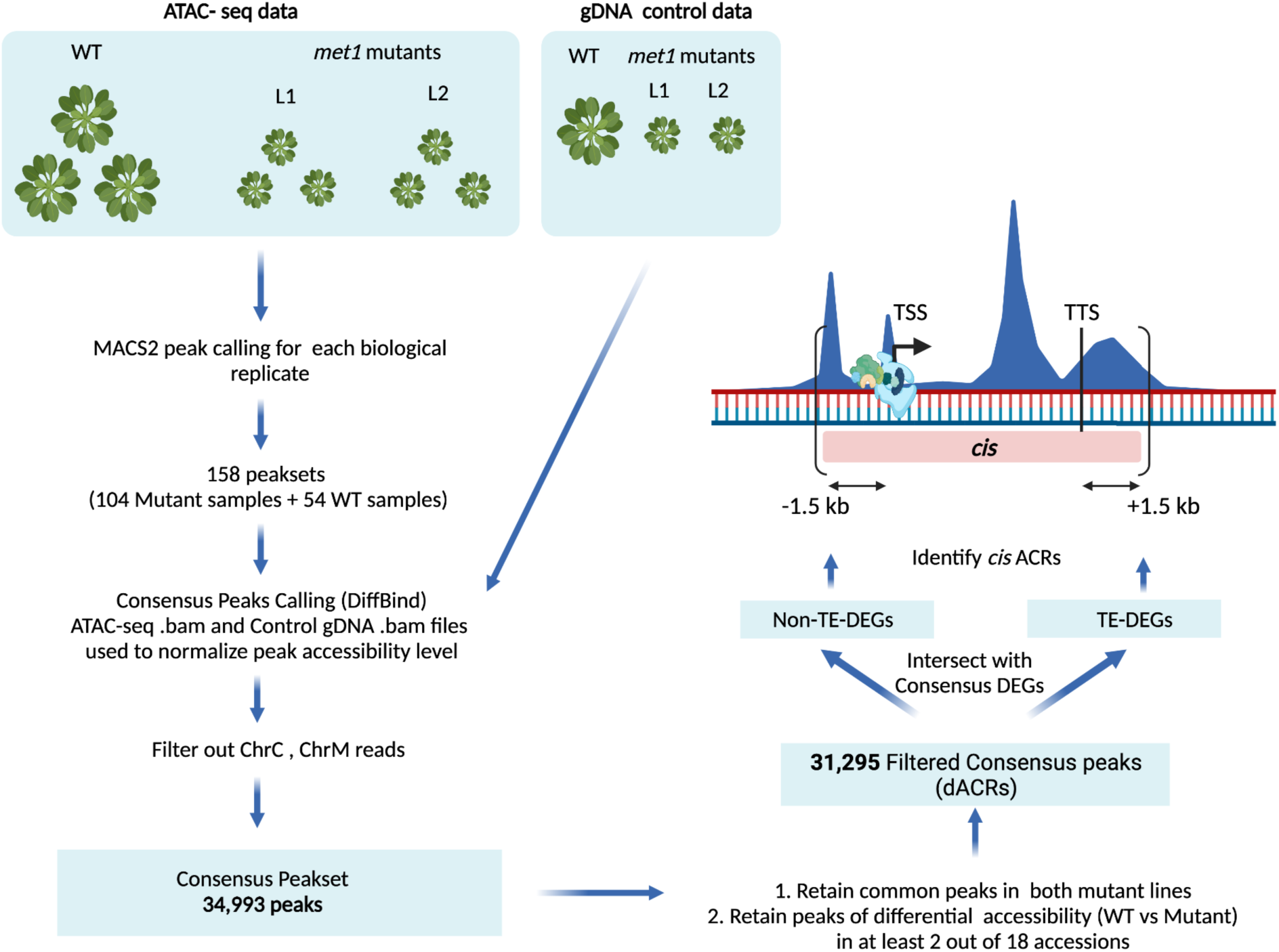
Diagram of ATAC-seq processing, generation of filtered consensus peaks (dACRs) and intersections with DEGs.

**Figure S21.**
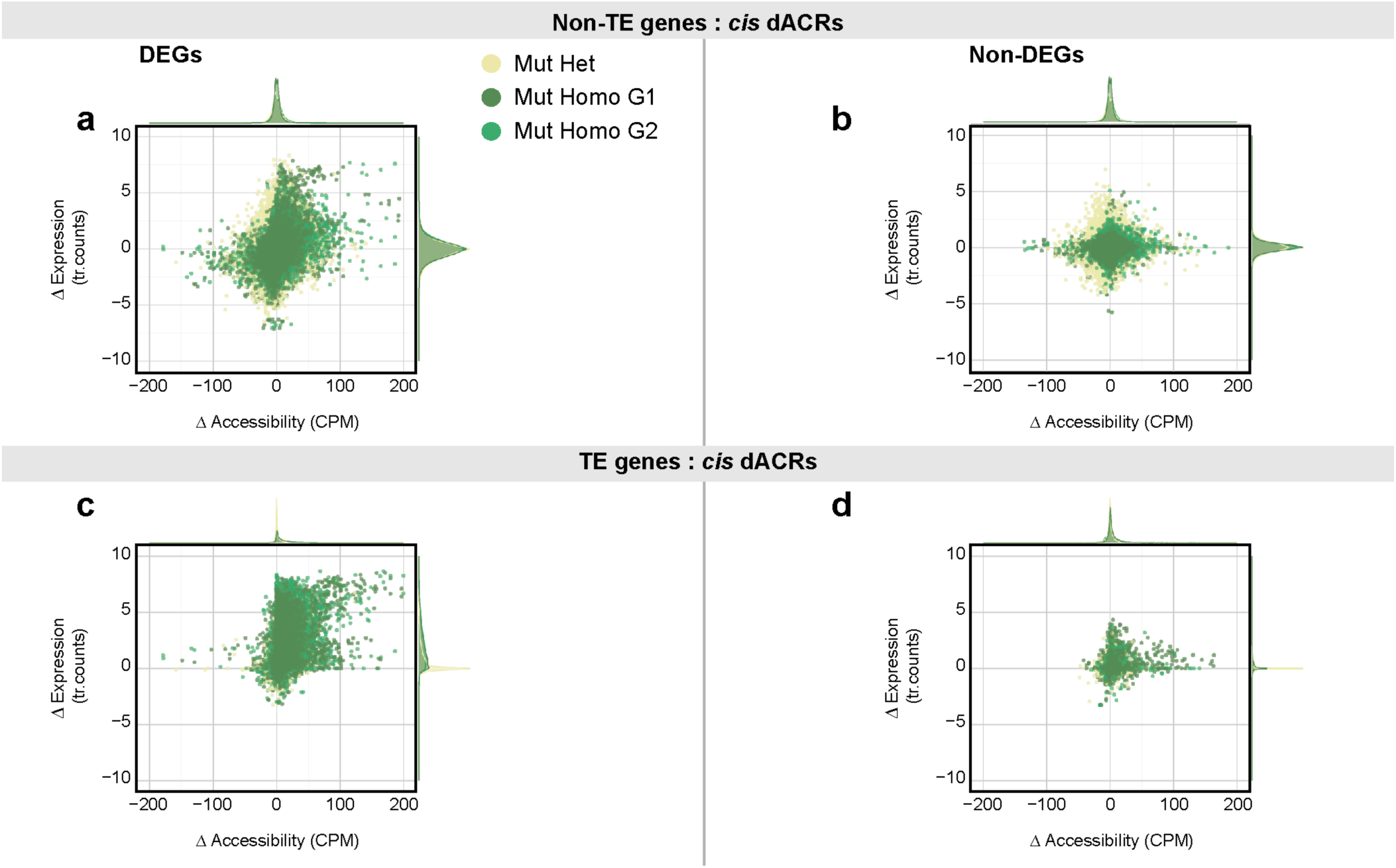
dACRs in *cis* to TE genes (c,d) and Non-TE genes (a,b). Scatter plots showing difference in Chromatin accessibility between *met1* mutants and wild-type plants against differences in gene expression. Dots in the scatter plot are colored by genotype of *met1* mutants; wildtypes (’WT’), heterozygous *met1* mutants (’Mut Het’), first generation homozygous *met1* mutants (’Mut Homo G1’) and second generation homozygous *met1* mutants (’Mut Homo G2’) with x- and y-axis density distributions of each genotype. Expression levels are represented as transformed read counts and accessibility levels are represented as TMM normalized values in counts per million (CPM).

**Figure S22.**
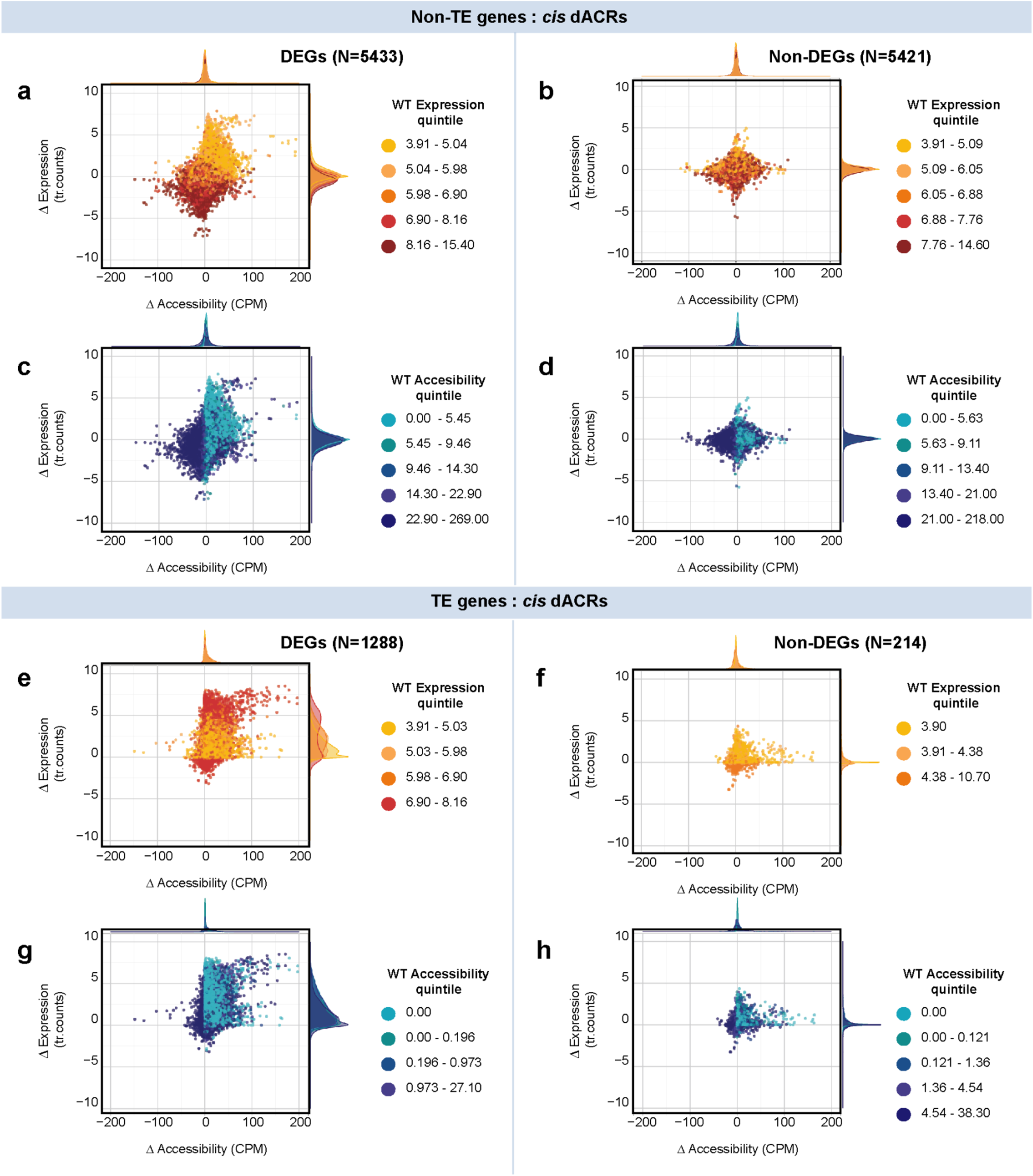
dACRs in *cis* to TE-DEGs (a-d) and Non-TE-DEGs (e-h). Scatter plots showing differences in chromatin accessibility between *met1* mutants and wild-type plants against differences in gene expression. Dots in the scatter plot are colored by wild-type expression quintiles **(a,b,e,f)** and wild-type accessibility quintiles **(c,d,g,h)** with x- and y-axis density distributions of each expression/accessibility quintile. Expression levels are represented as transformed read counts and accessibility levels are represented as TMM normalized values in counts per million (CPM).

**Figure S23.**
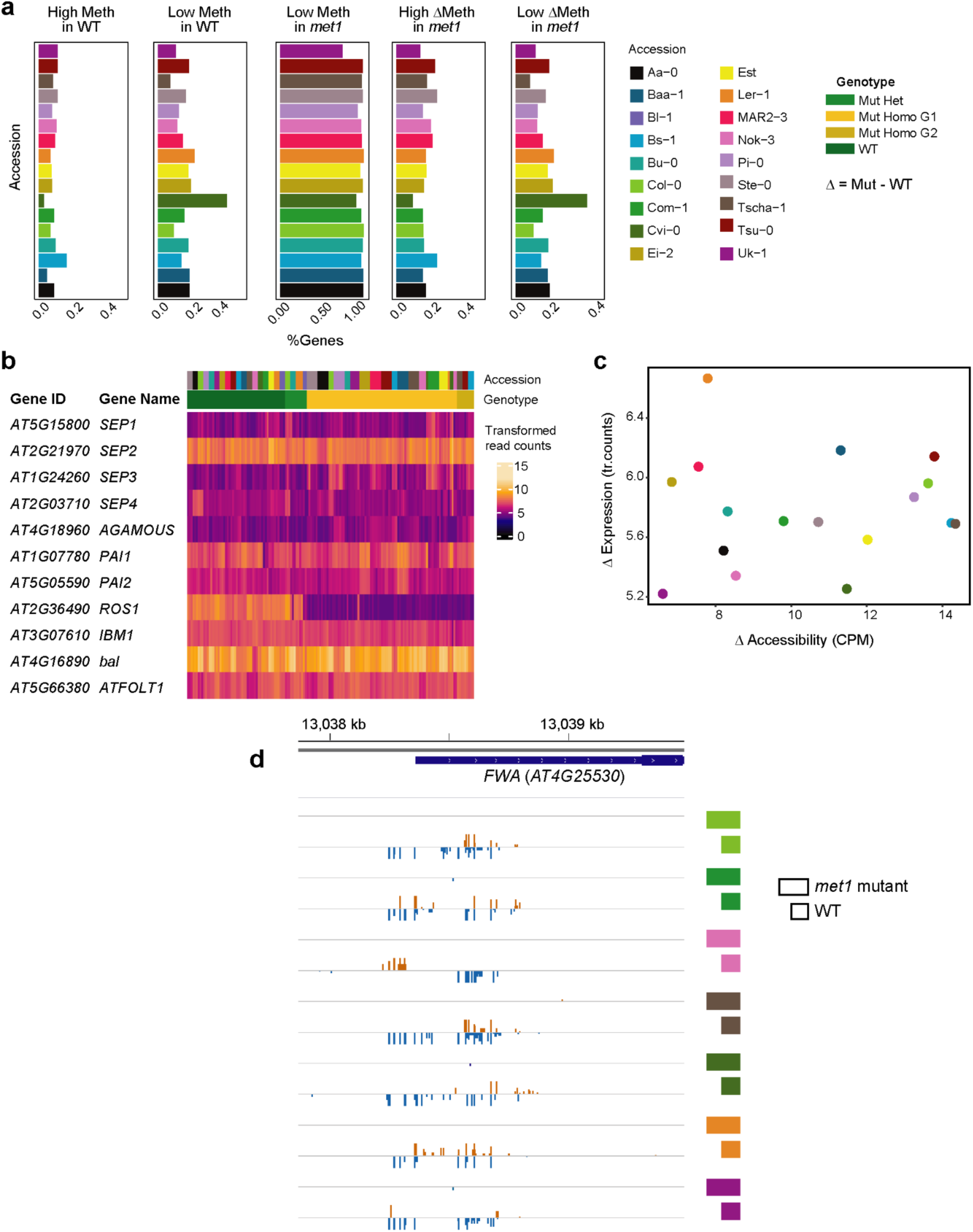
Variable epigenetic states of Non-TE genes across accessions. **(a)** Five panels showing fraction of genes in 17 accessions with (i) high CG methylation in wild type, (ii) low CG methylation in wild type, (iii) low CG methylation in *met1*, (iv) large methylation change in *met1*, (v) limited methylation change in *met1.* Colors represent different accessions. Low methylation is defined as CG methylation <= 10%, and high methylation is defined as CG methylation >= 90%. **(b)** Heatmap of transformed read counts at 11 epialleles across 158 RNA-seq libraries. The libraries are colored by accession-of-origin and genotype. **(c)** Scatterplot showing relationship between changes in accessibility and expression at the *FWA* locus. Accessibility is measured in counts per million (CPM) and expression is measured by transformed read counts. **(d)** Genome browser screenshot of methylated cytosines (all contexts) at the *FWA* locus for *met1* mutants and wild-type plants of seven accessions.

**Figure S24.**
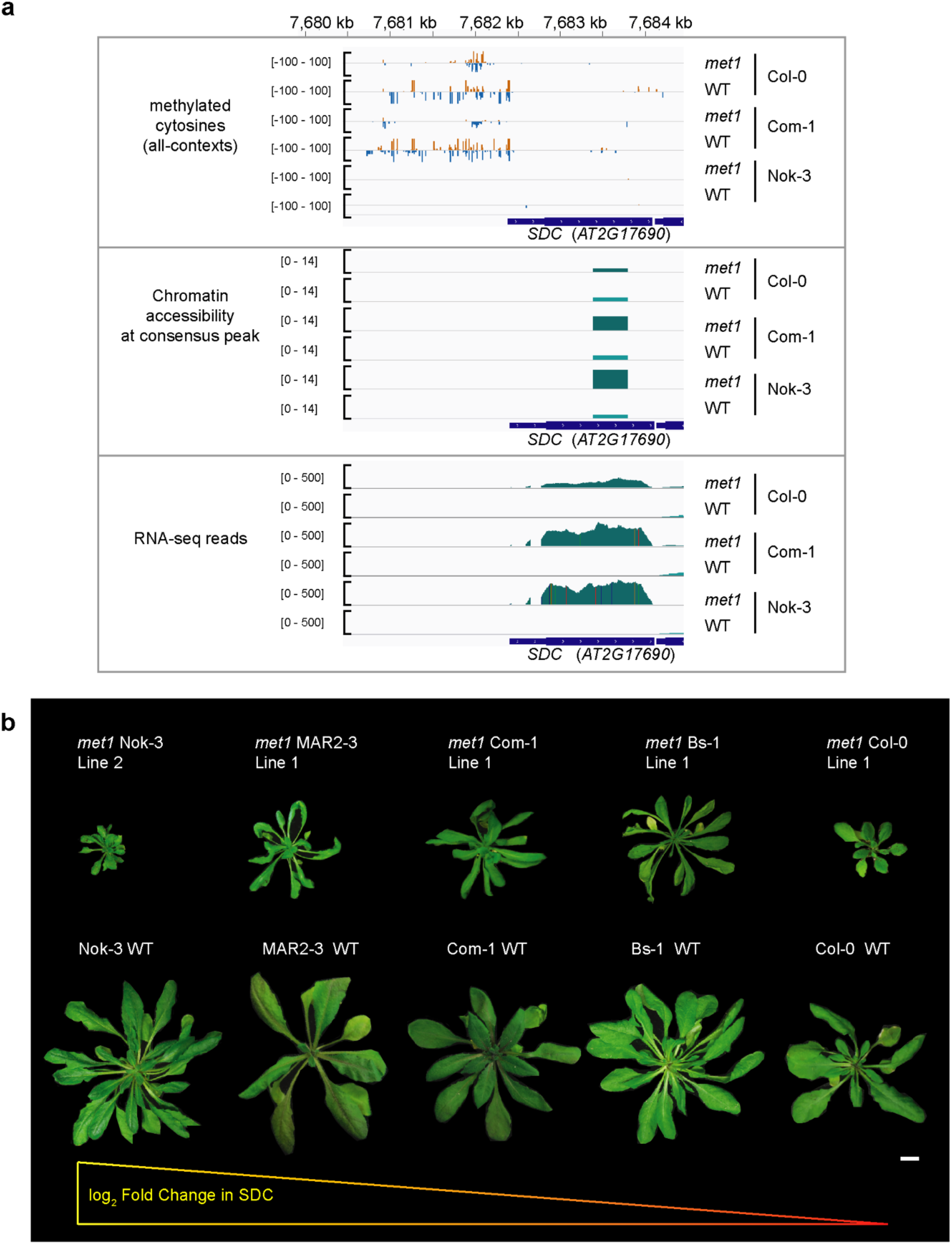
Epigenetic landscape at the *SDC* locus. **(a)** Genome browser screenshot of methylated cytosines, chromatin accessibility and RNA-seq read count in *met1* mutants and wildtypes of three accessions, Col-0, Com-1 and Nok-3. Chromatin accessibility is shown here as a region spanning 200 bp upstream and downstream of the peak summit. **(b)** Rosettes of *met1* mutants and wild-type plants from five accessions ordered by log_2_ fold change in SDC expression. White scale bar denotes 1 cm.

**Figure S25.**
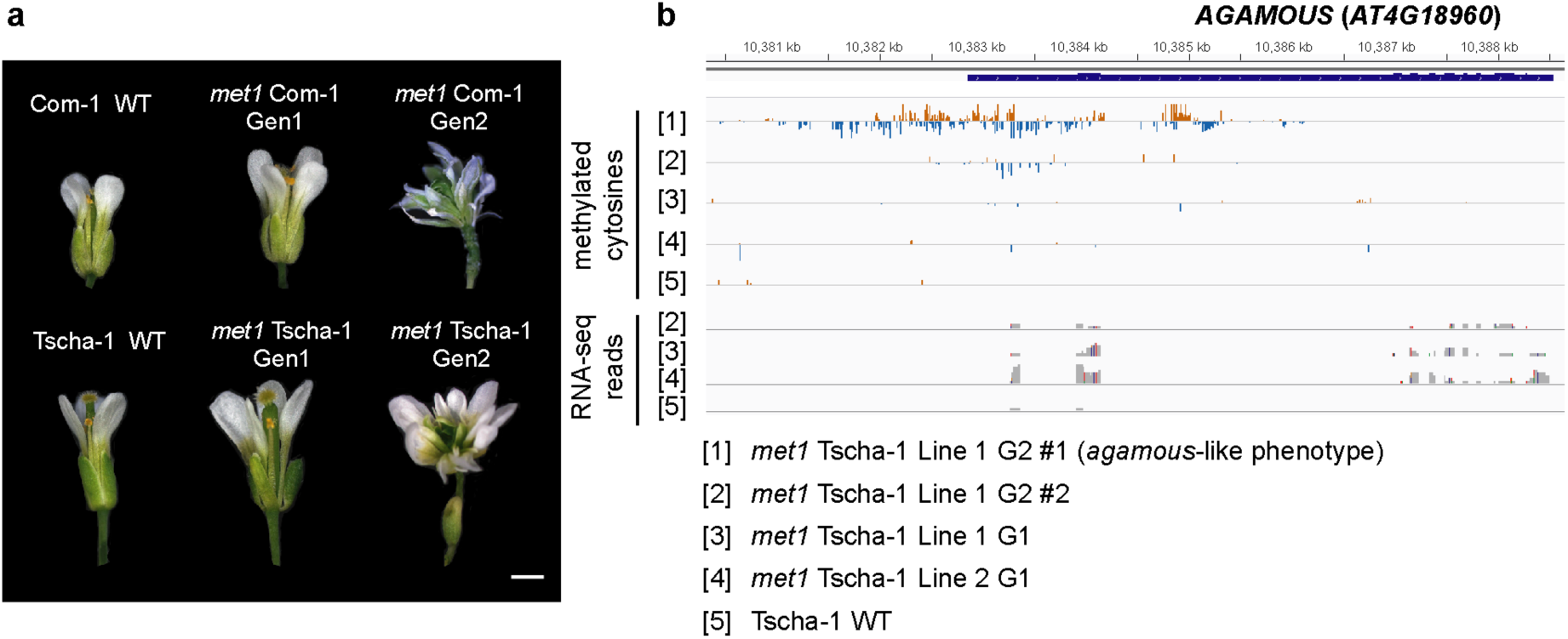
Transgenerational *ag-like* phenotypes in *met1* mutants of Com-1 and Tscha-1. A few *met1* mutant lines of the Tscha-1 and Com-1 accessions exhibited indeterminate flowers, a phenotype known to arise from genetic [74] and epigenetic inactivation [4] of the *AG* gene. BS-seq of one such line in Tscha-1 also showed an increase in methylation at this locus. **(a)** Flower phenotypes of wild-type and homozygous *met1* plants in two generations (Gen1 and Gen2). Scale bar denotes 1 mm. **(b)** Genome browser screenshot of methylated cytosines (all-contexts) and RNA-seq reads at the *AG* locus for various Tscha-1 lines.

**Figure S26.**
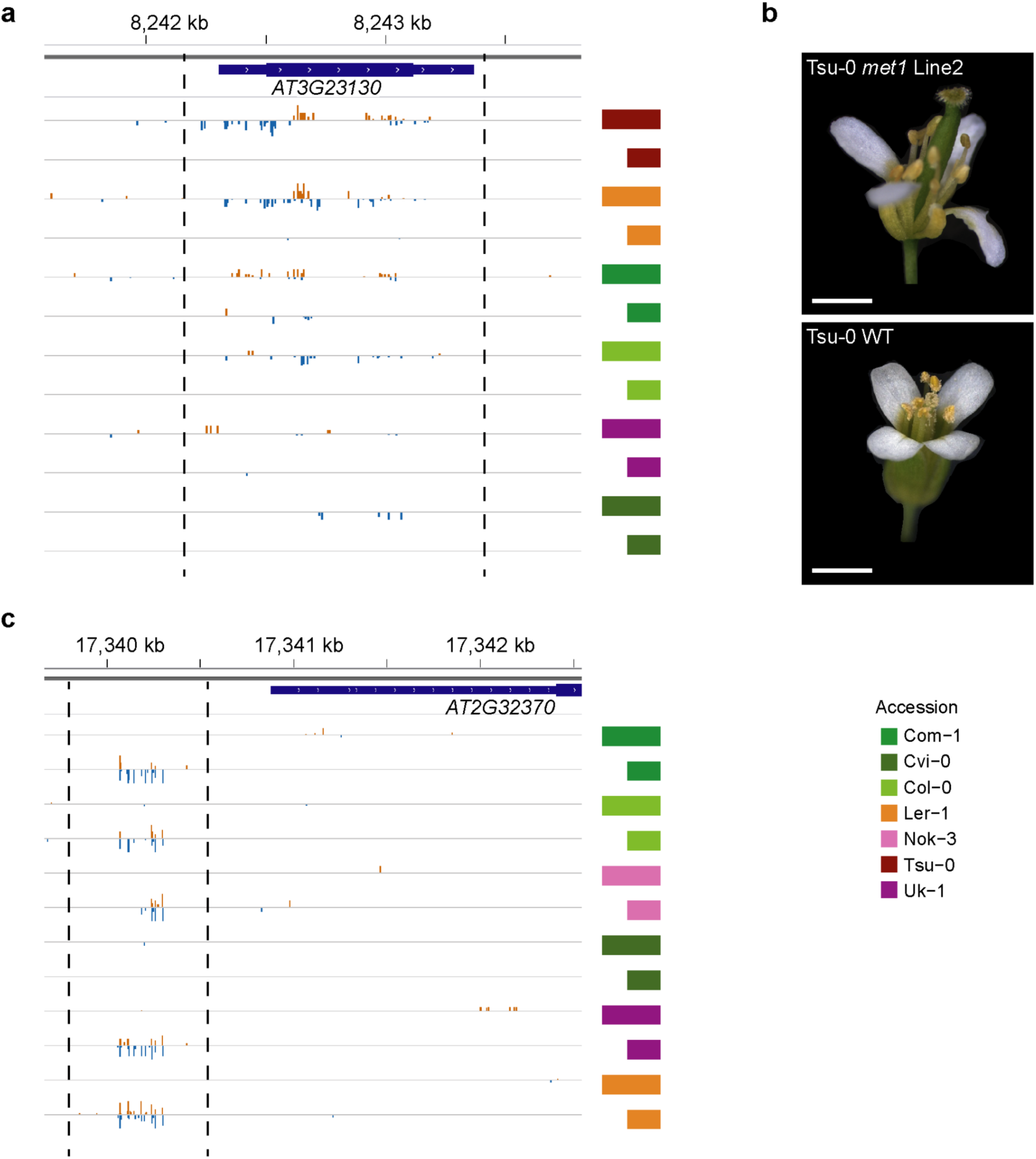
Variation in methylation levels at two epialleles in *met1* mutants and wild-type plants across a subset of accessions. **(a)** Genome browser view of methylated cytosines (all contexts) at the *SUP* (*AT3G23130*) locus. Gain of methylation in the gene body of *SUP* silences the gene and results in the formation of additional stamens [54]. **(b)** representative image of a Tsu-0 *met1* mutant flower with nine stamens and a Tsu-0 WT flower with six stamens. **(c)** Genome browser view of methylated cytosines (all contexts) at the *HDG3 (AT2G32370)* locus. Scale bars represent 1mm.

**Figure S27.**
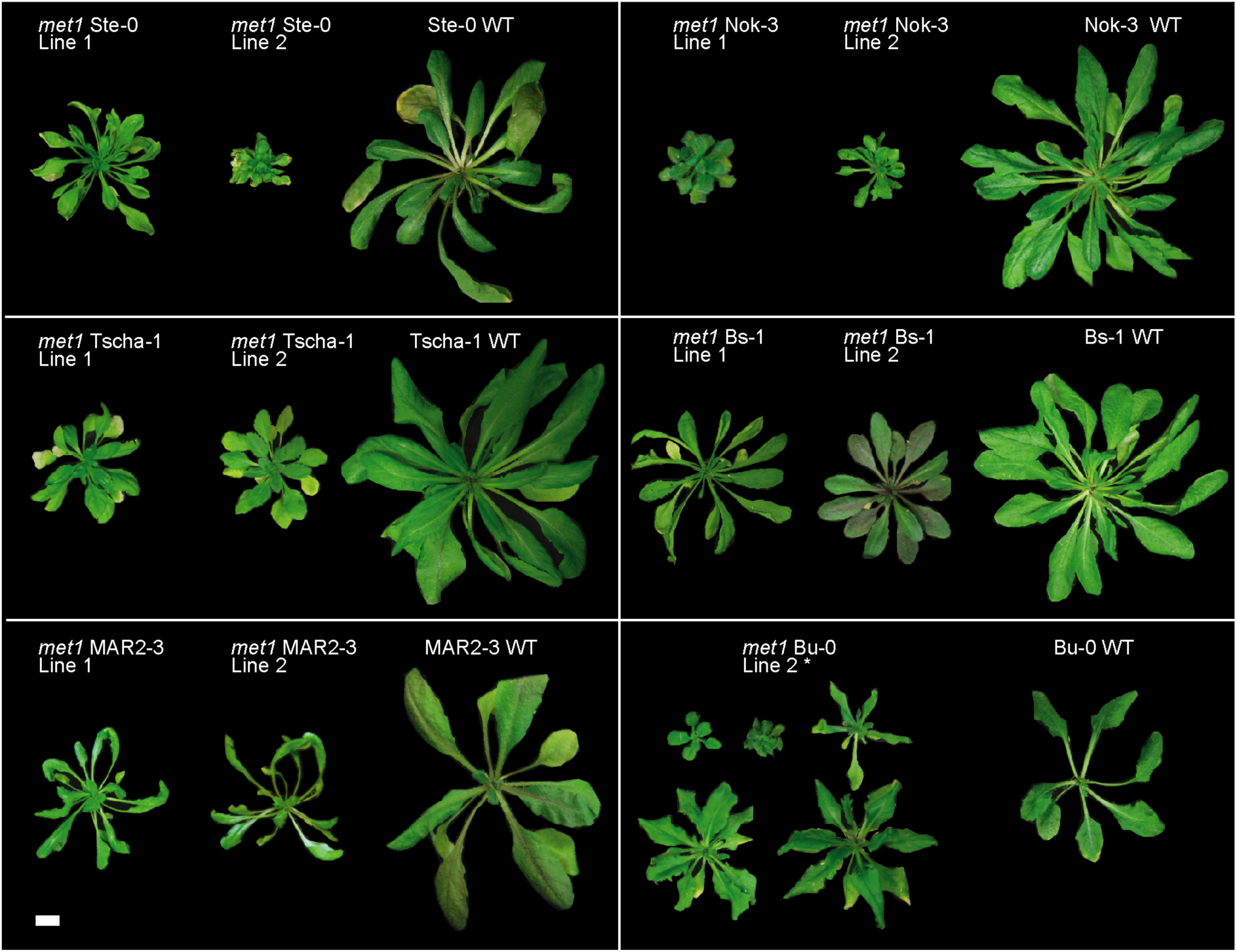
Rosettes of *met1* mutants in six accessions. Representative images of two independently derived mutant lines and a wild-type plant at six weeks after germination; scale bar denotes 1 cm. *met1* mutants of the accession Bu-0 are marked by an asterisk (*) since they were tetraploid and exhibited a wide range of phenotypes arising from different dosages of two mutant alleles and a wild-type allele.

**Figure S28.**
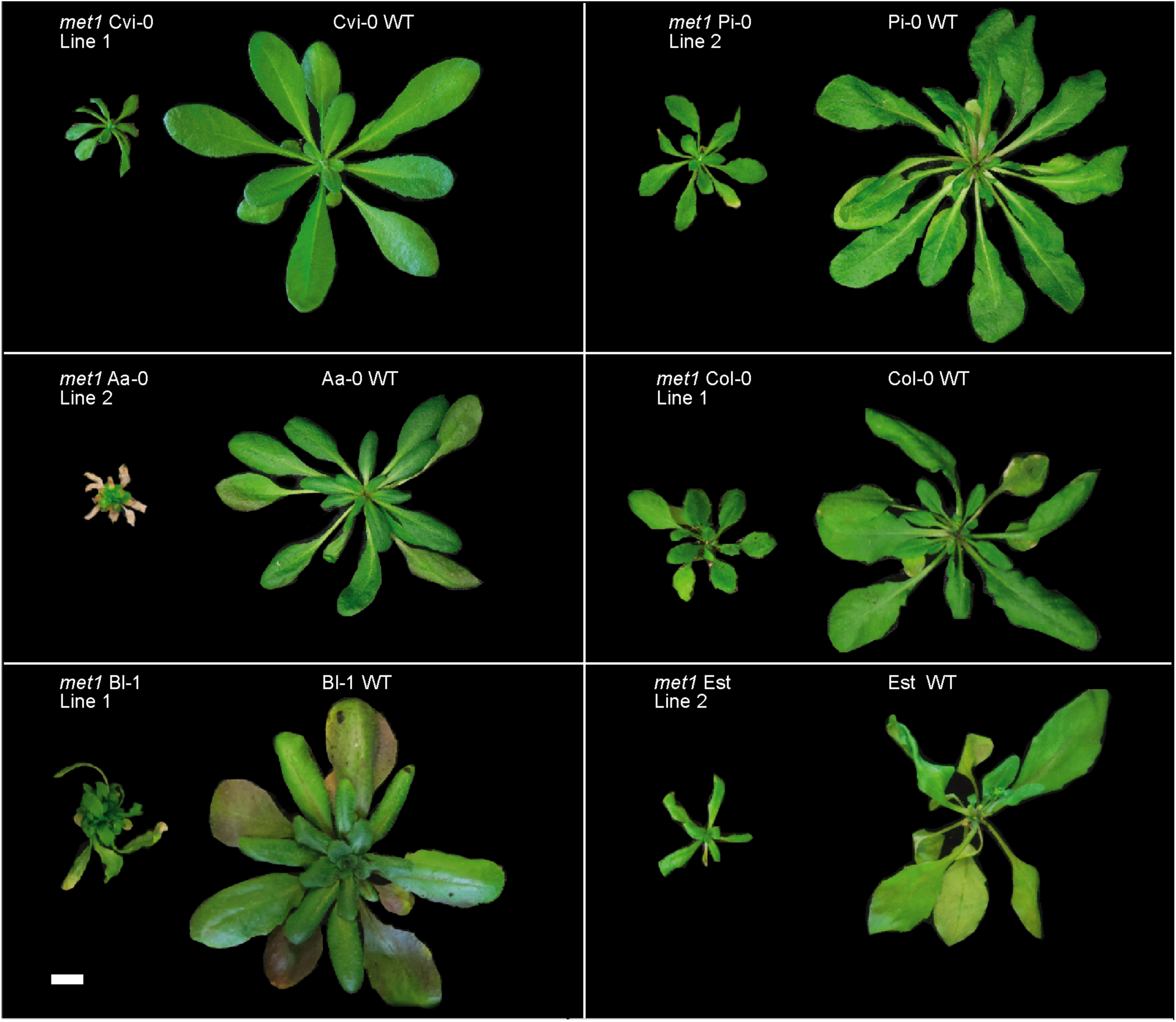
Rosettes of *met1* mutants in six accessions. Representative images of mutant and WT plants at six weeks after germination; white scale bar denotes 1 cm.

**Figure S29.**
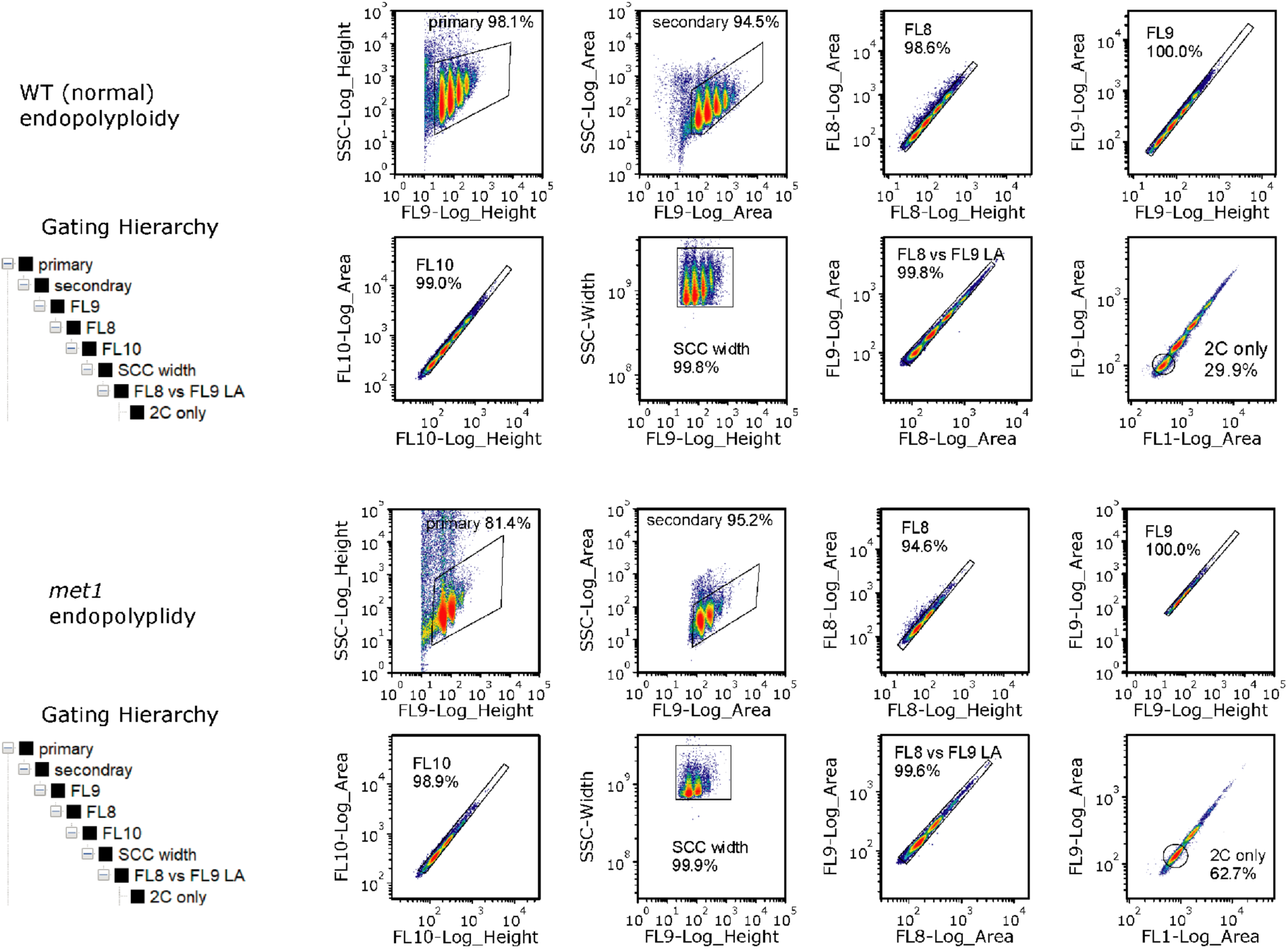
Reduced endopolyploidy in rosette leaf nuclei of *met1* mutants. An example of FACS gating profiles for WT and *met1* genotypes of the Nok-3 accession is shown.

**Figure S30.**
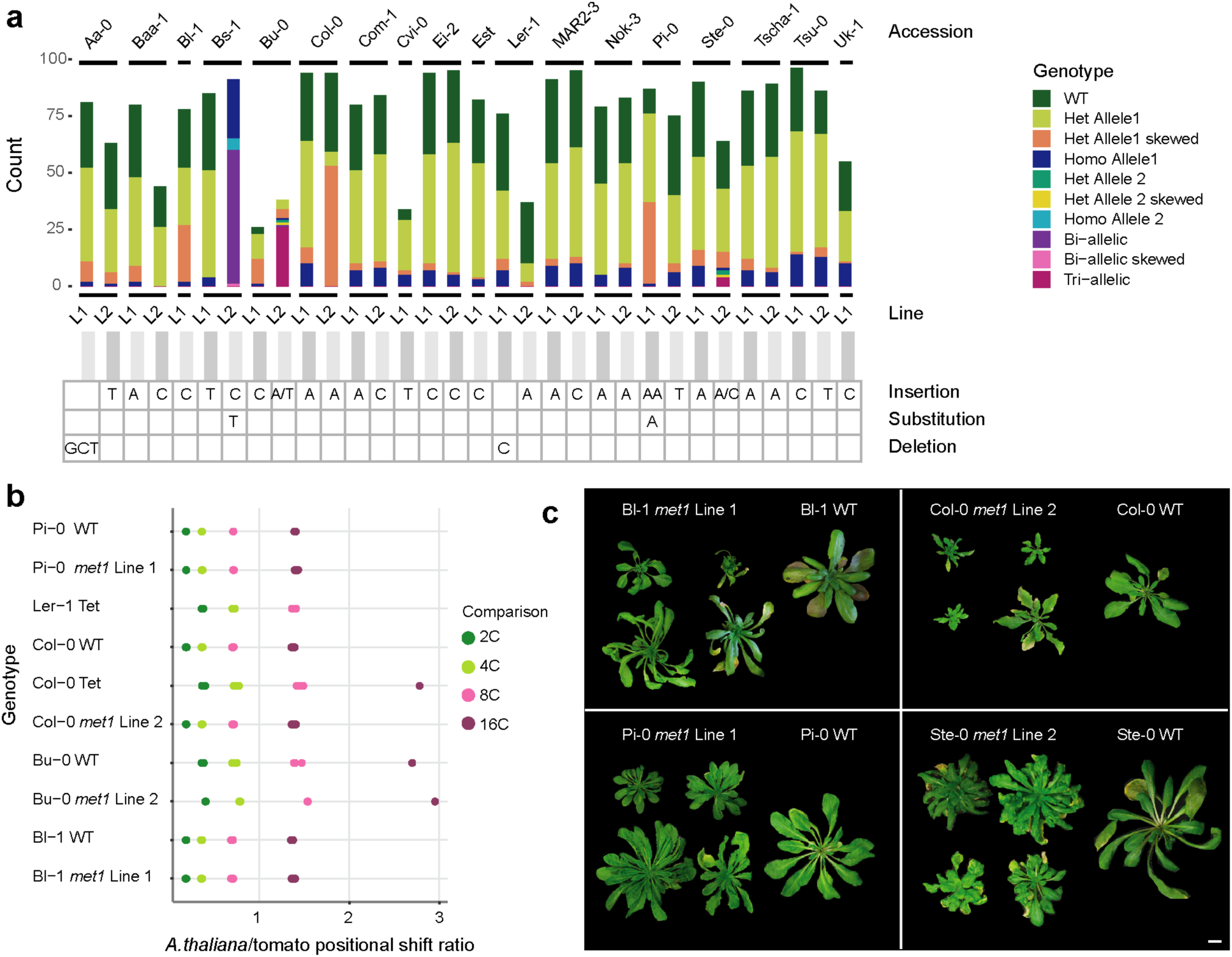
Segregation distortion in *met1* mutants and the presence of skewed heterozygous individuals. **(a)** Genotypes of segregating *met1* mutants representing sampled individuals and associated mutations for every line. **(b)** Scatter plot of endopolyploidy peak position ratios (from flow cytometry profiles) in candidate mutant lines and wild-type plants relative to the tomato internal standard. Col-0 and Ler-1 tetraploids (’Col-Tet’ and ’Ler-Tet’ respectively) were used as references for validating ploidy variation in candidate lines. **(c)** Phenotypic variation in heterozygous plants of Bl-1 Line 1, Col-0 Line 2, Pi-0 Line 1 and Ste-0 Line 2. Scale bar represents 1 cm.

## SUPPLEMENTARY DATA

### 1. Supplementary Tables

**Table S1:**

- CRISPR guide-RNA design for *MET1* knockout, primers used for PCR-based genotyping of frameshift mutations
- List of accessions used in this study
- List of mutants in each accession and identified mutations

**Table S2:**

- Numbers of DEGs, DMRs and dACRs identified across all samples
- Numbers of DEGs identified across all accession contrasts
- Numbers of unique genes identified in intersections between DEGs, DMRs and dACRs
- Intersections between DMR and dACRs with TEs and genes

**Table S3:**

- Segregation distortion results for each mutant line

**Table S4:**

- Frameshifted oligonucleotides (forward and reverse primers) used for amplicon-sequencing of multiple samples
- List of individual samples analyzed for each mutant line

### 2. Extended Methods

#### Generation of consensus ATAC-seq peaks using DiffBind

The following commands were used in R using the package “DiffBind” (where ‘dataset_Acc’ refers to the sample sheet containing details of MACS2-called peaks from individual libraries):

~~~
library(DiffBind)
dataset_Acc <- dba(sampleSheet=“./samplesheet_forDiffBind.csv”) dataset_Acc_consensus <- dba.peakset(dataset_Acc, consensus=-DBA_REPLICATE)
dataset_Acc_consensus<- dba(dataset_Acc_consensus, mask=dataset_Acc_consensus$masks$Consensus, minOverlap=1)
consensus_peaks <- dba.peakset(dataset_Acc_consensus, bRetrieve=TRUE)
dataset_Acc<- dba.count(dataset_Acc, peaks=consensus_peaks, score=DBA_SCORE_TMM_MINUS_FULL_CPM, summits=FALSE)
normCounts<- dba.peakset(dataset_Acc, bRetrieve=TRUE, DataType=DBA_DATA_FRAME)
~~~

#### Feature intersections between DEGs, DMRs and dACRs

To intersect positions of each consensus DEG set (TE-DEG/Non-TE-DEG) with DMRs, we identified the five closest DMRs in each context (CG DMRs, All-C DMRs) to each DEG. Based on their proximity to the DEG, each of these hits were classified as ’extended gene-body’ (distances within 100bp upstream of the TSS and 100bp downstream of the TTS), ’cis upstream/downstream’ (within 1.5kb upstream/downstream of the gene body) and ’trans’ if not falling within the above classes. All DEGs with DMR hits under the ’trans’ category were filtered out. DEGs with unique or multiple ’gene-body’ DMRs were classified as ’GB’, while DEGs with unique/multiple DMRs in gene-body and *cis*, or only in *cis* were classified as ’cis’. Subsequently, gene expression counts and methylation counts for the corresponding DMRs were extracted for these DEGs. Since many DEGs had multiple DMRs associated with them, we only retained DMRs which showed the highest difference in methylation level between mutant and WT for each mutant genotype, thereby aiming to represent only the strongest methylation signals that could explain gene expression differences.

For intersecting consensus DEGs with dACRs, we used a similar approach, where the closest five dACRs to each DEG were identified. After filtering out DEGs with dACRs in ’trans’, we retained all remaining DEGs (even with multiple ’gene-body’ and ’cis’ dACRs) in a single category called ’cis’. For each DEG, only dACRs with the highest difference in accessibility between mutant and WT for each mutant genotype were retained.

A more detailed explanation of the above methods is provided below :

#### 1. Intersects between DEGs and DMRs

Consensus DEGs (from all accessions) were first split as 5731 Non-TEDEGs and 1401 TEDEGs. Next, each set was intersected with positions of consensus DMRs (generated from all samples) in two contexts, CG-DMRs and All-C -DMRs, using *bedtools closest*, to find the top 5 closest DMRs to each DEG. Based on their proximity to the DEG, each of these hits were classified as gene-body (closest distance >=-100 bp and <=100 bp), cis_upstream (closest distance <=-100 bp and >-1.5 kb), cis downstream (closest distance >=100 bp and <=1.5 kb) and trans (beyond 1.5 kb upstream or downstream).

#Example of command used (to find 5 closest hits of DMRs to DEGs)

~~~
*bedtools closest -a Consensus_NonTEDEGs_coord.tab -b CG_DMRs_coord.bed -D a -k 5 > Consensus_NonTEDEGs_closestk5_CGDMRs.bed*
~~~

All DMR hits under the ’trans’ category were filtered out. Next, the number of duplicate DMR hits for each gene were counted. Each gene was further classified into the following categories: GB_unique (single DMR hit in gene body), GB_multi (multiple DMR hits in gene body), cis_upstream (single DMR hit in cis upstream), cis_downstream (single DMR hit in cis downstream), cis_multi (multiple cis DMRs), GBandcis (multiple DMRs in gene body and cis).

Genes classified as GB_unique and GB_multi were pooled together in a “GB_all” category. All remaining categories were pooled as a “multiple_cis_regulatory” category. Gene expression counts were obtained for genes in each category (for 158 samples and reduced to 73 samples to match the BS-seq dataset). Similarly, methylation levels for the corresponding DMR hit in each gene was also obtained for 73 samples.

Next, each gene was scanned to examine duplicate DMR hits and their methylation values. For each of the 73 samples, the DMR carrying the highest difference between wild-type and mutant methylation levels was retained. These genes were subsequently plotted based on their methylation differences and gene expression differences.

#### 2. Intersects between DEGs and dACRs

Each set of consensus DEGs (TE-DEGs and Non-TE-DEGs) was intersected with positions of consensus dACRs (generated from all samples) using *bedtools closest*, to find the top 5 closest dACRs to each DEG. Based on their proximity to the DEG, each of these hits were classified as gene-body (closest distance >=-100 bp and <=100 bp), cis_upstream (closest distance <=-100 bp and >-1.5 kb), cis downstream (closest distance >=100 bp and <=1.5 kb) and trans (beyond 1.5 kb upstream or downstream).

#Example of command used (to find 5 closest hits of dACRs to DEGs)

~~~
*bedtools closest -a Consensus_NonTEDEGs_coord.tab -b dACRs_coord.bed -D a -k 5 > Consensus_NonTEDEGs_closestk5_dACRs.bed*
~~~

All dACR hits under the ’trans’ category were filtered out. All other genes and corresponding dACR hits were pooled as a “multiple_cis_regulatory” category. Gene expression counts were obtained for genes in each category (for 158 ATAC-seq samples and reduced to 73 samples to match the Methylation dataset. Similarly, ATAC-seq accessibility levels (TMM) for the corresponding dACR hit in each gene was also obtained for 158 samples and reduced to 73 samples.

Next, each gene was scanned to examine duplicate dACR hits and their accessibility values. For each of the 73 samples, the dACR carrying the highest difference between wild-type and mutant accessibility levels was retained. These genes were subsequently plotted based on these accessibility differences and gene expression differences.

#### 3. Feature intersections between CG DMRs and dACRs

CG DMRs were annotated with unique IDs, and subsequently intersected with positions of dACRs using the *intersect* command of *bedtools*. Each CG DMR could have a unique or multiple intersections with dACRs. The methylation levels and accessibility levels were extracted for each of these intersected positions across 73 samples (the accessibility levels for 158 samples used for ATAC-seq analysis were averaged to match the 73 samples used for BS-seq analysis). For every CG DMR, only the dACR which showed the highest difference in accessibility between *met1* mutants and wildtypes for each accession background, was retained. Finally, the differences in CG methylation and the corresponding differences in accessibility (between mutants and wildtypes) for all CG DMRs were plotted for homozygous mutants across accessions.

#### 3. Supplementary Datasets

**Appendix 1.**

| DATA | FILE/FOLDER |
| --- | --- |
| 1. Sample metadata of ATAC-seq, Bisulfite-seq, RNA-seq and Amplicon-seq libraries uploaded on ENA | Libraries_uploaded_on_ENA/ |
| 2. Mapping efficiency, chloroplast read conversion efficiency of BS-seq libraries | Consolidated_mapping_conversionEff_results_BSseq_73libraries_modified.tab |
| 3. Mapping efficiency of ATAC-seq libraries, number of MACS2 peaks identified for each sample | ATAClib_peaks_readcounts_metadata.tab |
| 4. Mapping efficiency of gDNA libraries | gDNAlib_readcounts_metadata.tab |
| 5. Mapping efficiency of RNA-seq samples | RNAseq_mappingstats_metadata_158libraries.tab |
| 6. High count genes which were filtered out before DEG calling | HighCountgenes_forfiltering_beforeDESeq.tab |

**Appendix 2.**
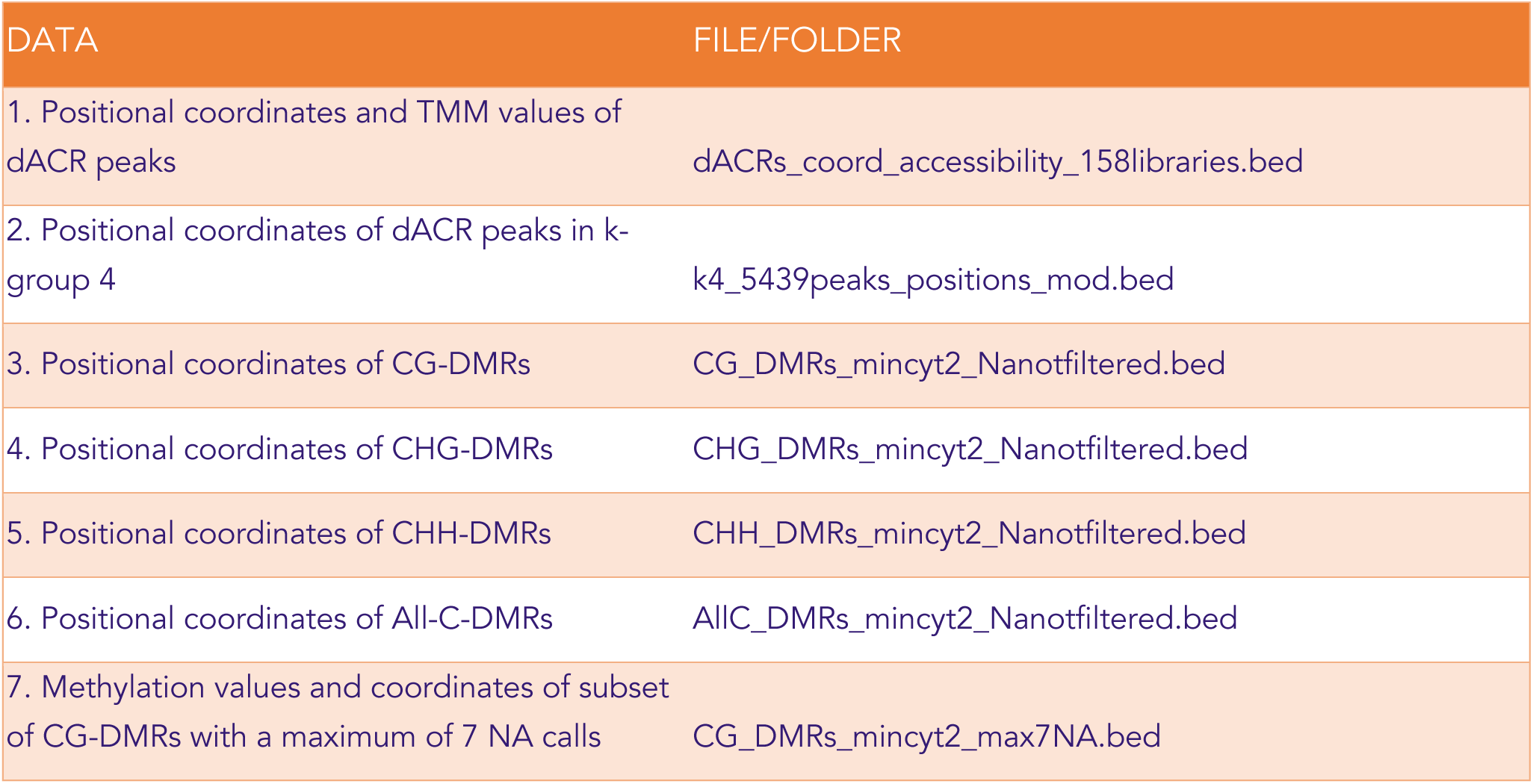
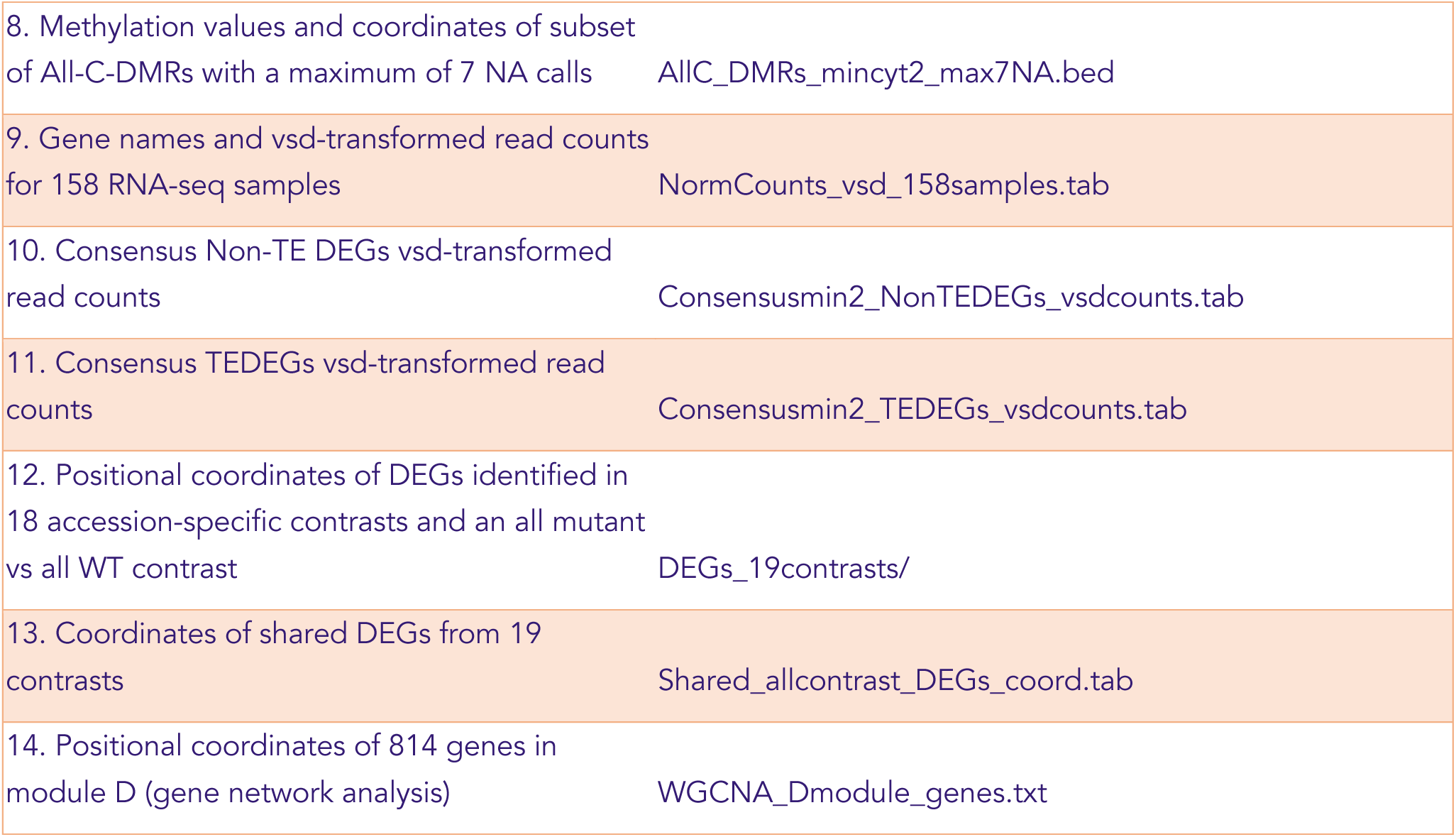

## Notes

### Competing Interest Statement

The authors have declared no competing interest.

## REFERENCES

1. Law JA, Jacobsen SE. Establishing, maintaining and modifying DNA methylation patterns in plants and animals. Nat Rev Genet. Nature Publishing Group; 2010;11:204.

2. Zhang H, Lang Z, Zhu J-K. Dynamics and function of DNA methylation in plants. Nat Rev Mol Cell Biol. 2018;19:489–506.

3. Finnegan EJ, Peacock WJ, Dennis ES. Reduced DNA methylation in Arabidopsis thaliana results in abnormal plant development. Proc Natl Acad Sci U S A. 1996;93:8449–54.

4. Jacobsen SE, Sakai H, Finnegan EJ, Cao X, Meyerowitz EM. Ectopic hypermethylation of flower-specific genes in Arabidopsis. Curr Biol. 2000;10:179–86.

5. Kankel MW, Ramsey DE, Stokes TL, Flowers SK, Haag JR, Jeddeloh JA, et al. Arabidopsis MET1 cytosine methyltransferase mutants. Genetics. 2003;163:1109–22.

6. Saze H, Mittelsten Scheid O, Paszkowski J. Maintenance of CpG methylation is essential for epigenetic inheritance during plant gametogenesis. Nat Genet. 2003;34:65–9.

7. Tariq M, Saze H, Probst AV, Lichota J, Habu Y, Paszkowski J. Erasure of CpG methylation in Arabidopsis alters patterns of histone H3 methylation in heterochromatin. Proc Natl Acad Sci U S A. 2003;100:8823–7.

8. Deleris A, Stroud H, Bernatavichute Y, Johnson E, Klein G, Schubert D, et al. Loss of the DNA Methyltransferase MET1 Induces H3K9 Hypermethylation at PcG Target Genes and Redistribution of H3K27 Trimethylation to Transposons in Arabidopsis thaliana [Internet]. PLoS Genetics. 2012. p. e1003062. Available from: http://dx.doi.org/10.1371/journal.pgen.1003062

9. Soppe WJJ, Jasencakova Z, Houben A, Kakutani T, Meister A, Huang MS, et al. DNA methylation controls histone H3 lysine 9 methylation and heterochromatin assembly in Arabidopsis. EMBO J. 2002;21:6549–59.

10. Zhong Z, Feng S, Duttke SH, Potok ME, Zhang Y, Gallego-Bartolomé J, et al. DNA methylation-linked chromatin accessibility affects genomic architecture in Arabidopsis. Proc Natl Acad Sci U S A [Internet]. 2021;118. Available from: http://dx.doi.org/10.1073/pnas.2023347118

11. Mathieu O, Reinders J, Caikovski M, Smathajitt C, Paszkowski J. Transgenerational stability of the Arabidopsis epigenome is coordinated by CG methylation. Cell. 2007;130:851–62.

12. Stroud H, Greenberg MVC, Feng S, Bernatavichute YV, Jacobsen SE. Comprehensive analysis of silencing mutants reveals complex regulation of the Arabidopsis methylome. Cell. 2013;152:352–64.

13. 1001 Genomes Consortium. 1,135 Genomes Reveal the Global Pattern of Polymorphism in Arabidopsis thaliana. Cell. 2016;166:481–91.

14. Quadrana L, Silveira AB, Mayhew GF, LeBlanc C, Martienssen RA, Jeddeloh JA, et al. The Arabidopsis thaliana mobilome and its impact at the species level [Internet]. eLife. 2016. Available from: http://dx.doi.org/10.7554/elife.15716

15. Baduel P, Leduque B, Ignace A, Gy I, Gil J Jr, Loudet O, et al. Genetic and environmental modulation of transposition shapes the evolutionary potential of Arabidopsis thaliana. Genome Biol. 2021;22:138.

16. Kawakatsu T, Huang S-SC, Jupe F, Sasaki E, Schmitz RJ, Urich MA, et al. Epigenomic Diversity in a Global Collection of Arabidopsis thaliana Accessions. Cell. 2016;166:492–505.

17. Stuart T, Eichten SR, Cahn J, Karpievitch YV, Borevitz JO, Lister R. Population scale mapping of transposable element diversity reveals links to gene regulation and epigenomic variation. Elife [Internet]. 2016;5. Available from: http://dx.doi.org/10.7554/eLife.20777

18. Dubin MJ, Zhang P, Meng D, Remigereau M-S, Osborne EJ, Paolo Casale F, et al. DNA methylation in Arabidopsis has a genetic basis and shows evidence of local adaptation. Elife. 2015;4:e05255.

19. Sasaki E, Kawakatsu T, Ecker JR, Nordborg M. Common alleles of CMT2 and NRPE1 are major determinants of CHH methylation variation in Arabidopsis thaliana. PLoS Genet. 2019;15:e1008492.

20. Shen X, De Jonge J, Forsberg SKG, Pettersson ME, Sheng Z, Hennig L, et al. Natural CMT2 variation is associated with genome-wide methylation changes and temperature seasonality. PLoS Genet. 2014;10:e1004842.

21. Sasaki E, Gunis J, Reichardt-Gomez I, Nizhynska V, Nordborg M. Conditional GWAS of non-CG transposon methylation in Arabidopsis thaliana reveals major polymorphisms in five genes [Internet]. bioRxiv. 2022 [cited 2022 Mar 20]. p. 2022.02.09.479810. Available from: https://www.biorxiv.org/content/10.1101/2022.02.09.479810v1.full

22. Shahzad Z, Moore JD, Zilberman D. Gene body methylation mediates epigenetic inheritance of plant traits. bioRxiv [Internet]. biorxiv.org; 2021; Available from: https://www.biorxiv.org/content/10.1101/2021.03.15.435374v1.abstract

23. Zhang Y, Wendte JM, Ji L, Schmitz RJ. Natural variation in DNA methylation homeostasis and the emergence of epialleles. Proc Natl Acad Sci U S A. 2020;117:4874–84.

24. Meng D, Dubin M, Zhang P, Osborne EJ, Stegle O, Clark RM, et al. Limited Contribution of DNA Methylation Variation to Expression Regulation in Arabidopsis thaliana. PLoS Genet. 2016;12:e1006141.

25. Shirai K, Sato MP, Nishi R, Seki M, Suzuki Y, Hanada K. Positive selective sweeps of epigenetic mutations regulating specialized metabolites in plants. Genome Res. 2021;31:1060–8.

26. He L, Wu W, Zinta G, Yang L, Wang D, Liu R, et al. A naturally occurring epiallele associates with leaf senescence and local climate adaptation in Arabidopsis accessions. Nat Commun. 2018;9:460.

27. Keller TE, Lasky JR, Yi SV. The multivariate association between genome wide DNA methylation and climate across the range of Arabidopsis thaliana. Mol Ecol. 2016;25:1823–37.

28. Wu R, Lucke M, Jang Y-T, Zhu W, Symeonidi E, Wang C, et al. An efficient CRISPR vector toolbox for engineering large deletions in Arabidopsis thaliana [Internet]. Plant Methods. 2018. Available from: http://dx.doi.org/10.1186/s13007-018-0330-7

29. Lippman Z, Martienssen R. The role of RNA interference in heterochromatic silencing. Nature. 2004;431:364–70.

30. Slotkin RK, Martienssen R. Transposable elements and the epigenetic regulation of the genome. Nat Rev Genet. 2007;8:272–85.

31. Oberlin S, Sarazin A, Chevalier C, Voinnet O, Marí-Ordóñez A. A genome-wide transcriptome and translatome analysis of Arabidopsis transposons identifies a unique and conserved genome expression strategy for Ty1/Copia retroelements. Genome Res. 2017;27:1549–62.

32. Zhang X, Yazaki J, Sundaresan A, Cokus S, Chan SW-L, Chen H, et al. Genome-wide high-resolution mapping and functional analysis of DNA methylation in arabidopsis. Cell. 2006;126:1189–201.

33. Lister R, O’Malley RC, Tonti-Filippini J, Gregory BD, Berry CC, Millar AH, et al. Highly integrated single-base resolution maps of the epigenome in Arabidopsis. Cell. 2008;133:523–36.

34. Marí-Ordóñez A, Marchais A, Etcheverry M, Martin A, Colot V, Voinnet O. Reconstructing de novo silencing of an active plant retrotransposon. Nat Genet. 2013;45:1029–39.

35. Mirouze M, Reinders J, Bucher E, Nishimura T, Schneeberger K, Ossowski S, et al. Selective epigenetic control of retrotransposition in Arabidopsis. Nature. 2009;461:427–30.

36. Kato M, Miura A, Bender J, Jacobsen SE, Kakutani T. Role of CG and non-CG methylation in immobilization of transposons in Arabidopsis. Curr Biol. 2003;13:421–6.

37. Le NT, Harukawa Y, Miura S, Boer D, Kawabe A, Saze H. Epigenetic regulation of spurious transcription initiation in Arabidopsis. Nat Commun. 2020;11:3224.

38. The Arabidopsis Genome Initiative. Analysis of the genome sequence of the flowering plant Arabidopsis thaliana. Nature. 2000;408:796–815.

39. Klepikova AV, Kasianov AS, Gerasimov ES, Logacheva MD, Penin AA. A high resolution map of the Arabidopsis thaliana developmental transcriptome based on RNA-seq profiling. Plant J. 2016;88:1058–70.

40. Vu TM, Nakamura M, Calarco JP, Susaki D, Lim PQ, Kinoshita T, et al. RNA-directed DNA methylation regulates parental genomic imprinting at several loci in Arabidopsis. Development. 2013;140:2953–60.

41. Hsieh T-F, Shin J, Uzawa R, Silva P, Cohen S, Bauer MJ, et al. Regulation of imprinted gene expression in Arabidopsis endosperm. Proc Natl Acad Sci U S A. 2011;108:1755–62.

42. Soppe WJ, Jacobsen SE, Alonso-Blanco C, Jackson JP, Kakutani T, Koornneef M, et al. The late flowering phenotype of fwa mutants is caused by gain-of-function epigenetic alleles of a homeodomain gene. Mol Cell. 2000;6:791–802.

43. Kinoshita Y, Saze H, Kinoshita T, Miura A, Soppe WJJ, Koornneef M, et al. Control of FWA gene silencing in Arabidopsis thaliana by SINE-related direct repeats. Plant J. Wiley Online Library; 2007;49:38–45.

44. Henderson IR, Jacobsen SE. Tandem repeats upstream of the Arabidopsis endogene SDC recruit non-CG DNA methylation and initiate siRNA spreading. Genes Dev. 2008;22:1597–606.

45. Tran RK, Henikoff JG, Zilberman D, Ditt RF, Jacobsen SE, Henikoff S. DNA methylation profiling identifies CG methylation clusters in Arabidopsis genes. Curr Biol. 2005;15:154–9.

46. Reynoso MA, Kajala K, Bajic M, West DA, Pauluzzi G, Yao AI, et al. Evolutionary flexibility in flooding response circuitry in angiosperms. Science. 2019;365:1291–5.

47. Marand AP, Chen Z, Gallavotti A, Schmitz RJ. A cis-regulatory atlas in maize at single-cell resolution. Cell. 2021;184:3041–55.e21.

48. Shu H, Wildhaber T, Siretskiy A, Gruissem W, Hennig L. Distinct modes of DNA accessibility in plant chromatin. Nat Commun. 2012;3:1281.

49. Bender J, Fink GR. Epigenetic control of an endogenous gene family is revealed by a novel blue fluorescent mutant of Arabidopsis. Cell. 1995;83:725–34.

50. Rigal M, Kevei Z, Pélissier T, Mathieu O. DNA methylation in an intron of the IBM1 histone demethylase gene stabilizes chromatin modification patterns. EMBO J. 2012;31:2981–93.

51. Stokes TL, Kunkel BN, Richards EJ. Epigenetic variation in Arabidopsis disease resistance. Genes Dev. 2002;16:171–82.

52. Williams BP, Pignatta D, Henikoff S, Gehring M. Methylation-sensitive expression of a DNA demethylase gene serves as an epigenetic rheostat. PLoS Genet. 2015;11:e1005142.

53. Lei M, Zhang H, Julian R, Tang K, Xie S, Zhu J-K. Regulatory link between DNA methylation and active demethylation in Arabidopsis. Proc Natl Acad Sci U S A. 2015;112:3553–7.

54. Jacobsen SE, Meyerowitz EM. Hypermethylated SUPERMAN epigenetic alleles in arabidopsis. Science. 1997;277:1100–3.

55. Ronemus MJ, Galbiati M, Ticknor C, Chen J, Dellaporta SL. Demethylation-induced developmental pleiotropy in Arabidopsis. Science. 1996;273:654–7.

56. FitzGerald J, Luo M, Chaudhury A, Berger F. DNA methylation causes predominant maternal controls of plant embryo growth. PLoS One. 2008;3:e2298.

57. Niederhuth CE, Bewick AJ, Ji L, Alabady MS, Kim KD, Li Q, et al. Widespread natural variation of DNA methylation within angiosperms. Genome Biol. 2016;17:194.

58. Zhou P, Lu Z, Schmitz RJ. Stable unmethylated DNA demarcates expressed genes and their cis-regulatory space in plant genomes. Proceedings of the [Internet]. National Acad Sciences; 2020; Available from: https://www.pnas.org/content/117/38/23991.short

59. Borges F, Parent J-S, van Ex F, Wolff P, Martínez G, Köhler C, et al. Transposon-derived small RNAs triggered by miR845 mediate genome dosage response in Arabidopsis. Nat Genet. 2018;50:186–92.

60. Pignatta D, Erdmann RM, Scheer E, Picard CL, Bell GW, Gehring M. Natural epigenetic polymorphisms lead to intraspecific variation in Arabidopsis gene imprinting [Internet]. eLife. 2014. Available from: http://dx.doi.org/10.7554/elife.03198

61. Pignatta D, Novitzky K, Satyaki PRV, Gehring M. A variably imprinted epiallele impacts seed development. PLoS Genet. 2018;14:e1007469.

62. Noshay JM, Marand AP, Anderson SN, Zhou P. Cis-regulatory elements within TEs can influence expression of nearby maize genes. BioRxiv [Internet]. biorxiv.org; 2020; Available from: https://www.biorxiv.org/content/10.1101/2020.05.20.107169v1.abstract

63. Pavlopoulou A, Kossida S. Plant cytosine-5 DNA methyltransferases: structure, function, and molecular evolution. Genomics. 2007;90:530–41.

64. Clough SJ, Bent AF. Floral dip: a simplified method for Agrobacterium-mediated transformation of Arabidopsis thaliana. Plant J. 1998;16:735–43.

65. LeBlanc C, Zhang F, Mendez J, Lozano Y, Chatpar K, Irish VF, et al. Increased efficiency of targeted mutagenesis by CRISPR/Cas9 in plants using heat stress. Plant J. 2018;93:377–86.

66. Wibowo A, Becker C, Durr J, Price J, Spaepen S, Hilton S, et al. Partial maintenance of organ-specific epigenetic marks during plant asexual reproduction leads to heritable phenotypic variation. Proc Natl Acad Sci U S A. 2018;115:E9145–52.

67. Krueger F, Andrews SR. Bismark: a flexible aligner and methylation caller for Bisulfite-Seq applications. Bioinformatics. 2011;27:1571–2.

68. Hüther P, Hagmann J, Nunn A, Kakoulidou I, Pisupati R, Langenberger D, et al. MethylScore, a pipeline for accurate and context-aware identification of differentially methylated regions from population-scale plant WGBS data [Internet]. bioRxiv. 2022 [cited 2022 Feb 8]. p. 2022.01.06.475031. Available from: https://www.biorxiv.org/content/10.1101/2022.01.06.475031v1

69. Yaffe H, Buxdorf K, Shapira I, Ein-Gedi S, Moyal-Ben Zvi M, Fridman E, et al. LogSpin: a simple, economical and fast method for RNA isolation from infected or healthy plants and other eukaryotic tissues. BMC Res Notes. 2012;5:45.

70. Cambiagno DA, Giudicatti AJ, Arce AL, Gagliardi D, Li L, Yuan W, et al. HASTY modulates miRNA biogenesis by linking pri-miRNA transcription and processing. Mol Plant. 2021;14:426–39.

71. Langfelder P, Horvath S. WGCNA: an R package for weighted correlation network analysis. BMC Bioinformatics. 2008;9:559.

72. Symeonidi E, Regalado J, Schwab R, Weigel D. CRISPR-finder: A high throughput and cost-effective method to identify successfully edited Arabidopsis thaliana individuals. Quantitative Plant Biology [Internet]. Cambridge University Press; 2021 [cited 2022 Jul 14];2. Available from: https://www.cambridge.org/core/journals/quantitative-plant-biology/article/crisprfinder-a-high-throughput-and-costeffective-method-to-identify-successfully-edited-arabidopsis-thaliana-individuals/47277DE29C86370619B0A3842C70666D

73. Loureiro J, Rodriguez E, Dolezel J, Santos C. Two new nuclear isolation buffers for plant DNA flow cytometry: a test with 37 species. Ann Bot. 2007;100:875–88.

74. Yanofsky MF, Ma H, Bowman JL, Drews GN, Feldmann KA, Meyerowitz EM. The protein encoded by the Arabidopsis homeotic gene agamous resembles transcription factors. Nature. 1990;346:35–9.

